# SRSF1 coordinates transcription and alternative splicing to modulate p53 and apoptosis

**DOI:** 10.64898/2026.10.16.759576

**Authors:** Sean Paz, Elena Bisotto, Massimo Caputi

## Abstract

Serine/arginine-rich splicing factor 1 (SRSF1) regulates alternative splicing and acts as a proto-oncogene, but its transcriptional functions remain poorly characterized. We combined time-course RNA sequencing and experimental validation to characterize SRSF1-dependent transcription and alternative splicing. We validated 66 of 142 splicing events, including 43 unannotated in GENCODE and 15 containing previously unannotated exons. Thirty-five events showed concordant regulation across HEK293, MCF-7, and PC-3 cells, including 17 in 16 genes associated with p53 functions. SRSF1 overexpression reduced total p53, increased basal Ser15 phosphorylation, and increased apoptosis under basal and cisplatin-treated conditions but not following doxorubicin. SRSF1 increased transcription of 28 genes, including 18 consistently regulated across all three cell lines. Mechanistic studies linked transcriptional activation to P-TEFb mobilization from the 7SK snRNP. Analyses of 33 TCGA cancer cohorts and 47 GTEx normal-tissue cohorts identified widespread positive associations between SRSF1 expression and these genes. Additional analyses identified associations with breast cancer subtypes, thyroid differentiation, and renal cancer clinical outcomes. These findings establish concurrent SRSF1-dependent transcriptional and splicing programs associated with cellular signaling, genome maintenance, p53 regulation, and cancer biology.

## Introduction

Serine/arginine-rich splicing factor 1 (SRSF1) is a multifunctional RNA-binding protein that has been characterized as a proto-oncogene and whose expression is frequently elevated in solid and hematologic malignancies (1–3). SRSF1 is the prototypical member of the serine/arginine-rich (SR) protein family and one of its most extensively studied proteins. SRSF1 binds exonic and intronic sequence elements and promotes early spliceosome assembly by facilitating recruitment of core spliceosomal components to nascent pre-mRNA. Although SRSF1 is classically viewed as a positive regulator of exon inclusion through binding to exonic splicing enhancers, its activity is context dependent. Its position relative to splice sites, local RNA structure, splice-site strength, and interactions with neighboring RNA-binding proteins can also promote exon skipping when SRSF1 associates with intronic or repressive regulatory elements (1,4,5). Beyond pre-mRNA splicing, SRSF1 functions in mRNA export, RNA stability, translation, nonsense-mediated decay, and transcription-associated RNA processing (1,6).

Increased SRSF1 expression has been associated with breast, lung, colorectal, liver, renal, pancreatic, prostate, and brain tumors, as well as hematologic malignancies (2,3). SRSF1 overexpression can result from gene amplification, MYC-dependent transcriptional activation, or disruption of autoregulatory and cross-regulatory mechanisms that normally constrain SR-protein expression (7–9). Much of the evidence linking SRSF1 to tumorigenesis derives from its effects on alternative splicing. SRSF1 regulates the splicing of BIN1, BIM, BCL-X, and MCL1, generating isoforms associated with increased cell survival, resistance to apoptosis, and transformation (2,10–12). In mammary epithelial cells, increased SRSF1 promotes proliferation, transformation, and altered acinar morphogenesis (2). In glioma, SRSF1 promotes oncogenic MYO1B splice switching and stabilizes the long noncoding RNA NEAT1, affecting cell-cycle progression (13,14). In pancreatic, lung, and breast cancer models, SRSF1-dependent regulation of AURKA and PTPMT1 splicing has been associated with tumor progression and therapeutic resistance (15–17). These studies establish alternative splicing as a major mechanism through which increased SRSF1 expression alters cancer-associated cellular programs.

Several SR proteins also regulate transcription. Following transcription initiation, RNA polymerase II (Pol II) frequently pauses near the promoter through the activities of NELF (Negative Elongation Factor) and DSIF (DRB Sensitivity-Inducing Factor). Productive elongation requires positive transcription elongation factor b (P-TEFb)-dependent phosphorylation of NELF, DSIF, and Ser2 residues in the Pol II C-terminal domain (18,19). P-TEFb activity is regulated in part by the 7SK small nuclear ribonucleoprotein (7SK snRNP), which contains the noncoding 7SK snRNA and sequesters P-TEFb in an inactive state; release of P-TEFb activates its kinase activity and promotes productive elongation (18,19). SRSF2 associates with promoter-proximal nascent RNA and facilitates P-TEFb release from 7SK snRNP, thereby promoting release of paused Pol II into productive elongation (19,20). We previously showed that SRSF1 binds overlapping sequence elements within HIV-1 TAR and 7SK snRNA, competes with Tat for TAR binding, and enhances basal HIV-1 transcription, supporting a role for SRSF1 in 7SK-dependent regulation of P-TEFb (21).

Although previous studies established the contribution of SRSF1-dependent alternative splicing to cancer-associated cellular functions, the broader consequences of increased SRSF1 expression for endogenous transcription and alternative splicing remain poorly characterized. How SRSF1 affects genes controlling genome maintenance, p53 regulation, and responses to DNA damage remains unclear. Moreover, the relationship between SRSF1 expression and its transcriptionally regulated genes across human tumors and normal tissues has not been systematically characterized. We addressed these questions using time-course RNA-seq, experimental validation of alternative-splicing events, RT-qPCR, nascent RNA and RNA-stability assays, and analyses of 7SK snRNP-dependent transcriptional regulation. We examined the conservation of these changes across HEK293, MCF-7, and PC-3 cells and evaluated p53 regulation and apoptosis under basal conditions and following cisplatin or doxorubicin treatment. We further analyzed SRSF1-associated gene expression across The Cancer Genome Atlas (TCGA) tumor cohorts and Genotype-Tissue Expression (GTEx) normal-tissue cohorts, including associations with cancer subtype, differentiation, and clinical outcomes. Our findings identify distinct, concurrent SRSF1-dependent transcriptional and alternative-splicing programs linked to P-TEFb regulation, genome maintenance, p53 function, and cellular stress responses, with corresponding gene-expression associations across human tissues and cancers.

## Materials and Methods

### Cell lines and culture

HEK293 (ATCC CRL-1573; RRID:CVCL_0045), MCF-7 (ATCC HTB-22; RRID:CVCL_0031), and PC-3 (ATCC CRL-1435; RRID:CVCL_0035) cells were obtained directly from ATCC and maintained according to ATCC recommendations. HEK293 and MCF-7 cells are female-derived, whereas PC-3 cells are male-derived. Cell sex was not used as an experimental variable because the study compared responses within each cell line rather than male- and female-derived samples of the same cellular model. Cell identity was based on direct acquisition from ATCC and the corresponding catalog identifiers. No additional short tandem repeat authentication was performed after receipt. Cells were tested for mycoplasma contamination using the MycoAlert® PLUS Mycoplasma Detection Kit (Lonza) according to the manufacturer’s instructions and were confirmed to be mycoplasma negative. Mycoplasma testing was performed before each assay and every 4 weeks during culture. All experiments were performed using cells maintained for 5-20 passages after thawing.

### RNA sequencing and bioinformatic analysis

Total RNA was isolated from HEK293 cells 15 and 48 h after transfection with the control pCMV-Sport6 vector expressing the T7 tag (pControl) or pCMV-T7-SRSF1 (pSRSF1-T7), which we previously described (21). Samples with RNA integrity number ≥7 underwent rRNA depletion with Ribo-Zero (Illumina) and library preparation with the TruSeq RNA Library Prep Kit (Illumina). Three independent biological replicates were analyzed for each condition. Libraries were sequenced on an Illumina HiSeq 2500, generating approximately 40 million 80-bp paired-end reads per sample (Supplementary Figure S1).

Reads were trimmed with Trimmomatic v0.40 (RRID:SCR_011848) and aligned to GRCh38 with HISAT2 v2.2.2 (RRID:SCR_015530). Differential gene expression was analyzed with Cuffdiff v2.2.1 (RRID:SCR_001647). Alternative splicing was analyzed with rMATS-turbo v4.3.0 (RRID:SCR_023485) using GENCODE release 49 (RRID:SCR_014966) (Supplementary Figure S1). Genes were selected when they showed |log2 fold change| ≥1.5 at 15 h and |log2 fold change| ≥2.0 with *P* ≤0.05 at 48 h, with concordant regulation at both time points. The 15-h dataset served solely as an early directional-response filter and was not treated as an independently significant DEG set. Alternative-splicing events (ASEs) required |ΔΨ| ≥0.10 with FDR ≤0.05 at 15 h and |ΔΨ| ≥0.25 with FDR ≤0.05 at 48 h, with concordant regulation at both time points. NCBI Gene annotations were used to classify gene biotypes. Pathway analysis identified 171 ASEs within selected functional categories; consolidation of closely mapping events yielded 142 nonredundant ASEs in 136 genes for experimental validation.

### Cell transfection and expression vectors

HEK293 cells were transfected with 0.3 µg pControl or pSRSF1-T7 using Lipofectamine 2000 (Invitrogen). MCF-7 and PC-3 cells were transfected with 0.5 µg pCMV-RLuc-IRES-EGFP or pCMV-SRSF1-IRES-EGFP. These vectors were generated in the pCMV-Sport6 backbone by inserting the HCV IRES upstream of EGFP and downstream of SRSF1 or Renilla luciferase (RLuc).

To study the role of the 7SK snRNP in SRSF1-regulated gene expression, HEK293 cells were cotransfected with 0.3 µg pControl or pSRSF1-T7 and either two phosphorothioate-modified antisense DNA oligonucleotides targeting 7SK RNA (AS94, 5′-CTAGCCAGCCAGATCAGCCG-3′; AS99, 5′-CCTTGAGAGCTTGTTTGGAGG-3′; Sigma; 100 nM each) or a control phosphorothioate-modified oligonucleotide (200 nM). For HEXIM1 or HEXIM2 overexpression, cells were cotransfected with 0.3 µg of either the HEXIM1 or HEXIM2 expression plasmid and 0.3 µg pControl or pSRSF1-T7. Transfections were performed with 1 µL Lipofectamine 2000 (Invitrogen) per reaction. HEXIM1 and HEXIM2 expression vectors were generated by cloning each open reading frame (ORF; GenScript) into the pCMV-Sport6 backbone.

Because these plasmids were generated in our laboratory, no RRIDs are available. GFP-positive cells were isolated 48 h after transfection using a BD FACSAria Fusion (BD Biosciences).

### RNA-stability assays

For RNA-stability assays, HEK293 cells were transfected with pControl or pSRSF1-T7. At 24 h, replicate cultures were harvested as the 0-h samples or treated with actinomycin D (10 µg/mL; Sigma-Aldrich) or DMSO as a control. RNA was isolated at the indicated times and analyzed by RT-qPCR. RNA levels were expressed relative to the corresponding 0-h value.

### DNA-damage treatments and apoptosis measurements

For DNA-damage assays, HEK293 cells were transfected as above and, 24 h later, treated with doxorubicin (0.5 µM; Sigma-Aldrich), cisplatin (10 µM; Sigma-Aldrich), or vehicle. Cells were harvested 6 or 24 h later for western blotting.

Apoptosis was measured with the RealTime-Glo Annexin V Apoptosis Assay (Promega). Luminescence was recorded at the indicated times using a CLARIOstar microplate reader (BMG LABTECH) and corrected using no-cell wells containing the corresponding drug or vehicle. For graphical presentation, apoptosis was expressed relative to the corresponding mock-transfected control at each time point. Statistical analyses used background-corrected raw relative luminescence unit (RLU) values.

### Western blotting

Cells were lysed in RIPA buffer supplemented with protease inhibitors, NaF, and Na VO . Total protein concentration was measured using the Pierce™ BCA Protein Assay Kit (Thermo Scientific) according to the manufacturer’s instructions. Proteins were resolved by 10% SDS-PAGE and transferred to 0.2-µm nitrocellulose membranes.

Membranes were probed with mouse monoclonal anti-SRSF1/ASF-1 clone AK96 (Sigma-Aldrich, Cat. MABE163, RRID: AB_11205367), anti-T7 (Fortis Life Sciences, Cat. A190-116, RRID: AB_670415), anti-GAPDH (Cell Signaling Technology, Cat. 2118; RRID:AB_561053), anti-p53 (Cell Signaling Technology, Cat. 9282; RRID:AB_331476), and anti-phospho-p53 Ser15 (Cell Signaling Technology, Cat. 9284; RRID:AB_331464), anti-FLAG M2 (Sigma-Aldrich, F3165; RRID:AB_259529), anti-CDK9 (Cell Signaling Technology, Cat. 2316, RRID:AB_2291505), anti-Cyclin T1 (Santa Cruz Biotechnology, Cat. sc-271348, RRID:AB_10608086), anti-HEXIM1 (Fortis Life Sciences, Cat. A303-111, RRID:AB_10893099). The RRIDs for the three Cell Signaling Technology antibodies are established in the Antibody Registry. Signals were detected with SuperSignal West Femto substrate (Thermo Fisher Scientific) and imaged using a LI-COR Odyssey FC system. Band intensity was quantified using LI-COR Image Studio. For total p53 analysis, lysate loading was adjusted to equivalent amounts of total protein. For phospho-p53 Ser15 analysis, lysate loading was adjusted to equivalent amounts of total p53. Western blot source images are provided in the Source Western Blot Images dataset.

### Co-immunoprecipitation

HEK293 cells were seeded at 3.5 × 10 cells per 10-cm dish and cotransfected the following day with 10 µg 3×FLAG-LARP7 (p3XFL-LARP7) and 5 µg pSRSF1-T7 or pControl using 15 µL Lipofectamine 2000 (Invitrogen). After 48 h, cells were washed and collected in ice-cold PBS, pelleted at 1,000 × *g* for 5 min, and lysed in 200 µL buffer containing 50 mM Tris-HCl (pH 7.5), 100 mM NaCl, 1% NP-40, 5% glycerol, 2 mM EDTA, 50 mM NaF, 1 mM Na VO, 1 mM PMSF, and 1× EDTA-free protease inhibitor cocktail (Roche). Lysates were incubated for 30 min at 4 °C with rotation, sonicated using a Bioruptor Pico (Diagenode) for six cycles of 30 s on/30 s off, and centrifuged at 10,000 × *g* for 10 min at 4 °C. Protein concentrations in clarified lysates were determined by BCA assay (Thermo Fisher Scientific).

For immunoprecipitation, 25 µL Dynabeads Protein G (Invitrogen) were washed with PBS containing 0.05% Tween-20 (PBS-T) and incubated with 3 µg anti-FLAG M2 antibody (Sigma-Aldrich, F3165; RRID:AB_259529) for 1 h at room temperature with rotation. A parallel negative-control immunoprecipitation used purified rabbit IgG (Bethyl Laboratories, P120-301; RRID:AB_479831). After three washes with PBS-T, antibody-coupled beads were incubated with 500 µg clarified lysate in 125 µL lysis buffer for 2 h at 4 °C with rotation; 5% of each lysate was reserved as input. Beads were washed five times with 500 µL wash buffer containing 50 mM Tris-HCl (pH 7.5), 250 mM NaCl, 1% NP-40, 1 mM PMSF, and 1× EDTA-free protease inhibitor cocktail. Bound proteins were eluted in 100 µL 1× Laemmli sample buffer at 100 °C for 5 min and analyzed together with input samples by SDS-PAGE and western blotting. The p3XFL-LARP7 plasmid was generated by cloning the LARP7 ORF (GenScript) into pcDNA3.1-3XFLAG in frame with the N-terminal 3×FLAG tag.

### RT-qPCR and RT-PCR analysis

RNA was extracted with TRIzol (Invitrogen), treated with TURBO DNase (Invitrogen), and reverse transcribed using random hexamers and SuperScript II (Invitrogen). qPCR was performed with Green-2-Go SYBR Green qPCR Master Mix (BioBasic) on an Agilent AriaMx system using 95°C for 10 min followed by 40 cycles of 95°C for 30 s and 62°C for 1 min. Relative RNA levels were calculated by the 2^−ΔΔCt method using RPL13A as the endogenous control. Each biological replicate was analyzed in duplicate technical reactions. Complete datasets, including no-RT controls, are provided in the Source Data.

Endpoint RT-PCR was performed with Phire Green Hot Start II DNA Polymerase using 35 amplification cycles. Products were resolved on 1.5-2.0% agarose gels and visualized using a LI-COR Odyssey FC system. Band intensity was quantified using LI-COR Image Studio. Primer sequences are listed in Appendix Table S1. ASEs were considered validated when independent biological experiments reproduced the change in isoform ratio and/or direction. Source images are provided in the RT-PCR Source Images and Source Data.

### BrU nascent RNA labeling assay

HEK293 cells transfected with pControl or pSRSF1-T7 were labeled 12 h later with 2 mM 5-bromouridine (BrU; Sigma-Aldrich) for 3 h. BrU-labeled RNA was immunoprecipitated using anti-BrdU antibody (BD Biosciences, **#**347580 RRID:AB_10015219) coupled to streptavidin magnetic beads (Promega) through biotinylated protein A and rabbit anti-mouse IgG (ThermoFisher Cat. 31188, RRID:AB_228419). Beads were washed with PBS containing 0.2% Triton X-100, and bound RNA was eluted at 95°C and analyzed by RT-qPCR. Samples processed without BrU served as negative controls.

### Functional enrichment and pathway analysis

Functional enrichment analysis used Gene Ontology Biological Process annotations and Reactome pathways with Bioconductor clusterProfiler v4.8.2 (RRID:SCR_016884) and g:Profiler g:GOSt release 2024_01 (RRID:SCR_006809). All protein-coding human genes were used as the background gene set. Statistical significance was assessed using Benjamini-Hochberg correction, with FDR <0.05 considered significant. The RRIDs for clusterProfiler and g:Profiler are registered in SciCrunch.

### Construction of the p53 regulatory interaction network

The p53-centered network incorporated STRING v11 (RRID:SCR_005223) interactions with combined confidence scores ≥0.70 and experimentally supported regulatory relationships from the literature (Appendix Table S4). Reactome, Gene Ontology, and literature-curated information provided functional annotations. Interactions were classified as activating, inhibitory, or functional associations according to their documented effects on p53 stability, activation, or transcriptional activity. Indirect or context-dependent relationships were classified as functional associations.

### TCGA and GTEx expression analyses

We analyzed SRSF1-associated target expression separately in 33 TCGA tumor cohorts and 47 GTEx normal-tissue cohorts using the 28 experimentally validated transcriptional targets and the nested 18-gene subset validated across HEK293, MCF-7, and PC-3 cells. Gene-set expression scores were standardized within each cohort. We assessed associations using continuous standardized log (SRSF1 + 1) expression across all eligible samples and comparisons between SRSF1-high and SRSF1-low samples, defined by the upper and lower expression quartiles, respectively. All analyses were performed within individual cohorts and adjusted for sample-composition covariates and log (junction-library size + 1). We calculated covariate-adjusted partial correlations and used two-sided regression tests with heteroscedasticity-consistent HC3 standard errors. Benjamini–Hochberg correction was applied within the prespecified gene-set test families, retaining the original 400 planned tests and treating TCGA and GTEx separately within each model/test family.

Individual-gene analyses used log (TPM + 1)-transformed target expression and continuous SRSF1 expression with the same covariate adjustments. Benjamini–Hochberg correction included all 2,240 planned comparisons (28 genes × 80 cohorts). Figure 6B displays the 13 TCGA and 24 GTEx cohorts in which either gene-set score met the FDR-adjusted *P* < 0.000005 criterion; individual-gene adjusted *P* values retained the original 2,240-test correction. Breast cancer analyses were performed separately in luminal A, basal-like, luminal B, and HER2-enriched tumors, additionally adjusting for age and grouped stage. Thyroid carcinoma models examined associations between target expression and the thyroid differentiation score, adjusting for age, sex, grouped stage, sample composition, and junction-library size, with or without additional adjustment for SRSF1 expression. Subtype and differentiation analyses used HC3 standard errors and Benjamini–Hochberg correction within their prespecified test families. We also compared SRSF1 expression across breast cancer molecular subtypes, adjusting for age, grouped stage, sample composition, and junction-library size. Pairwise comparisons between the four subtypes were corrected for multiple testing using the Benjamini–Hochberg procedure across all six comparisons.

We evaluated overall survival and progression-free interval (PFI) using complete-case Cox proportional-hazards models. Hazard ratios represent a one-standard-deviation increase in log (TPM + 1)-transformed target expression. Clear-cell renal carcinoma models adjusted for age and sex and stratified baseline hazards by stage I–II versus III–IV to address a stage-related proportional-hazards violation. Overall-survival analyses included 525 patients and 172 events, while PFI analyses included 523 patients and 157 events. Benjamini–Hochberg correction of the 56 rerun KIRC tests retained the original denominator of 168 planned individual-gene survival tests for each model specification. Pancreatic adenocarcinoma PFI models adjusted for age, sex, and grouped stage (174 patients; 104 events). We classified genes as low abundance when fewer than 10% of samples had expression ≥1 TPM. Clinical analyses were exploratory, with FDR-adjusted *P* < 0.05 considered significant.

### Statistical analysis

Quantitative data are presented as mean ± SEM unless otherwise indicated. RT-qPCR measurements comprised duplicate technical reactions from three independent biological replicates. We compared total p53 and phospho-p53 Ser15 levels between matched pControl- and pSRSF1-transfected cells using paired, two-tailed Student’s *t*-tests (n = 3 biological replicates). We did not perform formal normality tests because of the small sample size; paired *t*-tests assume approximately normally distributed within-pair differences.

Apoptosis measurements under vehicle, doxorubicin, and cisplatin treatment were analyzed separately using two-way repeated-measures ANOVA, with transfection condition (mock, pControl, or pSRSF1), time (0, 6, 12, 30, and 48 h), and their interaction included in the model. Time was treated as the repeated-measures factor. Šídák-adjusted comparisons evaluated differences between pSRSF1-transfected cells and mock- or pControl-transfected cells at each time point.

To assess how 7SK RNA depletion and HEXIM1/HEXIM2 overexpression affected SRSF1-dependent transcriptional activation, we calculated SRSF1-induced log RT-qPCR fold changes for each gene and experimental condition. For each of three matched biological replicates, we subtracted the reference SRSF1 response (pControl versus pSRSF1 alone) from the corresponding response under AS_CTL, AS_7SK, HEXIM1, or HEXIM2 conditions. Negative interaction contrasts indicated attenuation of SRSF1-dependent activation. We tested these contrasts against zero using two-sided, one-sample Student’s *t*-tests (n = 3, df = 2), equivalent to paired comparisons of the SRSF1 responses. We report unadjusted *P* values for these experiments.

Experimental statistical analyses were performed using GraphPad Prism v11.1.0 (RRID: SCR_002798). Unless otherwise indicated, *P* < 0.05 was considered statistically significant. Functional-enrichment analyses used Benjamini–Hochberg correction, with FDR < 0.05 considered significant. Statistical procedures for TCGA and GTEx analyses are described

## Results

### Time-course RNA-seq identifies SRSF1-dependent changes in gene expression and alternative splicing

We performed time-course RNA sequencing in HEK293 cells transiently expressing T7-tagged SRSF1 to characterize SRSF1-dependent changes in gene expression and alternative splicing. We used HEK293 cells as the discovery system because of their high transfection efficiency and functional transcriptional and RNA-processing pathways, including 7SK/P-TEFb regulation. T7-SRSF1 became detectable by 12 h and increased through 48 h (Fig. 1A). We analyzed cells at 15 and 48 h after transfection to distinguish early changes that emerged shortly after SRSF1 accumulation from changes maintained or increased at the later time point (Supplementary Figure S1). The 15 h time point also reduced the contribution of secondary effects that could develop through SRSF1-dependent regulation of other transcriptional or splicing factors, whereas the 48 h time point identified changes that persisted as SRSF1 expression increased. We prioritized gene-expression and alternative-splicing changes that occurred in the same direction at both time points.

**Figure 1.**
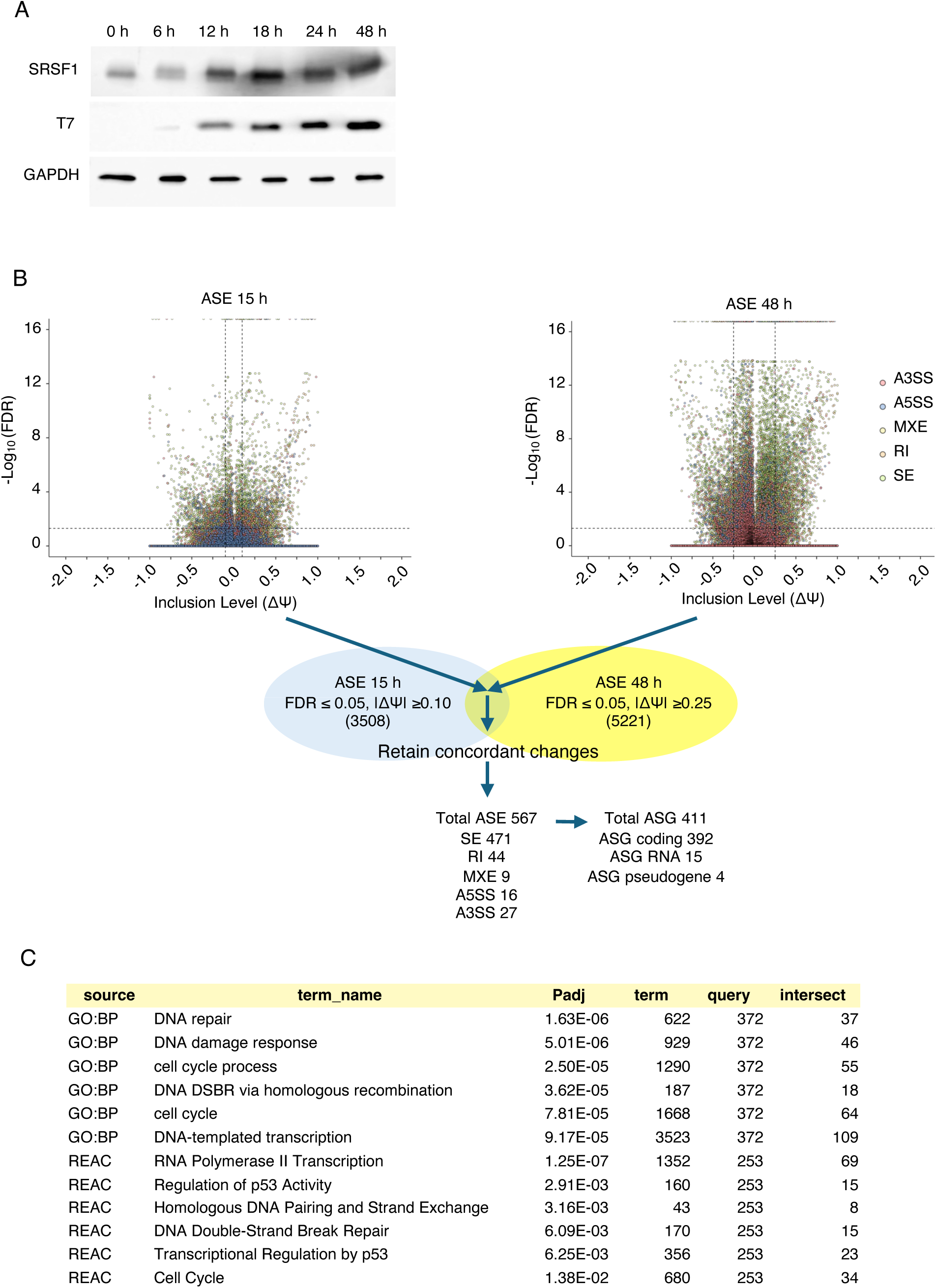
SRSF1 overexpression alters alternative splicing in genes associated with cell-cycle regulation, genome maintenance, and DNA repair. (A) Western blot analysis of endogenous and T7-tagged SRSF1 in HEK293 cells transfected with the control vector (pControl) or pSRSF1-T7. GAPDH served as a loading control. The 15- and 48-h time points were selected to examine early responses following increased SRSF1 expression and determine whether these changes persisted as SRSF1 expression increased. (B) SRSF1-regulated alternative-splicing events (ASEs) identified by rMATS analysis of HEK293 RNA-seq data. Events were selected based on |ΔΨ| ≥0.10 and FDR ≤0.05 at 15 h, |ΔΨ| ≥0.25 and FDR ≤0.05 at 48 h, and concordant regulation at both time points. (C) Functional enrichment analysis of protein-coding alternatively spliced genes (ASGs) identified biological processes associated with DNA repair, DNA-damage responses, cell-cycle regulation, homologous recombination, double-strand break repair, RNA polymerase II transcription, and p53 regulation.

### Alternative-splicing events regulated by SRSF1 are enriched in genes associated with cell-cycle control, DNA replication, and DNA-damage responses

SRSF1-regulated alternative-splicing events were identified using an FDR ≤0.05 and an absolute change in exon inclusion level (|ΔΨ|; difference in percent spliced in) ≥0.10 at 15 h and ≥0.25 at 48 h. Events were required to occur at both time points and change in the same direction. This analysis identified 567 SRSF1-regulated alternative-splicing events (ASEs) in 411 alternatively spliced genes (ASGs), including 392 protein-coding genes, 15 RNA-coding genes, and 4 pseudogenes (Fig. 1B and Appendix Table S2). The 567 events comprised 471 skipped exons, 44 retained introns, 9 mutually exclusive exons, 16 alternative 5′ splice sites, and 27 alternative 3′ splice sites (Fig. 1B).

Functional enrichment analysis of the protein-coding ASGs identified pathways associated with DNA repair, DNA-damage responses, cell-cycle regulation, homologous recombination, double-strand break repair, RNA polymerase II transcription, and p53 regulation (Fig. 1C). A total of 136 genes within these functional categories, comprising 142 nonredundant ASEs, were examined experimentally by RT-PCR. We validated 66 SRSF1-regulated splicing events in 62 genes (Fig. 2A, Supplementary Figure S2A, and Appendix Table S3). The remaining events either did not show reproducible SRSF1-dependent changes or failed to yield reliable amplification (Fig. 2B). The validated alternative splicing events affected genes modulating

**Figure 2.**
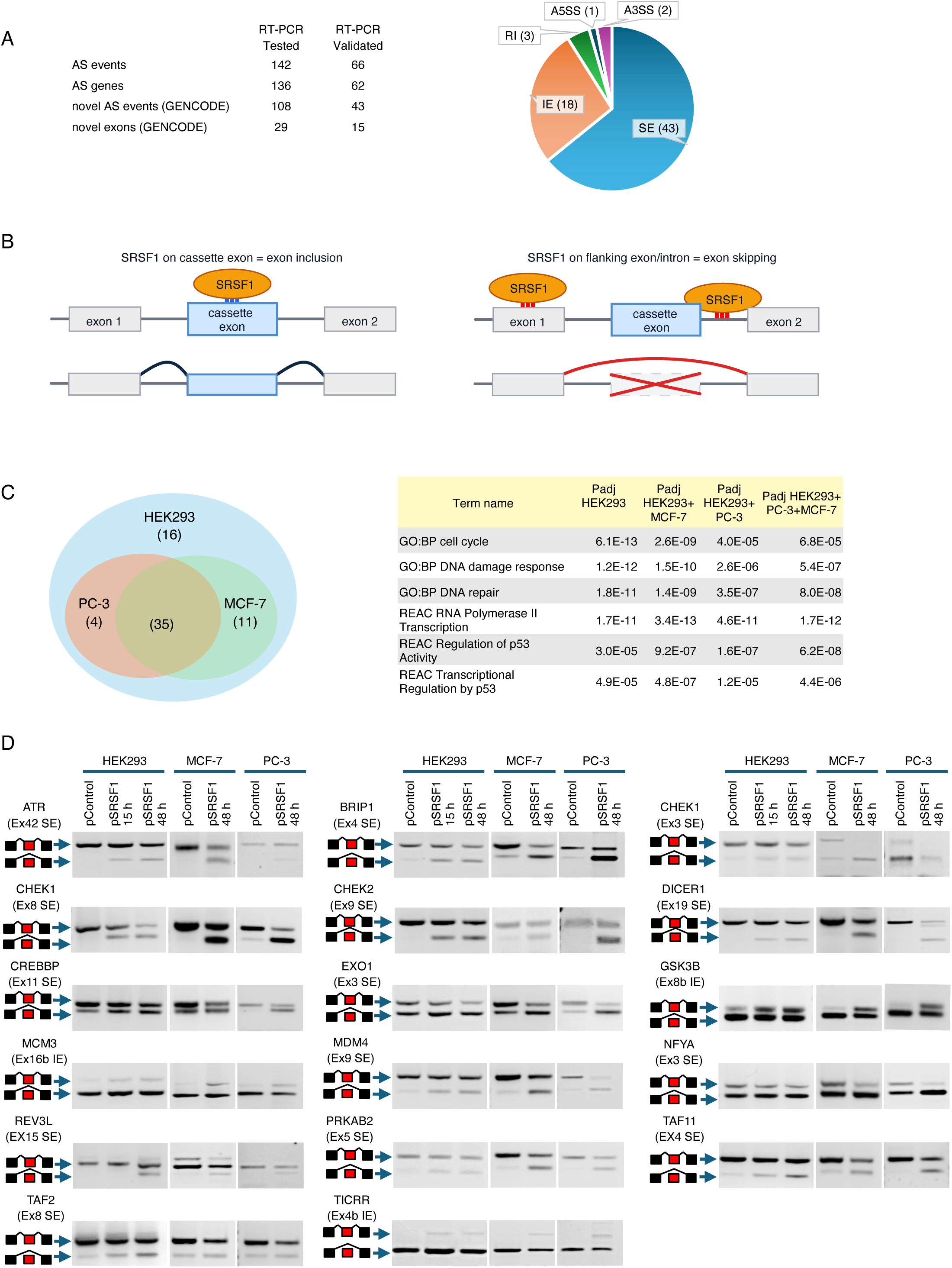
Validation and cross-cell-line reproducibility of SRSF1-regulated alternative-splicing events. **(A)** Summary of RT-PCR validation and distribution of the ASEs selected from the RNA-seq analysis. Of 142 predicted ASEs in 136 genes, 66 events in 62 genes were validated, including 43 events not annotated in GENCODE and 15 containing previously unannotated exons. The validated events comprised skipped exon (SE), included exon (IE), retained intron (RI), alternative 5′ splice site (A5SS), and alternative 3′ splice site (A3SS) events. **(B)** Schematic representation of the context-dependent effects of SRSF1 on cassette-exon splicing. SRSF1 binding within a cassette exon can promote exon inclusion, whereas binding to flanking exonic or intronic sequences can promote exon skipping. **(C)** Cross-cell-line comparison and functional enrichment analysis of SRSF1-regulated ASEs. Thirty-five events showed concordant SRSF1-dependent regulation in HEK293, PC-3, and MCF-7 cells; additional events were reproduced in one or two cell lines. Functional enrichment analysis shows selected pathways associated with cell-cycle regulation, DNA-damage responses, DNA repair, RNA polymerase II transcription, and p53 regulation. Values indicate adjusted P values. **(D)** ASEs in genes with functions in p53 regulation or p53-dependent response regulated following SRSF1 expression in HEK293, MCF-7 and PC-3 cell lines. Splicing was analyzed by RT-PCR at the indicated times after transfection with the corresponding control or SRSF1 expression plasmid. Schematics indicate the regulated exon or splicing event for each gene. Gels were cropped for presentation; uncropped source images are provided in the Source Images. Images are representative of three independent biological replicates.

DNA replication, genome maintenance, and stress-response pathways. Exon inclusion (18 events) and exon skipping (43 events) were the predominant validated event types (A2a). The presence of both event classes is consistent with the context-dependent activity of SRSF1 and indicates that increased SRSF1 expression does not uniformly promote exon inclusion (Fig. 2B). SRSF1-dependent splice-site selection is influenced by the position and sequence of local cis-regulatory elements, splice-site strength, exon architecture, and interactions with other RNA-binding proteins (22,23). We identified 43 validated SRSF1-regulated splicing events that are not annotated in GENCODE, including 15 events containing previously unannotated exons. These findings expand the set of SRSF1-responsive transcript isoforms in genes associated with cancer-relevant cellular pathways.

### SRSF1-regulated alternative-splicing events reproducible across distinct cellular backgrounds are enriched in p53-associated genes

Previous studies have linked SRSF1-dependent splicing to angiogenesis in prostate cancer (24) and to proliferation, apoptotic resistance, transformation, and tumor progression in breast cancer (2,17). We examined whether the 66 SRSF1-regulated splicing events validated in HEK293 cells also occurred in PC-3 prostate cancer and MCF-7 breast cancer cells. At 48 h after SRSF1 overexpression, 39 events in PC-3 cells and 46 in MCF-7 cells changed in the same direction as in HEK293 cells (Fig. 2C and Supplementary Figure S3). Among these, 35 events in 33 genes showed the same direction of SRSF1-dependent regulation in all three cell lines (Appendix Table S3).

Functional enrichment analysis of these 33 genes identified significant enrichment for p53 regulation and p53-dependent transcription, together with DNA repair, DNA-damage response, cell-cycle, and RNA polymerase II transcription pathways (Fig. 2C). Sixteen of the 33 genes have established roles in p53 regulation or p53-dependent functions and accounted for 17 validated events because CHEK1 contained two independently regulated alternative exons (Fig. 2D and Appendix Table S3). The three cell lines also differ in p53 functional status. MCF-7 cells retain wild-type, transcriptionally functional p53; PC-3 cells lack a functional TP53 gene; and HEK293 cells constitutively express adenoviral E1B-55K, which binds p53, alters its localization, and suppresses p53-dependent transcription (25–27). SRSF1 regulated the same 35 splicing events across these distinct p53 backgrounds, indicating that this pattern of regulation is not restricted to a single p53 functional state.

These 16 genes have established functions in replication-stress sensing, checkpoint signaling, DNA repair, metabolic-stress responses, and p53-dependent transcription (Fig. 2D, Table 1). These findings identify a reproducible pattern of SRSF1-dependent splicing among genes that regulate genome maintenance and p53 activity.

**Table 1.** p53-associated genes whose splicing is regulated by SRSF1. Functions of the SRSF1-regulated genes in p53 regulation and p53-mediated DNA-damage responses (DDR) are shown together with the predicted functional consequences of the corresponding alternative-splicing events.

| Gene | Functions in p53-mediated DDR | Predicted functional consequence of SRSF1-regulated AS |
| --- | --- | --- |
| ATR | Activates p53 directly through phosphorylation at Ser15 and Ser37 and indirectly through activation of CHEK1 and CHEK2 (36). | Predicted frameshift and loss of function likely to decrease checkpoint-driven p53 activation. |
| BRIP1 | Supports BRCA1-dependent DNA repair; BRIP1 loss promotes DNA damage and p53 activation (28). | Predicted frameshift and loss of function likely to impair fork stability, promote DSB formation, and increase DDR p53 activation. |
| CHEK1 [ex3] | Phosphorylates p53 at Ser20 and C-terminal residues in response to DNA damage, enhancing its stability (35). | Predicted frameshift and loss of function likely to impair CHEK1-dependent p53 activation and checkpoint signaling, while potentially increasing secondary DNA-damage signaling. |
| CHEK1 [ex8] | Phosphorylates p53 at Ser20 and C-terminal residues in response to DNA damage, enhancing its stability (35). | Predicted frameshift and loss of function likely to impair CHEK1-dependent p53 activation and checkpoint signaling, while potentially increasing secondary DNA-damage signaling. |
| CHEK2 | Phosphorylates p53 at Ser20, Ser15 and C-terminal in response to DNA damage, enhancing its stability (35). | Predicted frameshift and loss of function likely to decrease p53 activation in response to double-strand DNA breaks. |
| CREBBP | Functions as a coactivator of p53-dependent transcription (37). | Predicted frameshift and loss of function likely to decrease p53 transcriptional activity. |
| DICER1 | Binds unacetylated p53 and represses p53-dependent transcription through recruitment of SUV39H1 (29). | Predicted frameshift and loss of function likely to increase p53-dependent transcriptional activity. |
| EXO1 | Promotes DNA end resection and influences double-strand-break repair and checkpoint signaling (30). | Predicted frameshift and loss of function likely to reduce p53 activation following DNA damage and trigger baseline p53 activation due to genomic instability. |
| GSK3B | Phosphorylates p53 at Ser33 and increases its transcriptional activity (38). | Insertion of 13 aa within the kinase domain; predicted to alter substrate specificity and reduce GSK3β-dependent p53 activation. |
| MCM3 | Component of the MCM2–7 helicase complex, essential for replication-fork formation; MCM3 downregulation reduces CHK1/CHK2 phosphorylation (31). | Insertion of 51 aa within the helicase domain; predicted to increase replication stress that generates DSBs and activates p53 via ATR-CHK1 and ATM-CHK2. |
| MDM4 | MDM4 binds p53 and inhibits its transcriptional activity (32). | Exon 9 skipping generates MDM4-A, which lacks most of the acidic domain and may retain p53-inhibitory activity, although its endogenous function remains incompletely characterized. |
| NFYA | Binds p53 to regulate stress-induced gene expression (39). | Exon 3 skipping generates NF-YAs, which lacks a segment of the glutamine-rich transactivation domain. Predicted to alter NF-Y/p53-dependent transcription. |
| PRKAB2 | Regulatory β2 subunit of AMPK; activated AMPK promotes p53 activation during metabolic stress (40). | Predicted frameshift and loss of function likely to reduce AMPK-dependent p53 activation during metabolic stress. |
| REV3L | Supports translesion synthesis and limits persistent DNA damage; REV3L depletion activates DNA-damage signaling and p53 in p53-proficient cells (33). | Predicted frameshift and loss of function likely to cause chronic activation of p53. |
| TAF11 | Component of TFIID, which contributes to p53-dependent transcription (41). | Predicted frameshift and loss of function likely to decrease p53 target gene expression. |
| TAF2 | Component of TFIID, which contributes to p53-dependent transcription (41). | Exon 8 skipping likely to decrease p53 target gene expression. |
| TICRR | Controls replication initiation via the MCM2-7 replicative helicase complex. Its downregulation activates the ATM/CHK2 pathway (34). | Insertion of 59 aa within the C-terminal region; predicted to increase replication stress that generates DSBs and activates p53 via ATR-Chk1 and ATM-Chk2 pathways. |

### SRSF1-regulated and p53-associated splicing events occur in genes that have opposing functions in p53 signaling

We constructed a TP53-centered interaction network to define the functional relationships among p53-associated genes whose alternative splicing is regulated by SRSF1. The analysis combined high-confidence STRING interactions with experimentally supported regulatory relationships from the literature (Appendix Table S4). The network organized the SRSF1-regulated p53-associated genes according to established functions in replication stress and DNA repair, checkpoint signaling, transcriptional control, and RNA or metabolic regulation (Fig. 3A).

**Figure 3.**
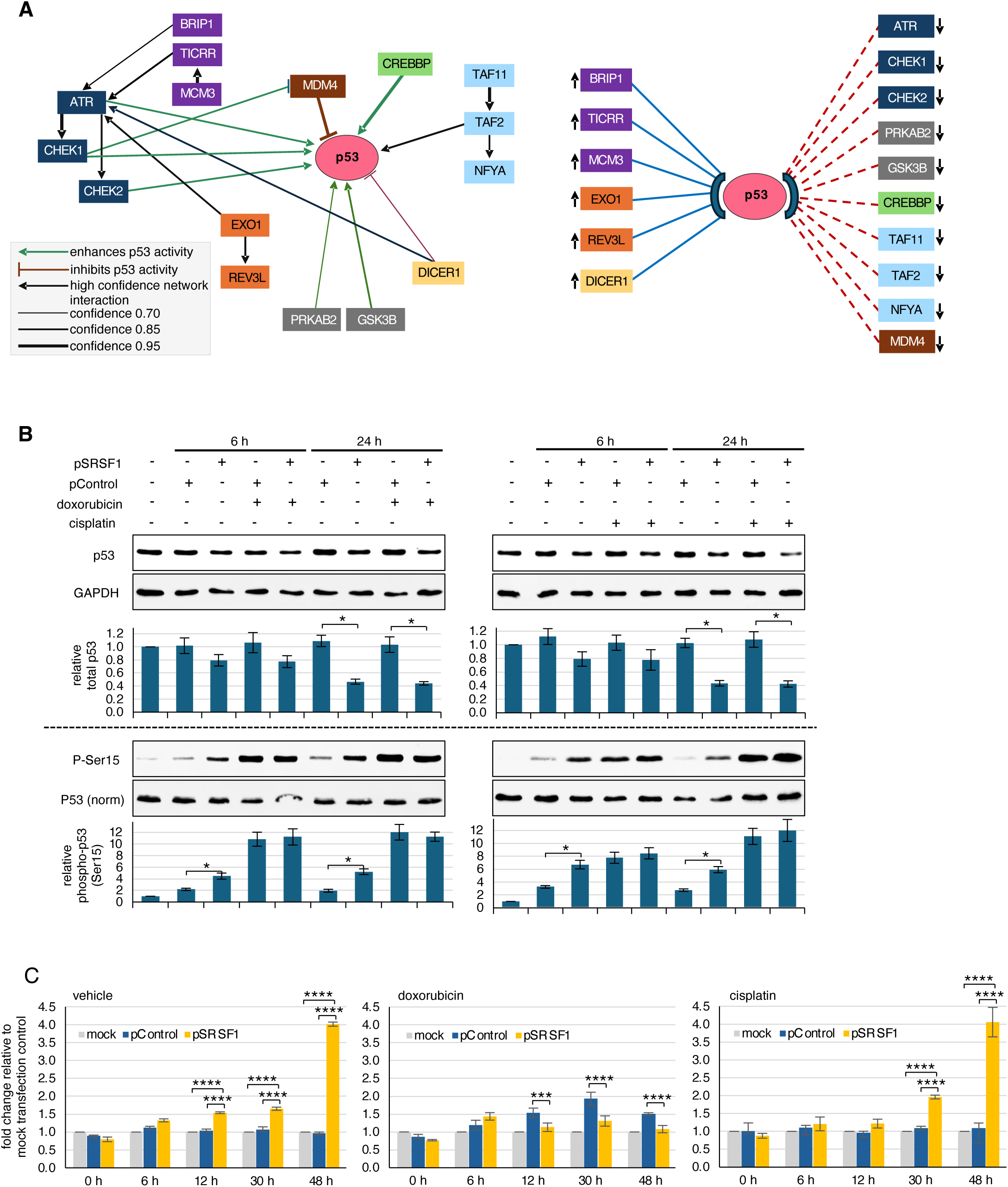
SRSF1 overexpression alters p53 protein levels, Ser15 phosphorylation, and apoptosis. **(A)** p53-centered interaction network and predicted functional consequences of SRSF1-regulated alternative-splicing events. The network was constructed from high-confidence STRING associations and experimentally supported regulatory relationships. Arrows indicate activating or inhibitory relationships, and line thickness reflects interaction confidence. Splicing events affecting BRIP1, TICRR, MCM3, EXO1, REV3L, and DICER1 were predicted to increase p53-activating stress signals, whereas events affecting ATR, CHEK1, CHEK2, PRKAB2, GSK3B, CREBBP, MDM4, TAF11, TAF2, and NFYA were predicted to attenuate checkpoint signaling, p53 stabilization, or p53-dependent transcription. **(B)** Western blot and densitometric analysis of total p53 and phospho-p53 Ser15 in HEK293 cells mock-transfected or transfected with the control empty plasmid (pControl) or SRSF1 expression plasmid (pSRSF1), followed by treatment with vehicle, doxorubicin, or cisplatin for 6 h / 24 h. Total p53 was normalized to total protein. For phospho-p53 Ser15 analysis, lysate loading was adjusted to equivalent amounts of total p53, and phospho-p53 Ser15 was normalized to total p53. Data represent mean ± SEM from three independent biological replicates. Matched pControl and pSRSF1 samples were compared using paired two-tailed Student’s t-tests (*P < 0.05). **(C)** Time-course analysis of apoptosis in HEK293 cells mock-transfected or transfected with pControl or pSRSF1 and treated with vehicle, doxorubicin, or cisplatin. Annexin V-dependent luminescence was measured at 0, 6, 12, 30, and 48 h after treatment. Values are expressed as fold change relative to the corresponding mock-transfected control and represent mean ± SEM from three independent biological replicates. Statistical significance was assessed by two-way repeated-measures ANOVA followed by Šídák-adjusted comparisons of pSRSF1 with mock and pControl at each time point. Only comparisons with P < 0.001 are indicated (***P < 0.001; ****P < 0.0001).

We integrated established gene functions, literature-supported regulatory interactions, database annotations, and analysis of the affected gene and protein sequences to evaluate the potential effects of the SRSF1-regulated isoforms on p53 signaling (Table 1, Appendix Table S4). Based on this analysis, splice changes in BRIP1, TICRR, MCM3, EXO1, REV3L, and DICER1 could increase basal or secondary signals associated with p53 activation (28–34), whereas changes in ATR, CHEK1, CHEK2, PRKAB2, GSK3B, CREBBP, MDM4, TAF11, TAF2, and NFYA could reduce checkpoint signaling, p53 stabilization, or p53-dependent transcription (35–41) (Fig. 3A). This literature-, database-, and sequence-based analysis indicates that SRSF1 regulates alternative splicing in genes with opposing roles in p53 signaling and predicts functional consequences for the affected isoforms.

### SRSF1 overexpression decreases p53 protein levels, increases basal Ser15 phosphorylation, and modulates apoptosis

The enrichment of SRSF1-regulated splicing events in genes associated with p53 regulation prompted us to examine whether SRSF1 overexpression altered cellular p53 protein levels and Ser15 phosphorylation under basal conditions and following DNA damage. We used cisplatin, which induces replication-associated DNA damage and preferentially activates ATR-CHEK1 signaling, and doxorubicin, which induces DNA double-strand breaks and activates ATM-CHEK2 signaling (42,43); both pathways promote p53 phosphorylation (35,44).

HEK293 cells transfected with the control empty plasmid or the SRSF1 expression plasmid were treated with cisplatin, doxorubicin, or vehicle. Total p53 and phospho-p53 Ser15 were analyzed 6 and 24 h after treatment. SRSF1 overexpression significantly reduced total p53 protein levels under all three conditions 24 h after treatment (Fig. 3B). Basal Ser15 phosphorylation relative to total p53 was also significantly increased by SRSF1. In contrast, following cisplatin- or doxorubicin-induced activation of p53 Ser15 phosphorylation, SRSF1 expression produced no further increase in phospho-p53 Ser15 levels (Fig. 3B). The strong induction of Ser15 phosphorylation by both genotoxic agents may limit detection of an additional SRSF1-dependent increase.

We then examined whether SRSF1-dependent changes in p53 regulation were accompanied by altered apoptosis. In vehicle-treated cells, SRSF1 overexpression produced a progressive increase in apoptosis relative to mock-transfected cells and cells transfected with the control empty plasmid (Fig. 3C). Cisplatin-treated cells showed a similar temporal pattern, with the SRSF1-dependent increase becoming evident at 30 h and markedly greater at 48 h. In contrast, doxorubicin treatment did not produce a comparable SRSF1-dependent increase in apoptosis. Apoptosis in cells transfected with pSRSF1 was not increased relative to control-transfected cells and did not exceed the response observed in cells treated with vehicle as a control or cisplatin at the later time points. These findings show that SRSF1 increases apoptosis under basal and cisplatin-treated conditions but not following doxorubicin treatment. Because both genotoxic agents strongly increased p53 Ser15 phosphorylation without a further SRSF1-dependent increase, Ser15 phosphorylation alone does not account for the different apoptotic responses.

### SRSF1 increases transcription of a distinct set of genes across different cellular backgrounds

We analyzed SRSF1-dependent changes in gene expression using the 15 and 48 h RNA-seq datasets. We identified 48 h DEGs using |log2 fold change| ≥2 and p ≤0.05 and cross-referenced them with genes showing |log2 fold change| ≥1.5 at 15 h. Because SRSF1 overexpression became detectable by 12 h, we used the 15 h dataset as an early directional-response filter to enrich for gene-expression changes occurring shortly after SRSF1 accumulation and to reduce the contribution of secondary effects that could develop by 48 h through SRSF1-dependent regulation of other transcriptional or RNA-processing factors. We selected genes that changed in the same direction at both time points. This analysis identified 128 SRSF1-responsive genes, including 56 protein-coding genes, 37 non-coding RNA genes, 24 pseudogenes, and 11 genes of unknown biotype (Fig. 4A and Appendix Table S5). Comparison of the 56 protein-coding SRSF1-responsive genes with the 392 protein-coding genes containing SRSF1-regulated alternative-splicing events revealed no overlap between the two groups (Fig. 4B), indicating that SRSF1-dependent changes in gene expression and alternative splicing affected distinct gene sets. Of the 56 protein-coding genes, 55 showed increased expression by RNA-seq. RT-qPCR analysis confirmed concordant increases for 32 of these genes, with ≥2-fold increases at 15 h and ≥4-fold increases at 48 h (Fig. 4 and Supplementary Figure S4). TRIM34, the only protein-coding gene identified as downregulated by RNA-seq, did not show a reproducible change by RT-qPCR at either time point.

**Figure 4.**
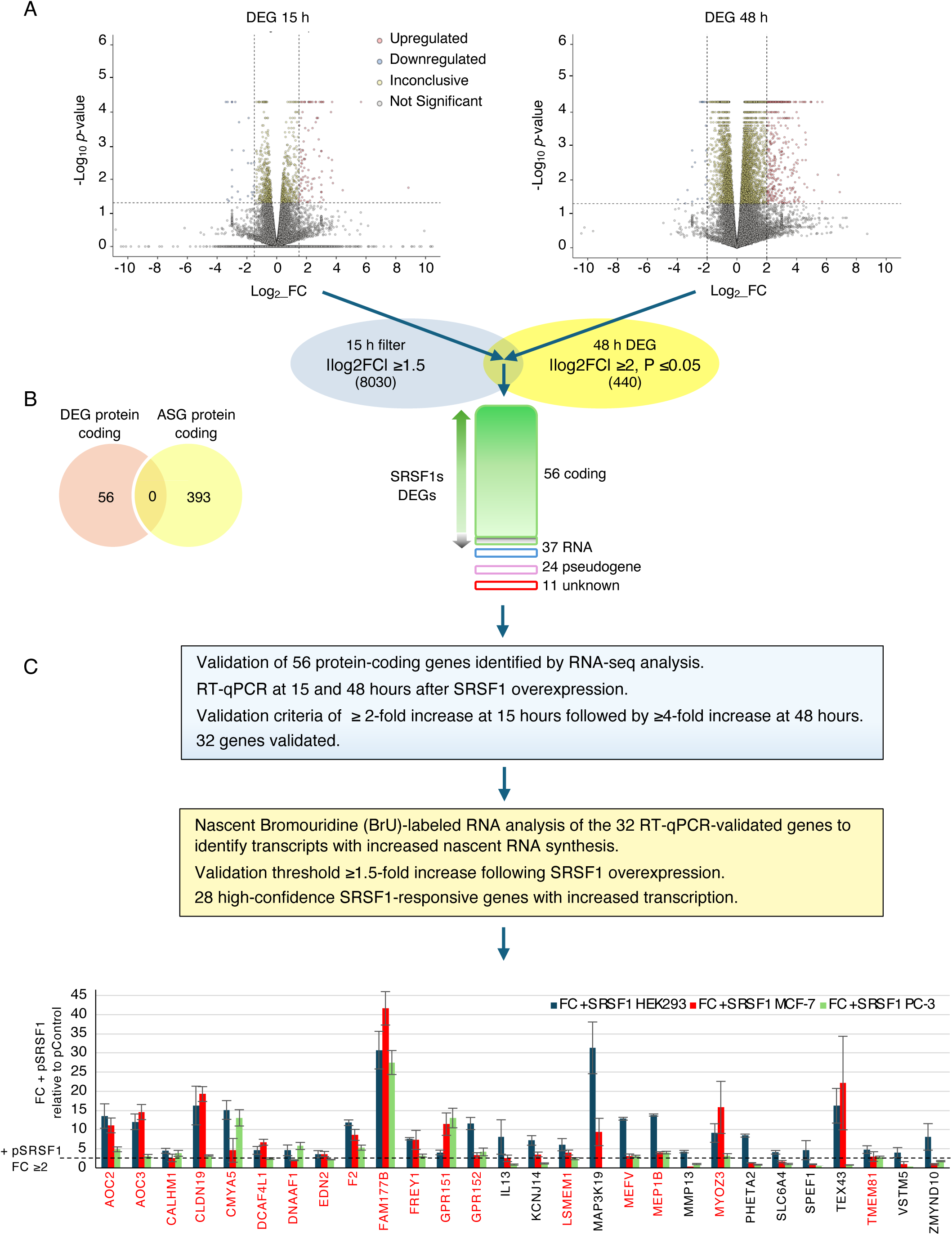
Identification and cross-cell-line validation of SRSF1-responsive genes with increased transcription. **(A)** Identification of SRSF1-responsive genes by time-course RNA-seq. Volcano plots show differential gene expression 15 and 48 h after SRSF1 overexpression. Genes with |log2 fold change| ≥2 and P ≤0.05 at 48 h were cross-referenced with genes showing |log2 fold change| ≥1.5 at 15 h and concordant regulation at both time points. The 15 h dataset served solely as an early directional-response filter and was not treated as an independently significant DEG set. This analysis identified 128 SRSF1-responsive genes, including 56 protein-coding genes, 37 non-coding RNA genes, 24 pseudogenes, and 11 genes of other or undetermined biotypes. **(B)** Comparison of the 56 protein-coding SRSF1-responsive genes with the 393 protein-coding ASGs identified by RNA-seq. The two groups showed no overlap. **(C)** Experimental validation of SRSF1-responsive protein-coding genes. The 56 protein-coding SRSF1-responsive genes identified by RNA-seq were analyzed by RT-qPCR at 15 and 48 h after SRSF1 overexpression. Thirty-two genes showed ≥2-fold increased RNA levels at 15 h and ≥4-fold increased RNA levels at 48 h and were considered validated. BrU nascent RNA analysis was then performed on these 32 RT-qPCR-validated genes. Twenty-eight genes showed ≥1.5-fold increased newly synthesized RNA following SRSF1 overexpression and were considered validated targets of SRSF1-dependent transcriptional activation. Cross-cell-line analysis of these 28 genes identified 18 with ≥2-fold increased RNA levels in HEK293, MCF-7, and PC-3 cells; these genes are indicated in red. Data represent three independent biological replicates, each analyzed in duplicate technical reactions. Values represent mean fold change relative to the corresponding control-transfected cells ± SEM from three independent biological replicates.

Next, we measured nascent RNA by BrU labeling to distinguish increased RNA synthesis from altered RNA degradation. Among the 32 RT-qPCR-validated genes, 28 showed a ≥1.5-fold increase in BrU-labeled RNA following SRSF1 overexpression (Fig. 4 and Supplementary Figure S4B), indicating that their increased steady-state RNA levels were associated with increased RNA synthesis. We considered these 28 genes validated targets of SRSF1-dependent transcriptional activation. Furthermore, we examined five SRSF1-responsive transcripts and the RPL13A reference transcript following transcriptional inhibition with actinomycin D to determine whether SRSF1-dependent changes reflected altered RNA stability (14,45). All SRSF1-responsive transcripts analyzed showed comparable decay kinetics in control and SRSF1-overexpressing cells (Supplementary Figure S5), further supporting increased RNA synthesis rather than enhanced RNA stability as the primary basis for their increased expression. We also examined whether the 28 validated SRSF1-responsive genes showed similar regulation in PC-3 and MCF-7 cells. Eighteen genes increased by at least twofold in all three cell lines (Fig. 4C), demonstrating reproducible SRSF1-dependent regulation across distinct cellular backgrounds. Functional annotation associated these genes with signaling, extracellular communication, adhesion, cytoskeletal organization, transcriptional regulation, and protein regulation (Table 2). The differential regulation of the remaining ten genes indicates that SRSF1-dependent transcriptional regulation is influenced in part by cellular context.

**Table 2.** SRSF1-dependent transcriptional targets upregulated across HEK293, MCF-7, and PC-3 cells. Genes whose expression increased by ≥2-fold following SRSF1 overexpression in all three cell lines, as measured by RT-qPCR. Genes are grouped by GO/Reactome functional categories, including inflammatory, GPCR, and calcium signaling; adhesion and barrier regulation; cytoskeletal remodeling; transcriptional and protein regulation; and ciliary or motile processes.

| Gene | Description | GO/REACTOME functional category | Functions |
| --- | --- | --- | --- |
| AOC2 | amine oxidase, copper containing 2 | GO Metabolic process | Amine oxidase involved in metabolic processing and oxidative stress regulation. |
| AOC3 | amine oxidase, copper containing 3 | GO Cell adhesion<br>GO Inflammatory response<br>GO Metabolic process | Adhesion-associated amine oxidase regulating leukocyte trafficking and extracellular matrix remodeling. |
| CALHM1 | calcium homeostasis modulator 1 | GO Signal transduction | Membrane channel protein modulating calcium signaling and cellular stress responses |
| CLDN19 | claudin 19 | GO Cell adhesion<br>REAC: Cell junction organization | Tight junction protein maintaining epithelial barrier integrity and cell–cell adhesion. |
| CMYA5 | cardiomyopathy associated 5 | GO Cytoskeleton organization,<br>GO Cell adhesion | Cytoskeletal anchoring protein contributing to structural organization and cell adhesion. |
| DCAF4L1 | DDB1 and CUL4 associated factor 4 like 1 | GO Regulation of transcription by RNA polymerase II | Ubiquitin ligase adaptor influencing protein turnover and transcriptional regulation. |
| DNAAF1 | dynein axonemal assembly factor 1 | GO Cytoskeleton organization<br>REAC Cytoskeletal remodeling processes | Microtubule-associated factor required for ciliary assembly and cytoskeletal organization. |
| EDN2 | endothelin 2 | GO Signal transduction<br>GO Chemotaxis | Secreted peptide ligand mediating GPCR signaling, chemotaxis, and paracrine communication. |
| F2 | coagulation factor II, thrombin | GO Signal transduction | Coagulation protease influencing inflammatory signaling and tumor microenvironment interactions. |
| FAM177B | family with sequence similarity 177 member B | GO Regulation of transcription by RNA polymerase II | Possible role in the regulation of gene expression programs. |
| FREY1 | Frey regulator of sperm-oocyte fusion 1 | GO Cytoskeleton organization | Associated with the endoplasmic reticulum; possible role in protein assembly and stability. |
| GPR151 | G protein-coupled receptor 151 | GO Signal transduction | Orphan G protein–coupled receptor mediating intracellular signal transduction. |
| GPR152 | G protein-coupled receptor 152 | GO Signal transduction | Orphan G protein–coupled receptor involved in modulatory signaling pathways. |
| LSMEM1 | leucine rich single-pass membrane protein 1 | GO Signal transduction | Small membrane protein involved in signal transduction and cellular communication. |
| MEFV | MEFV, pyrin innate immunity regulator | GO Inflammatory response | Innate immunity protein regulating inflammasome activation and inflammatory stress responses. |
| MEP1B | meprin A subunit beta | GO Extracellular matrix organization<br>GO Metabolic process | Metalloprotease involved in extracellular matrix processing and metabolic protein cleavage. |
| MYOZ3 | myozenin 3 | GO Cytoskeleton organization | Actin–myosin interaction protein contributing to cytoskeletal organization. |
| TMEM81 | transmembrane protein 81 | GO Cell–cell junction organization | Transmembrane protein involved in cell junction organization and membrane signaling. |

### SRSF1-dependent transcriptional activation is regulated by the 7SK snRNP

Previous studies showed that SR proteins can regulate transcription elongation through the 7SK small nuclear ribonucleoprotein (snRNP), which sequesters P-TEFb in an inactive state (18,19,21). P-TEFb is a heterodimeric kinase complex composed of CDK9 and a T-type cyclin, most commonly cyclin T1 (CCNT1). To determine whether the 7SK snRNP regulates SRSF1-dependent activation of cellular genes, we examined the effects of 7SK snRNA depletion and HEXIM1 or HEXIM2 overexpression on five validated SRSF1-responsive genes: AOC2, AOC3, MEFV, MYOZ3, and TEX43.

Antisense oligonucleotides targeting 7SK snRNA efficiently reduced 7SK levels (Fig. 5B) and increased expression of all five genes, consistent with disruption of 7SK-dependent P-TEFb sequestration and increased availability of P-TEFb for RNA polymerase II elongation (Fig. 5A). Under these conditions, the additional transcriptional activation produced by SRSF1 was attenuated for all five genes, whereas the control antisense oligonucleotide had no effect on SRSF1-dependent activation. Together, these results link SRSF1-dependent gene activation to regulation of P-TEFb by the 7SK snRNP.

**Figure 5.**
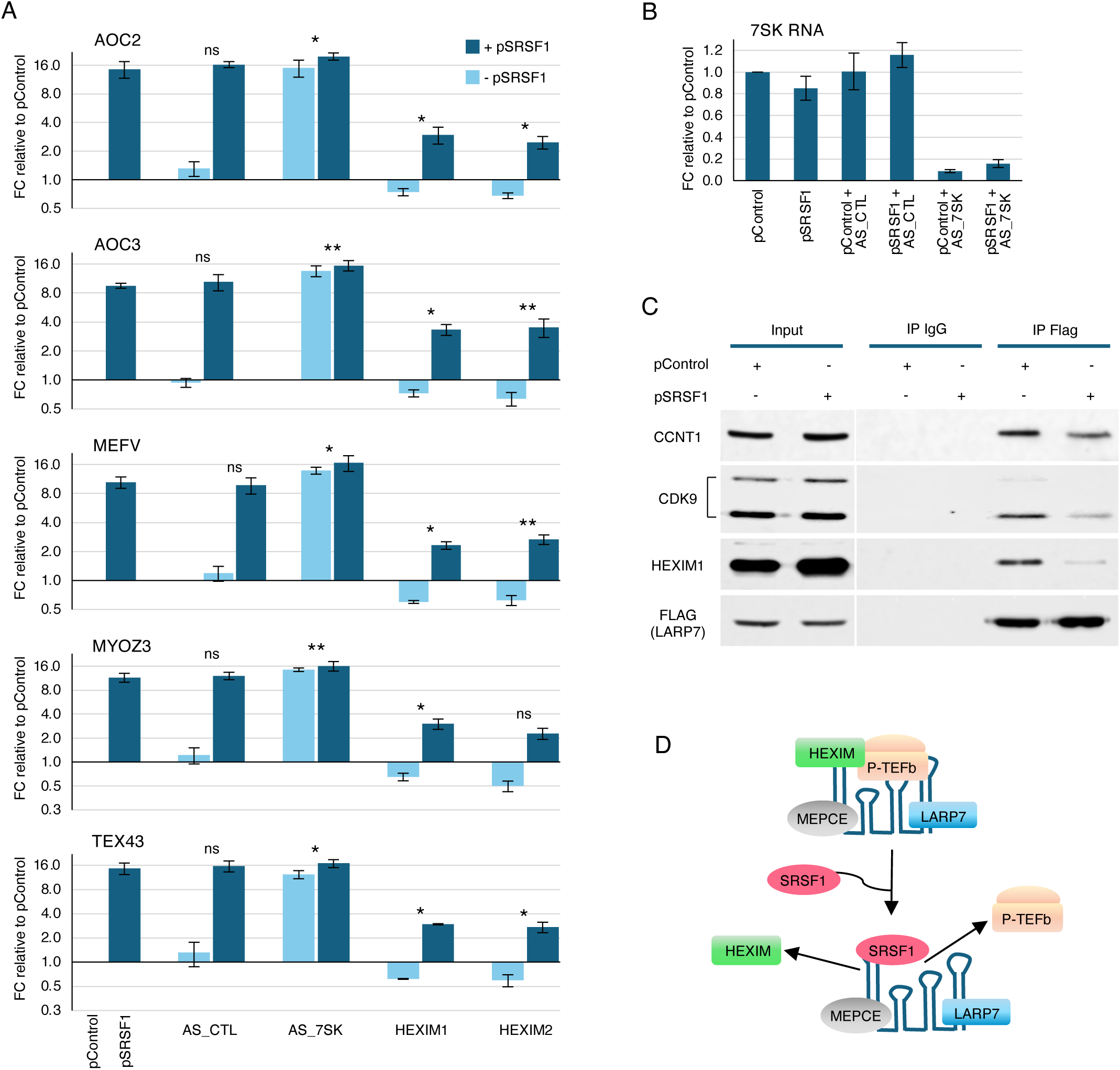
SRSF1 promotes transcriptional activation of cellular genes by facilitating P-TEFb mobilization from the 7SK snRNP. **(A)** Relative expression of AOC2, AOC3, MEFV, MYOZ3, and TEX43 in cells transfected with pControl or pSRSF1, alone or together with control antisense oligonucleotides (AS_CTL), 7SK-targeting antisense oligonucleotides (AS_7SK), or HEXIM1/HEXIM2 expression vectors. RT-qPCR values were normalized to pControl and are shown as mean ± SEM from three independent experiments. SRSF1-dependent activation was compared across treatments using replicate-specific interaction contrasts and two-sided one-sample Student’s t-tests against zero, as described in Methods. \**P* < 0.05; **P < 0.01; ns, not significant. **(B)** RT-qPCR analysis of 7SK snRNA in cells expressing pControl or pSRSF1 following treatment with AS_CTL or AS_7SK. Values are relative to pControl. **(C)** Effect of SRSF1 overexpression on association of P-TEFb and HEXIM1 with the 7SK snRNP. FLAG-LARP7 was immunoprecipitated from cells expressing pControl or pSRSF1, and input lysates and immunoprecipitates were analyzed for CCNT1, CDK9, HEXIM1, and FLAG-LARP7. SRSF1 overexpression reduced recovery of CCNT1, CDK9, and HEXIM1 with FLAG-LARP7. IgG immunoprecipitation served as a negative control. **(D)** Mechanism of SRSF1-mediated P-TEFb mobilization. The 7SK snRNP contains 7SK snRNA, LARP7, MEPCE, HEXIM proteins, and P-TEFb (CDK9–Cyclin T1). Increased SRSF1 expression facilitates P-TEFb mobilization from the 7SK snRNP to promote transcription elongation of SRSF1-responsive genes.

Next, we examined whether increasing HEXIM1 or HEXIM2 expression affected SRSF1-dependent gene activation. HEXIM proteins bind 7SK snRNA and promote a conformational rearrangement that facilitates P-TEFb recruitment into the 7SK snRNP. HEXIM then inhibits P-TEFb by masking the catalytic site of CDK9. Mobilization of P-TEFb from the HEXIM-containing 7SK snRNP relieves CDK9 inhibition, generating active P-TEFb that promotes RNA polymerase II pause release and productive elongation. Overexpression of either HEXIM protein reduced expression of all five genes in the absence of SRSF1 overexpression (Fig. 5A). HEXIM1 significantly attenuated SRSF1-dependent activation of all five genes.

To determine whether SRSF1 alters P-TEFb association with the 7SK snRNP, we immunoprecipitated FLAG-tagged LARP7, a constitutive core component of the 7SK snRNP that binds and stabilizes 7SK snRNA, from control and SRSF1-overexpressing cells and analyzed the associated proteins. SRSF1 overexpression reduced recovery of HEXIM1, CDK9, and CCNT1 with LARP7 (Fig. 5C), indicating reduced association of HEXIM1 and P-TEFb with the 7SK snRNP. Together, these results support a model in which SRSF1 promotes P-TEFb mobilization from the 7SK snRNP, thereby increasing P-TEFb availability for RNA polymerase II pause release and productive transcription elongation (Fig. 5D).

### SRSF1 expression associates with target-gene expression across tumors and normal tissues

Although The Cancer Genome Atlas (TCGA) and Genotype-Tissue Expression (GTEx) datasets support large-scale transcriptomic analyses, limited junction coverage and low variation in several validated splicing events restricted our assessment of SRSF1-associated alternative splicing. We instead examined the comparatively less-characterized transcriptional functions of SRSF1 by analyzing 28 experimentally validated targets, including the 18 whose activation was conserved across HEK293, MCF-7, and PC-3 cells, in 33 TCGA tumor cohorts and 47 GTEx normal-tissue cohorts. We evaluated associations using continuous SRSF1 expression and comparisons between SRSF1-high and SRSF1-low samples, defined by the upper and lower expression quartiles.

After adjustment for composition covariates, continuous SRSF1 expression showed significant positive associations with the 18- and 28-gene expression scores in 19 and 20 TCGA cohorts, respectively, and with both scores in 40 GTEx tissues (FDR-adjusted *P* <0.05; Fig. 6A). High-versus-low comparisons generally showed concordant associations, although fewer reached statistical significance. Because these comparisons did not reveal a more consistent pattern, we used continuous SRSF1 expression for subsequent analyses, retaining all eligible samples and the full expression range.

**Figure 6.**
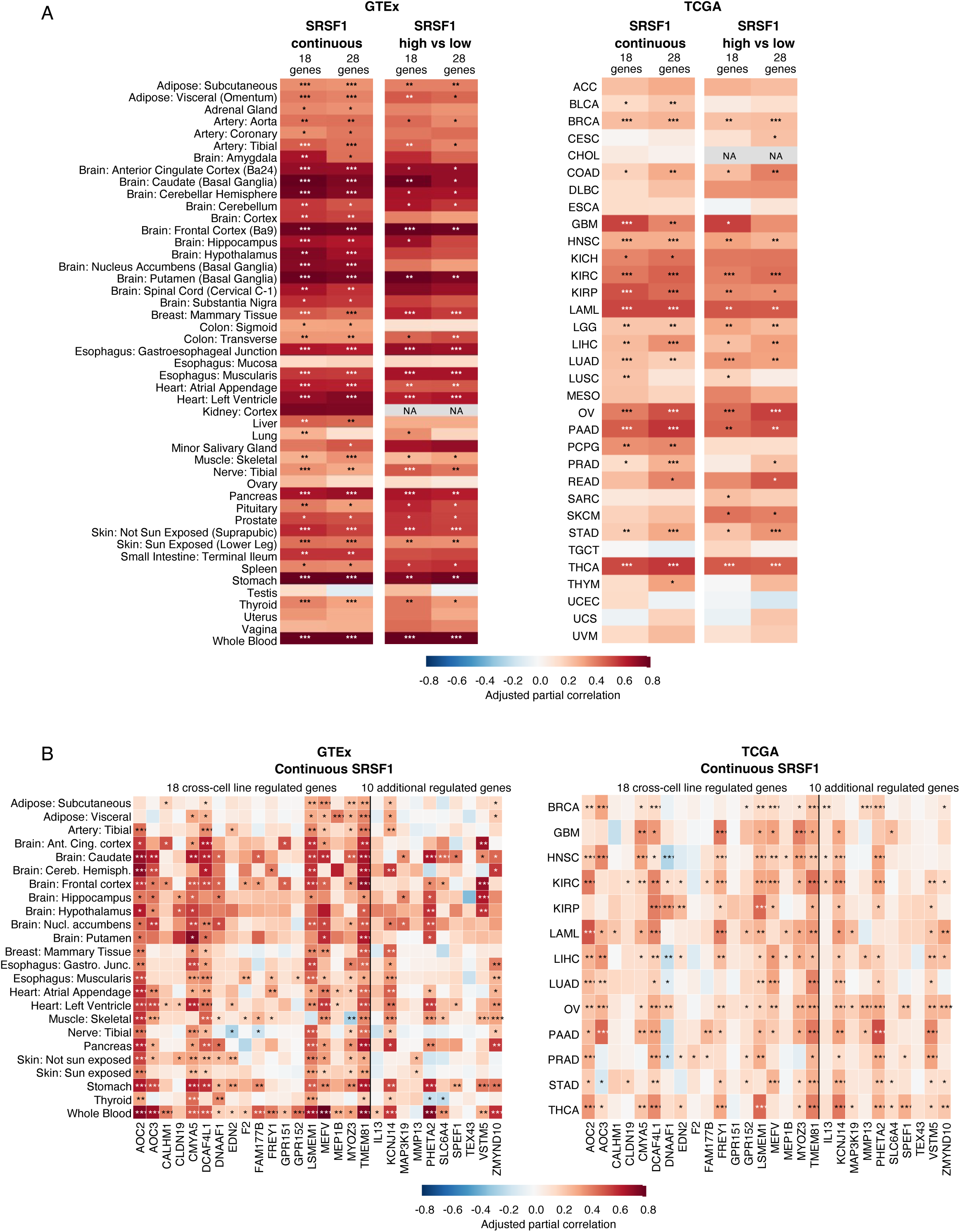
SRSF1 expression correlates with experimentally validated transcriptionally regulated genes across tumors and normal tissues. (A) Associations between SRSF1 expression and the 18- and 28-gene expression scores across 47 normal-tissue cohorts from the Genotype-Tissue Expression (GTEx) project and 33 tumor cohorts from The Cancer Genome Atlas (TCGA). The 28-gene panel comprises genes whose transcriptional activation by SRSF1 was experimentally validated; the 18-gene subset includes those activated across HEK293, MCF-7, and PC-3 cells. Continuous analyses examined SRSF1 expression across all eligible samples, whereas high-versus-low comparisons included the upper and lower expression quartiles, excluding the middle 50%. (B) Associations between continuous SRSF1 expression and individual genes in 24 GTEx tissues and 13 TCGA cancers selected because at least one gene-set score met FDR-adjusted P < 0.000005. Colors indicate adjusted partial correlations, which measure the strength and direction of the association between SRSF1 and gene expression after accounting for sample composition and log -transformed junction-library size. Two-sided regression tests used heteroscedasticity-consistent HC3 standard errors. Benjamini–Hochberg correction controlled the false discovery rate (FDR) within prespecified testing families; individual-gene analyses retained correction across all 2,240 planned comparisons. Asterisks indicate FDR-adjusted P values: * < 0.05, ** < 0.0005, and *** < 0.000005. Absence of asterisks indicates FDR-adjusted P ≥ 0.05; NA indicates unavailable estimates.

Individual-gene analyses of the 13 cancers and 24 normal tissues meeting the gene-set significance threshold revealed widespread positive associations across multiple transcriptional targets (Fig. 6B). TMEM81 showed significant associations in all 13 cancers and 24 tissues, while DCAF4L1 was significant in all 13 cancers and 22 tissues. LSMEM1, AOC2, MEFV, and CMYA5 also showed recurrent positive correlations. Particularly large numbers of targets were associated with SRSF1 in whole blood, stomach, frontal cortex, and left ventricular myocardium, as well as ovarian, clear-cell renal, and thyroid carcinomas. The 18 targets conserved across the three experimental cell lines generally showed more widespread associations in normal tissues than the ten additional genes. However, KCNJ14, PHETA2, VSTM5, and IL13 exhibited recurrent positive associations in several cancers, indicating that targets not conserved across all three cell lines also contribute to the expression patterns observed in human tumors.

We further examined SRSF1 expression and its transcriptional targets across breast cancer molecular subtypes and in relation to thyroid differentiation (Supplementary Figure S6A–B). SRSF1 expression differed significantly among breast cancer subtypes, with lower levels in basal-like tumors than in luminal A, luminal B, and HER2-enriched cancers. Despite these differences, AOC3, DCAF4L1, MEFV, PHETA2, and TMEM81 showed significant positive associations with SRSF1 in both luminal A and basal-like tumors. Eight additional targets were significant in luminal A but not basal-like tumors, including AOC2, CMYA5, GPR152, and VSTM5, whereas FREY1, KCNJ14, SPEF1, and ZMYND10 were significant only in basal-like tumors. Fewer associations reached significance in luminal B and HER2-enriched tumors. These findings identify shared and subtype-associated transcriptional targets, although differences in significance do not establish differences in association strength between subtypes.

In thyroid carcinoma, SRSF1 expression showed a weak positive association with the thyroid differentiation score that did not remain significant after FDR correction. In contrast, DNAAF1, DCAF4L1, VSTM5, SPEF1, and SLC6A4 retained significant positive associations with thyroid differentiation after adjustment for SRSF1 expression and clinical and composition covariates, whereas TMEM81 showed a negative association (Supplementary Figure S6B). These findings identify distinct relationships between experimentally validated SRSF1 transcriptional targets and thyroid tumor differentiation.

Finally, exploratory survival analyses in clear-cell renal carcinoma identified distinct clinical associations among SRSF1-regulated transcriptional targets, although SRSF1 expression itself was not significantly associated with overall survival (OS) or progression-free interval (PFI).

Higher TMEM81 and MMP13 expression was associated with both outcomes, corresponding to approximately 25% and 20% higher hazards of death and 35% and 20% higher hazards of PFI events, respectively. Higher LSMEM1 and DCAF4L1 expression was also associated with approximately 25% and 15% higher hazards of death, whereas CMYA5 expression was associated with a 30% lower hazard of a PFI event (Supplementary Figure S6C). In contrast, no individual target showed a significant association with PFI in pancreatic adenocarcinoma after multiple-testing correction (Supplementary Figure S6D). Together, these analyses identify widespread associations between SRSF1 and its transcriptional targets, alongside gene-specific relationships with tumor subtype, differentiation, and clinical outcomes.

### Integrated model of SRSF1-dependent transcriptional and alternative-splicing programs

Integrated analysis of the transcriptional and alternative-splicing datasets showed that SRSF1 increased transcription of genes associated with signaling, extracellular communication, adhesion and motility, cytoskeletal organization, and gene regulation (Fig. 7). This transcriptional activation was linked to SRSF1-dependent P-TEFb mobilization from the 7SK snRNP. TCGA and GTEx analyses further identified widespread positive associations between SRSF1 expression and these experimentally regulated genes across human tumors and normal tissues. In cancer, SRSF1 expression differed among breast cancer subtypes, while individual regulated genes showed distinct associations with thyroid differentiation and clinical outcomes in clear-cell renal carcinoma.

**Figure 7.**
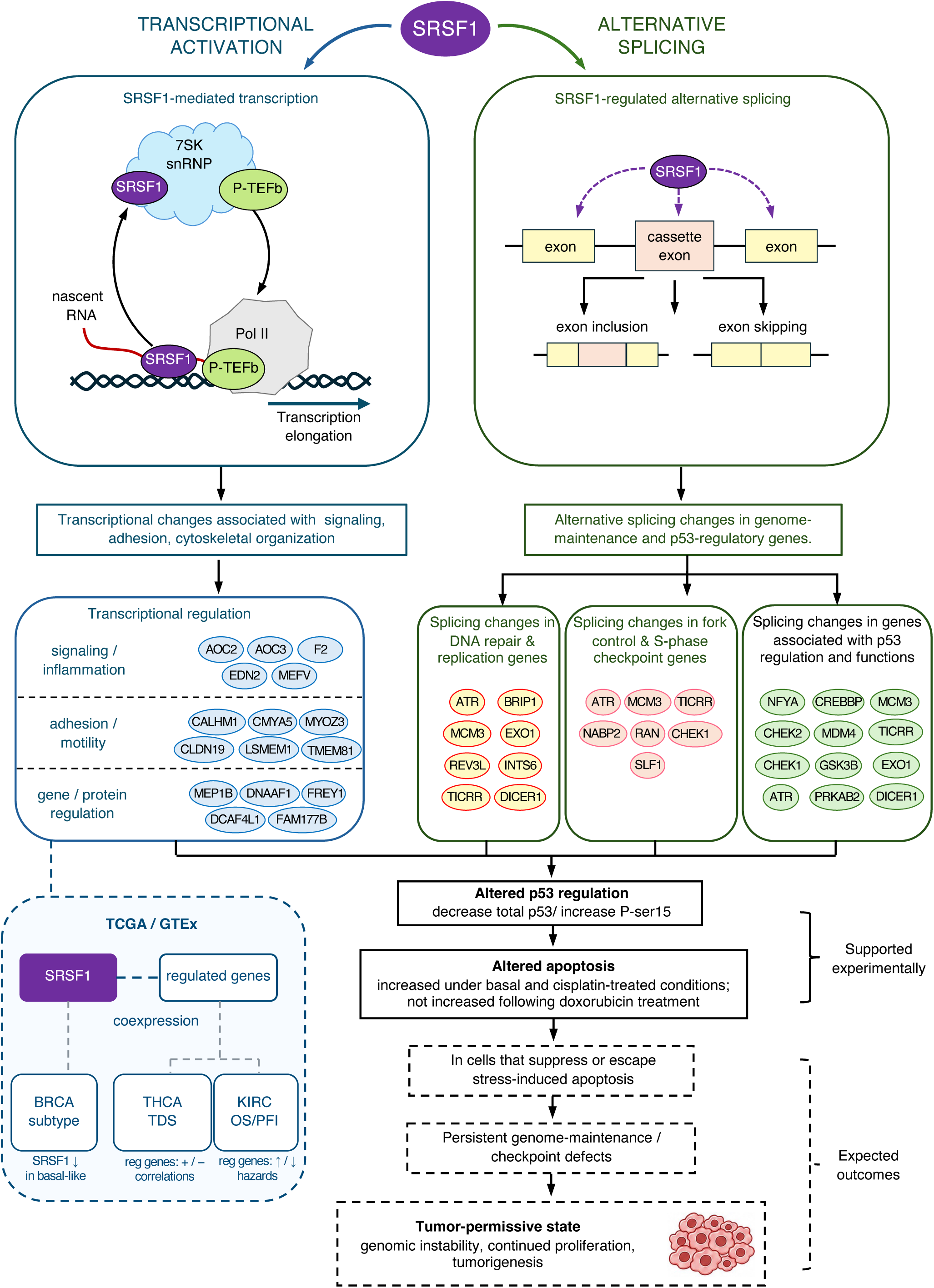
Integrated model of SRSF1-dependent transcription and alternative splicing. Schematic representation of SRSF1-dependent transcriptional activation through positive transcription elongation factor b (P-TEFb) regulation by the 7SK small nuclear ribonucleoprotein (snRNP), alternative splicing of genome-maintenance and p53-regulatory genes, and changes in p53 expression, phosphorylation, and apoptosis. Analyses of TCGA and GTEx datasets illustrate SRSF1-associated gene expression and relationships with breast cancer (BRCA) subtype, thyroid carcinoma (THCA) differentiation score (TDS), and kidney renal clear cell carcinoma (KIRC) overall survival (OS) and progression-free interval (PFI). The correlation plot is schematic. Solid connections indicate experimental findings, dashed blue connections indicate observational associations, and dashed black connections indicate proposed outcomes. Positive (+) and negative (−) symbols indicate correlation direction; upward (↑) and downward (↓) arrows indicate increases and decreases in the indicated measurements.

In parallel, SRSF1 modulated alternative splicing of genes controlling replication-fork function, DNA repair, DNA-damage responses, cell-cycle checkpoints, and p53 regulation. SRSF1 overexpression also reduced total p53 protein levels, increased basal Ser15 phosphorylation, and increased apoptosis. Together, these findings identify concurrent SRSF1-dependent transcriptional and alternative-splicing programs associated with cellular signaling, genome maintenance, p53 regulation, and stress responses. The integrated model combines experimentally supported changes with gene-expression and clinical associations in human cancers and distinguishes these findings from proposed downstream consequences of persistent genome-maintenance and checkpoint defects.

### Discussion

SRSF1 is an RNA-binding protein that regulates pre-mRNA splicing and functions as a proto-oncogene. Our findings identify two distinct gene-regulatory programs induced by increased SRSF1 expression: increased transcription of endogenous genes linked to P-TEFb mobilization from the 7SK snRNP and alternative-splicing changes enriched in genes controlling genome maintenance and p53 regulation. SRSF1 overexpression also reduced total cellular p53 and increased basal Ser15 phosphorylation. Functionally, SRSF1 increased apoptosis under basal conditions and following cisplatin treatment but not following doxorubicin treatment, indicating that the effect of SRSF1 on apoptosis differs according to the genotoxic stressor.

Previous studies established that loss of SRSF1 compromises genome stability through R-loop accumulation (46,47). R-loops are RNA:DNA hybrids whose persistence can impede replication-fork progression and promote DNA double-strand breaks and chromosomal rearrangements (46,47). SRSF1 depletion increases R-loop formation, replication-fork stalling, and DNA damage, whereas RNase H-mediated removal of R-loops suppresses these effects. SRSF1-dependent mRNA export was also shown to cooperate with FANCI/FANCD2 to restrict R-loop accumulation and DNA damage (48). These studies establish an important role for SRSF1 in genome maintenance. Consistent with these observations, the present study indicates that an increase in SRSF1 expression correlates with altered isoform ratios in genes regulating replication-fork function, DNA repair, checkpoint signaling, and p53 activity.

Seventeen of the 35 SRSF1-regulated splicing events detected across HEK293, PC-3, and MCF-7 cells affected 16 genes with established roles in p53 regulation or p53-dependent cellular functions. Literature, database, and sequence analyses predicted opposing effects of these splice changes on p53 signaling, although most of the affected isoforms have not been functionally characterized experimentally. Previous studies showed that increased SRSF1 can stabilize p53 through an RPL5-dependent mechanism and promote senescence in primary cells (49). In contrast, SRSF1 overexpression in HEK293 cells reduced total cellular p53 while increasing basal Ser15 phosphorylation and apoptosis, indicating that the apoptotic response cannot be explained solely by p53 abundance. Because HEK293 cells express adenoviral E1B-55K, which suppresses p53-dependent transcription (27), the apoptotic phenotype may also reflect p53-independent stress responses activated by SRSF1-dependent changes in DNA replication, DNA repair, and checkpoint genes.

The increase in apoptosis following SRSF1 overexpression does not conflict with its characterization as a proto-oncogene. SRSF1 can promote transformation and reduce apoptosis through regulation of specific splice isoforms (2), whereas increased SRSF1 can also activate cellular stress responses and promote senescence (49), indicating that its effects depend on cellular context and the nature of the cellular stress. In the present study, SRSF1-dependent changes in genes controlling DNA replication, replication-fork stability, DNA repair, and checkpoint signaling may generate cellular stress sufficient to promote apoptosis. The similar SRSF1-dependent apoptotic responses in vehicle-treated and cisplatin-treated cells indicate that this response is induced by SRSF1 and is not substantially enhanced by cisplatin. Cisplatin generates replication-blocking DNA lesions that activate ATR-CHEK1 signaling and increase dependence on replication- and repair-associated processes also altered by SRSF1-dependent splicing, consistent with the similar apoptotic profiles observed under the two conditions (28,30,31,33–36,42). In contrast, doxorubicin primarily induces DNA double-strand breaks and strong ATM-dependent signaling (43) and did not produce a comparable SRSF1-dependent increase in apoptosis, suggesting that the cellular response to doxorubicin modifies the SRSF1-dependent apoptotic phenotype. Both cisplatin and doxorubicin strongly increased p53 Ser15 phosphorylation, and SRSF1 produced no additional increase under either treatment (35,44). Together with the SRSF1-dependent reduction in total p53, these findings indicate that neither p53 abundance nor Ser15 phosphorylation alone explains the treatment-specific apoptotic responses. In cells that suppress or escape this stress-induced response, persistent alterations in genome maintenance and checkpoint control, together with SRSF1-dependent transcriptional changes, could instead favor genomic instability and continued proliferation and thereby contribute to a tumor-permissive state.

Our data identify 28 cellular genes whose transcription is increased by SRSF1. BrU labeling supported increased nascent RNA synthesis, while actinomycin D experiments showed that the higher steady-state RNA levels were not explained by altered RNA stability. Eighteen of these transcriptional targets responded similarly across HEK293, PC-3, and MCF-7 cells, indicating that SRSF1 regulates a conserved set of endogenous genes across distinct cellular backgrounds. This conserved transcriptional program is linked to 7SK snRNP-mediated control of P-TEFb. The effects of 7SK snRNA depletion and HEXIM1/2 overexpression, together with reduced association of P-TEFb and HEXIM1 with LARP7 after SRSF1 overexpression, support a model in which SRSF1 promotes P-TEFb mobilization from the 7SK snRNP and increases P-TEFb availability for transcription elongation. The present results extend our previous work showing SRSF1-dependent regulation of 7SK/P-TEFb in HIV-1 transcription to endogenous cellular genes and are consistent with earlier evidence that SR proteins promote P-TEFb release from 7SK (19,21). Direct measurements of P-TEFb activity and RNA polymerase II pause release at individual target loci will further define this mechanism.

The TCGA and GTEx analyses extend the relevance of the experimentally defined SRSF1 transcriptional program to human tumors and normal tissues, where SRSF1 expression showed widespread positive associations with the same genes across diverse cohorts. In breast cancer, basal-like tumors expressed less SRSF1 than other molecular subtypes, yet several regulated genes maintained positive correlations with SRSF1 within basal-like tumors. In thyroid carcinoma, several SRSF1-regulated genes remained associated with tumor differentiation after accounting for SRSF1 expression. In clear-cell renal carcinoma, higher TMEM81 and MMP13 expression was associated with increased hazards of both death and progression-free interval (PFI) events, whereas CMYA5 showed an association with PFI in the opposite direction. LSMEM1 and DCAF4L1 were also associated with increased mortality hazards, while SRSF1 itself showed no significant association with either clinical outcome. These findings indicate that the relationships between SRSF1-regulated gene expression and cancer characteristics vary across tumor types and may reflect additional transcriptional controls and tumor-specific biological processes. Although the TCGA/GTEx findings extend the relevance of the experimentally regulated genes to human cancers, these observational associations do not establish direct transcriptional regulation by SRSF1 in tumors or causal relationships with differentiation and disease progression and will require independent validation.

Transcription and alternative splicing are functionally connected, and several RNA-binding proteins, splicing factors, and transcription-associated proteins participate in both processes. Extensive work has shown that RNA polymerase II elongation, CTD phosphorylation, chromatin organization, and recruitment of RNA-processing factors can influence splice-site selection, while splicing factors and other RNA-processing proteins can in turn affect transcription elongation and chromatin-associated events (50–53). These interactions extend beyond SR proteins and include multiple classes of cellular factors that connect transcription with co-transcriptional RNA processing. Our results show that increased SRSF1 expression produces separate transcriptional and alternative-splicing programs with different functional outputs. The transcriptional program is linked to 7SK/P-TEFb mobilization and regulates genes associated with cellular signaling and organization, whereas the splicing program modulates genome-maintenance and p53-regulatory genes. Thus, SRSF1 can concurrently regulate distinct sets of cellular functions through transcription and alternative splicing.

The conserved transcriptional and alternative-splicing responses across HEK293, MCF-7, and PC-3 cells support their broader relevance, while the accompanying changes in p53 regulation and apoptosis connect these molecular programs to cellular stress responses. The TCGA/GTEx findings further establish associations between the experimentally regulated genes and human cancer characteristics, including molecular subtype, differentiation, and clinical outcomes. Together, these observations support an integrated model in which SRSF1 coordinates transcriptional activation and alternative splicing, with potential consequences for genome maintenance, checkpoint control, and cellular stress responses (Fig. 7). Cells that survive SRSF1-induced stress may retain alterations that favor a tumor-permissive state, although this proposed outcome requires experimental validation. As the prototypical member of the SR protein family, SRSF1 also provides a basis for determining whether other SR proteins similarly coordinate transcription and alternative splicing or regulate these processes through distinct mechanisms. Several SR proteins have been associated with tumorigenesis and cancer progression, making it important to establish which activities are shared across the family, which are protein specific, and how their combined effects influence genome maintenance, cell proliferation, stress responses, and malignant progression.

## Supporting information

Source Data and Statistical Analysis

Source images RT-PCR

Source images Western Blot

## Data Availability

RNA-sequencing data generated in this study are available through the Gene Expression Omnibus (GEO) under accession GSE124397 (https://www.ncbi.nlm.nih.gov/geo/query/acc.cgi?acc=GSE124397). Numerical source data underlying RT-qPCR, BrU nascent RNA, RNA-stability, p53 densitometry, apoptosis, TCGA/GTEx statistical analyses are provided in Source Data. Uncropped western blot and RT-PCR source images are provided in Source images WB and Source images RT-PCR gels, respectively. No custom code was generated for this study.

## Supplementary Data

Supplementary Data are available at NAR Online.

## Author contributions

Conceptualization, M.C.; methodology, M.C. and S.P.; investigation, S.P. and E.B.; validation, S.P. and E.B.; formal analysis, M.C. and S.P.; visualization, S.P. and E.B.; supervision, M.C.; project administration, M.C.; funding acquisition, M.C.; writing – original draft, M.C.; writing – review & editing, M.C. and S.P. All authors reviewed and approved the final manuscript.

## Acknowledgements

We thank Dr. Michael Lu for helpful discussions on cellular cancer models, Evan Clark for establishing the initial RNA-sequencing analysis pipeline, and Anastasia Ritchie for assistance with cell culture and gene-expression validation. Use of AI and large language models (LLMs) was limited to language editing with Microsoft Copilot. All scientific interpretation, data analysis, and final manuscript content were written, reviewed, and approved by the authors.

