## Supplementary Figure S1 for "SRSF1 coordinates transcription and alternative splicing to modulate p53 and apoptosis"

**Experimental design for RNA-seq analysis.** Experimental workflow for RNA-seq analysis of HEK293 cells transfected with the control vector (pControl) or pSRSF1-T7 and harvested 15 and 48 h after transfection. Total RNA underwent rRNA depletion, library preparation, paired-end sequencing, alignment to GRCh38, and analysis of differential gene expression and alternative-splicing events (ASEs), followed by experimental validation.

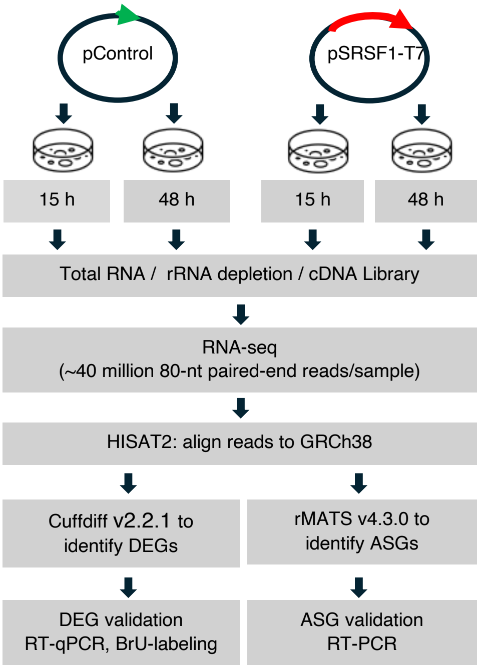
