## Supplementary Figure S2 for "SRSF1 coordinates transcription and alternative splicing to modulate p53 and apoptosis"

**RT-PCR validation of SRSF1-regulated alternative-splicing events identified by RNA-seq. (A)** RT-PCR analysis of SRSF1-regulated alternative-splicing events in HEK293 cells transfected with control or SRSF1 expression vectors and analyzed at 15 and 48 h. Schematics show the predicted splicing event and expected isoforms for each gene. Red gene names indicate events also regulated in MCF-7 and PC-3 cells; asterisks mark genes associated with p53 regulation.

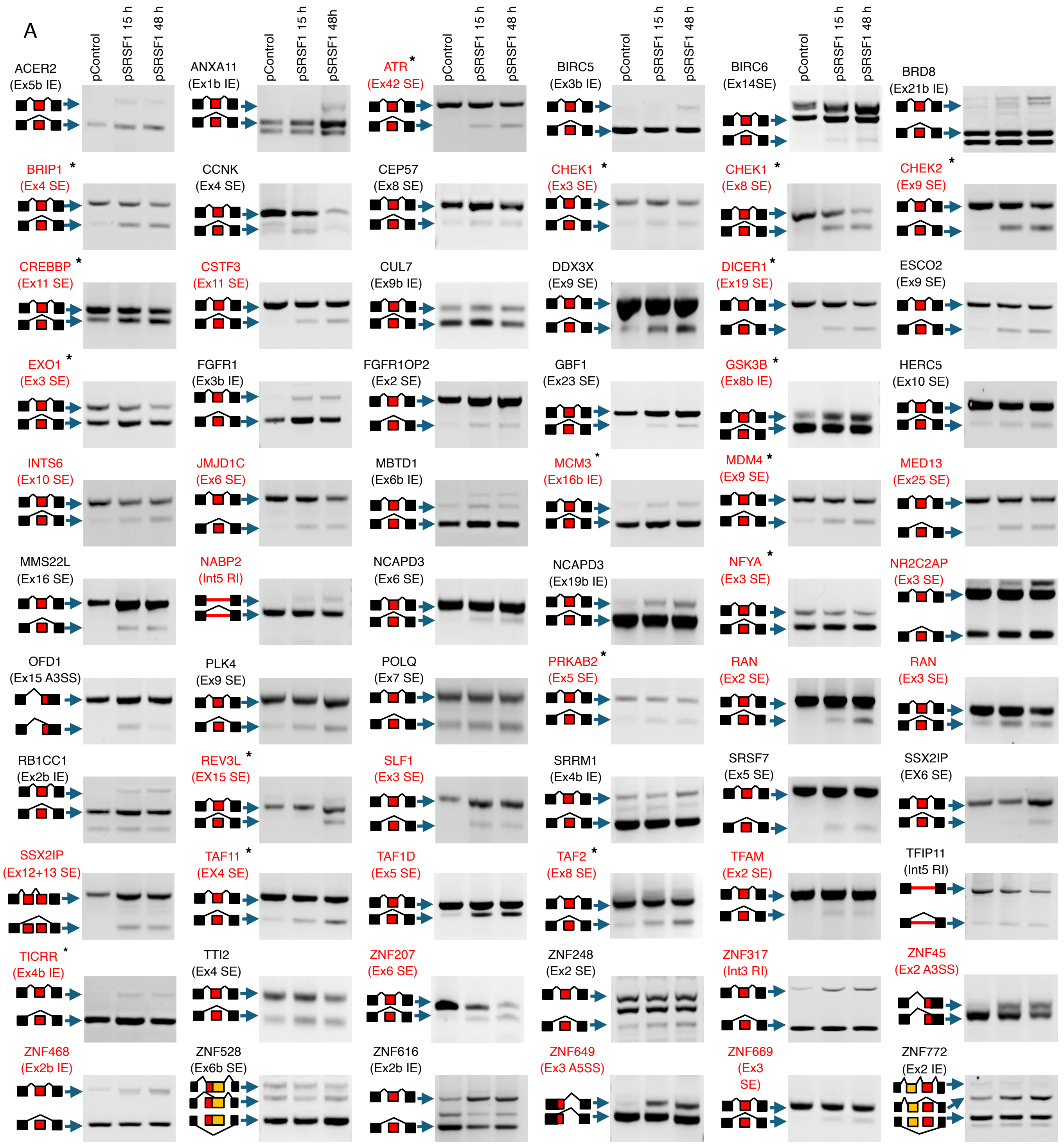

**Supplementary Figure S2.**

**(B)** RT-PCR analysis of predicted SRSF1-regulated events that did not meet validation criteria or could not be reliably amplified in HEK293 cells. Schematics show the predicted event for each gene. Images are representative of three independent biological replicates. Gels were cropped for presentation; uncropped source images are provided in the Source Images.

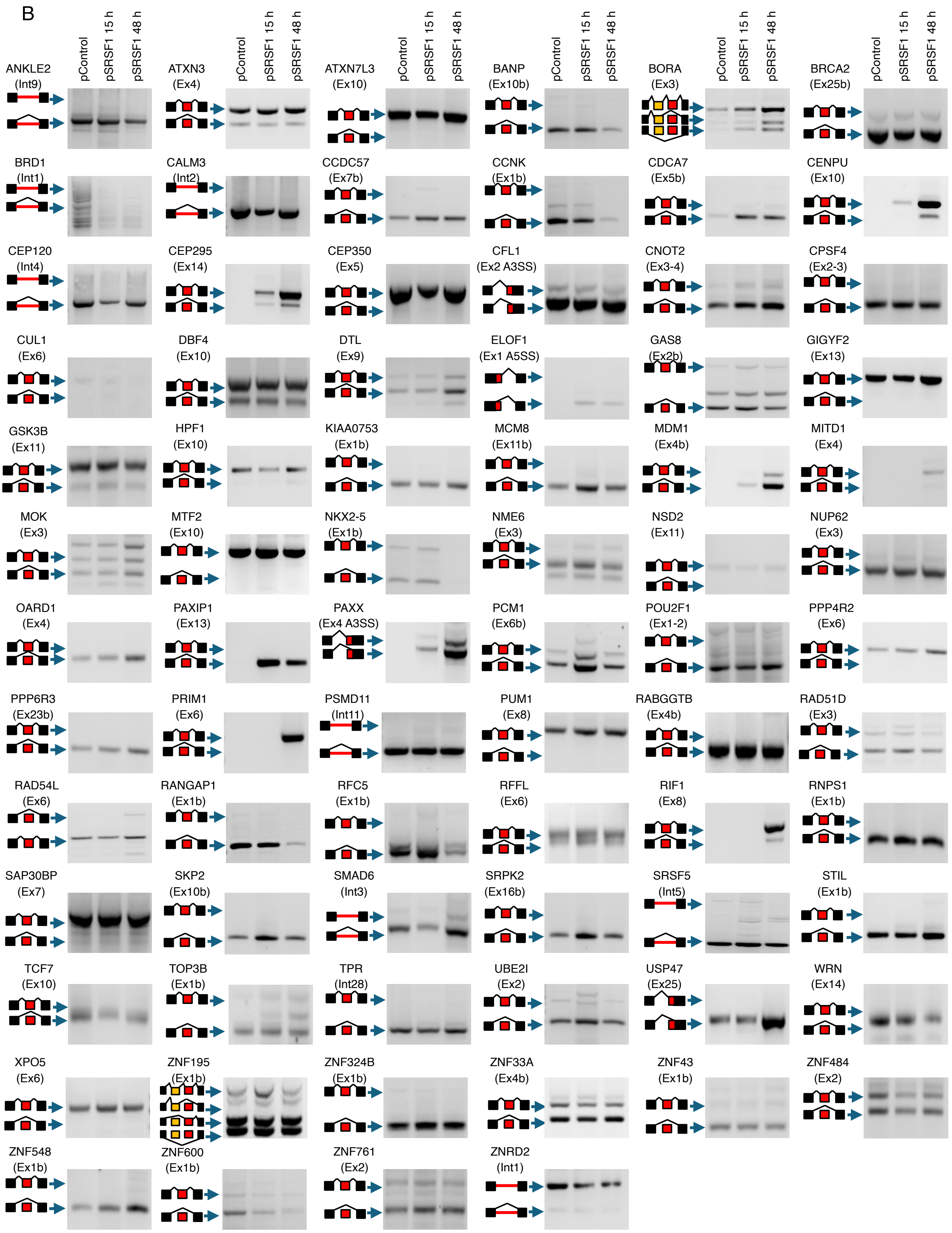
