## Supplementary Figure S3 for "SRSF1 coordinates transcription and alternative splicing to modulate p53 and apoptosis"

**Validation of SRSF1-regulated alternative-splicing events in MCF-7 and PC-3 cells. (A)** RT-PCR analysis of SRSF1-regulated alternative-splicing events in MCF-7 cells transfected with control or SRSF1 expression vectors. Schematics show the predicted splicing events and expected isoforms. Red gene names indicate events regulated in HEK293, MCF-7, and PC-3 cells; asterisks mark genes associated with p53 regulation. **(B)** RT-PCR analysis of predicted SRSF1-regulated events that did not meet validation criteria or could not be reliably amplified in MCF-7 cells.

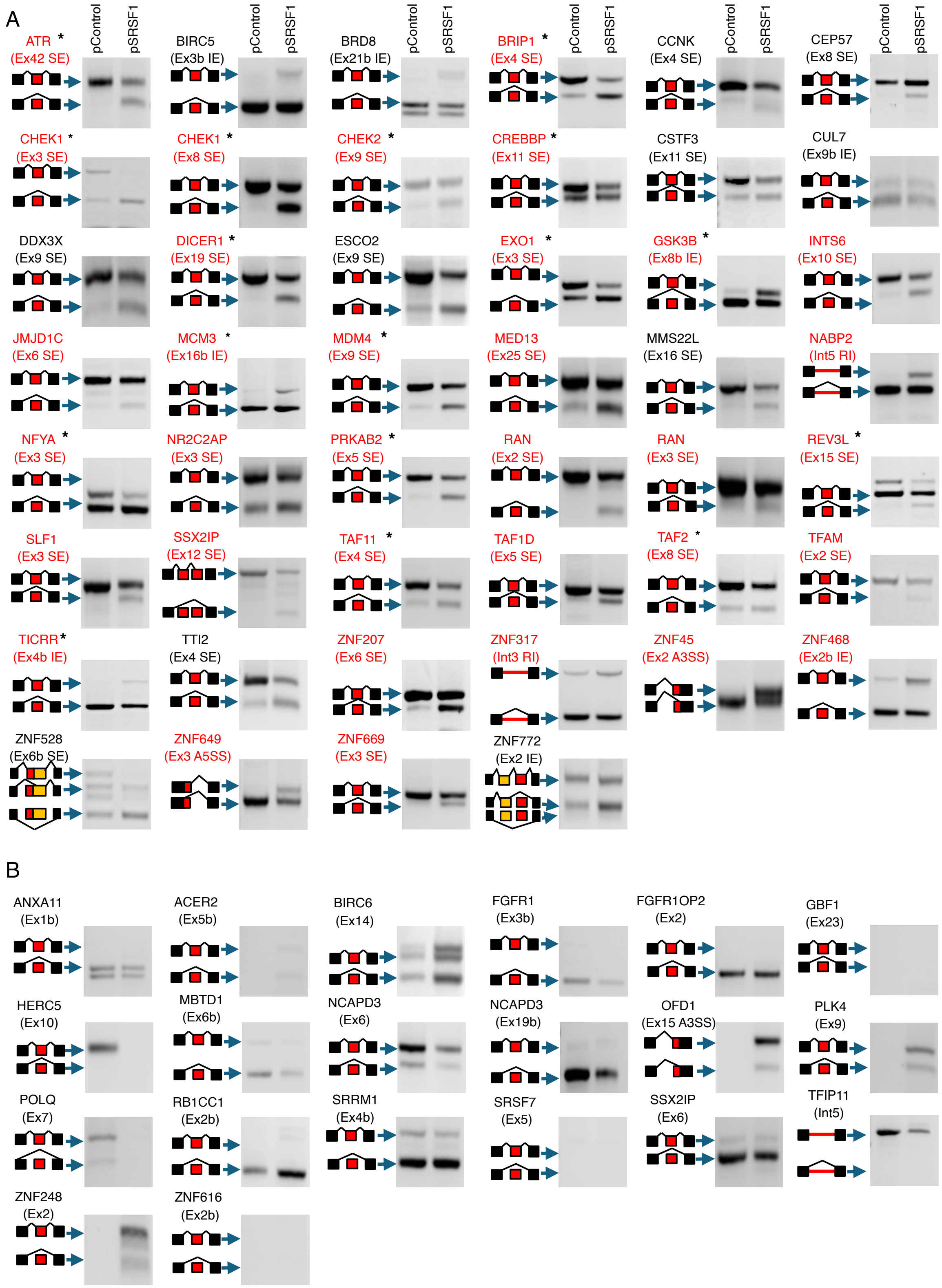

**Supplementary Figure S3.**

**(C)** RT-PCR analysis of SRSF1-regulated alternative-splicing events in PC-3 cells transfected with control or SRSF1 expression vectors. Schematics, red gene names, and asterisks are defined as in panel a. **(D)** RT-PCR analysis of predicted SRSF1-regulated events that did not meet validation criteria or could not be reliably amplified in PC-3 cells. Images are representative of three independent biological replicates. Gels were cropped for presentation; uncropped source images are provided in the Source Images.

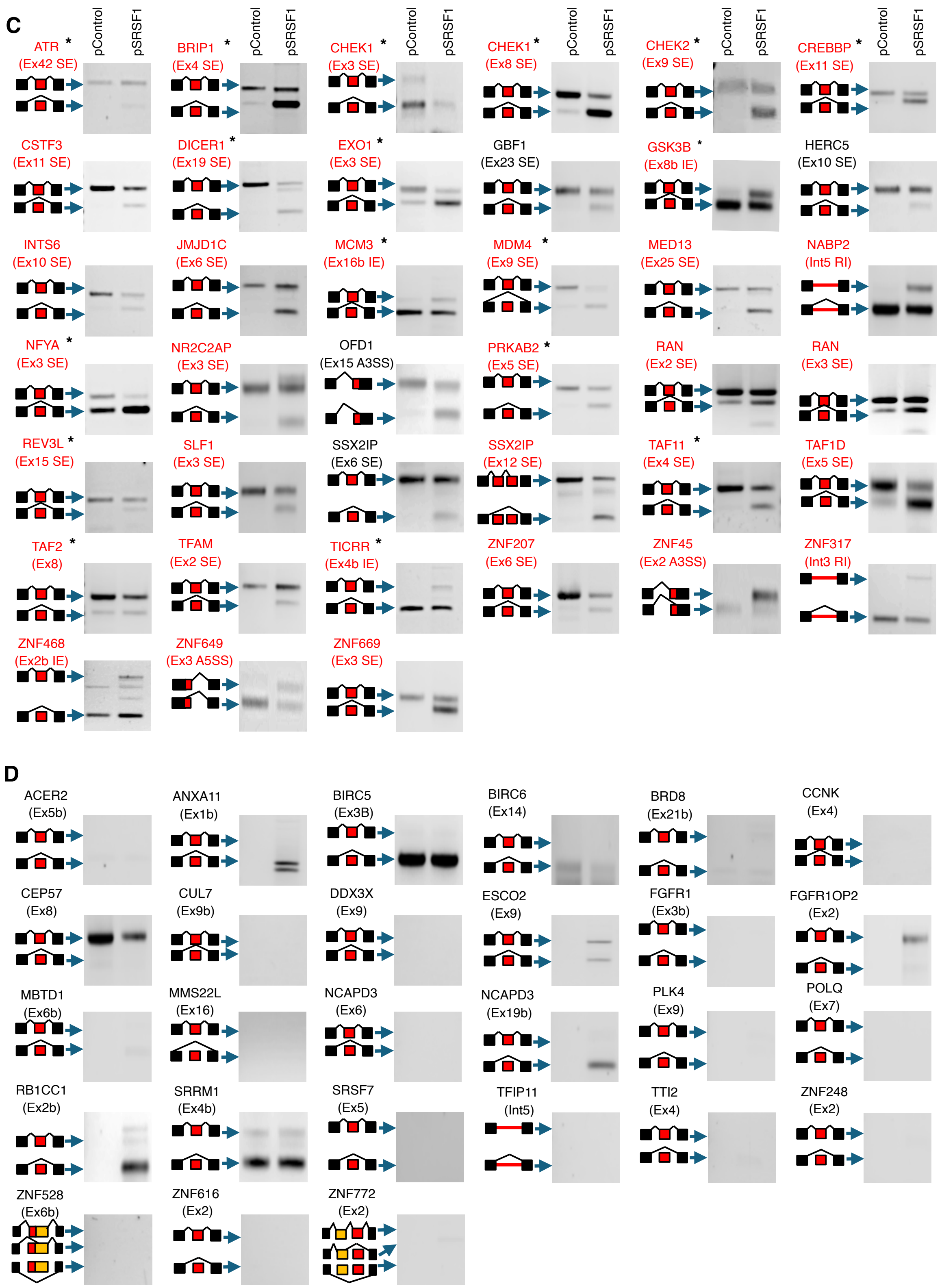
