## Supplementary Figure S4 for "SRSF1 coordinates transcription and alternative splicing to modulate p53 and apoptosis"

**Experimental validation of SRSF1-responsive genes. (A)** RT-qPCR validation of protein-coding SRSF1-responsive genes identified by RNA-seq. Gene expression was measured 15 and 48 h after SRSF1 overexpression and expressed as fold change relative to control-transfected cells. Dashed lines indicate the validation thresholds of ≥2-fold at 15 h and ≥4-fold at 48 h; validated genes are shown in red. **(B)** BrU nascent RNA analysis of RT-qPCR-validated genes. BrU-labeled RNA was immunoprecipitated and quantified by RT-qPCR to identify transcripts with increased nascent RNA synthesis following SRSF1 overexpression. Samples processed without BrU served as negative controls. The dashed line indicates the ≥1.5-fold validation threshold; validated genes are shown in red. Data represent mean ± SEM from three independent biological replicates.

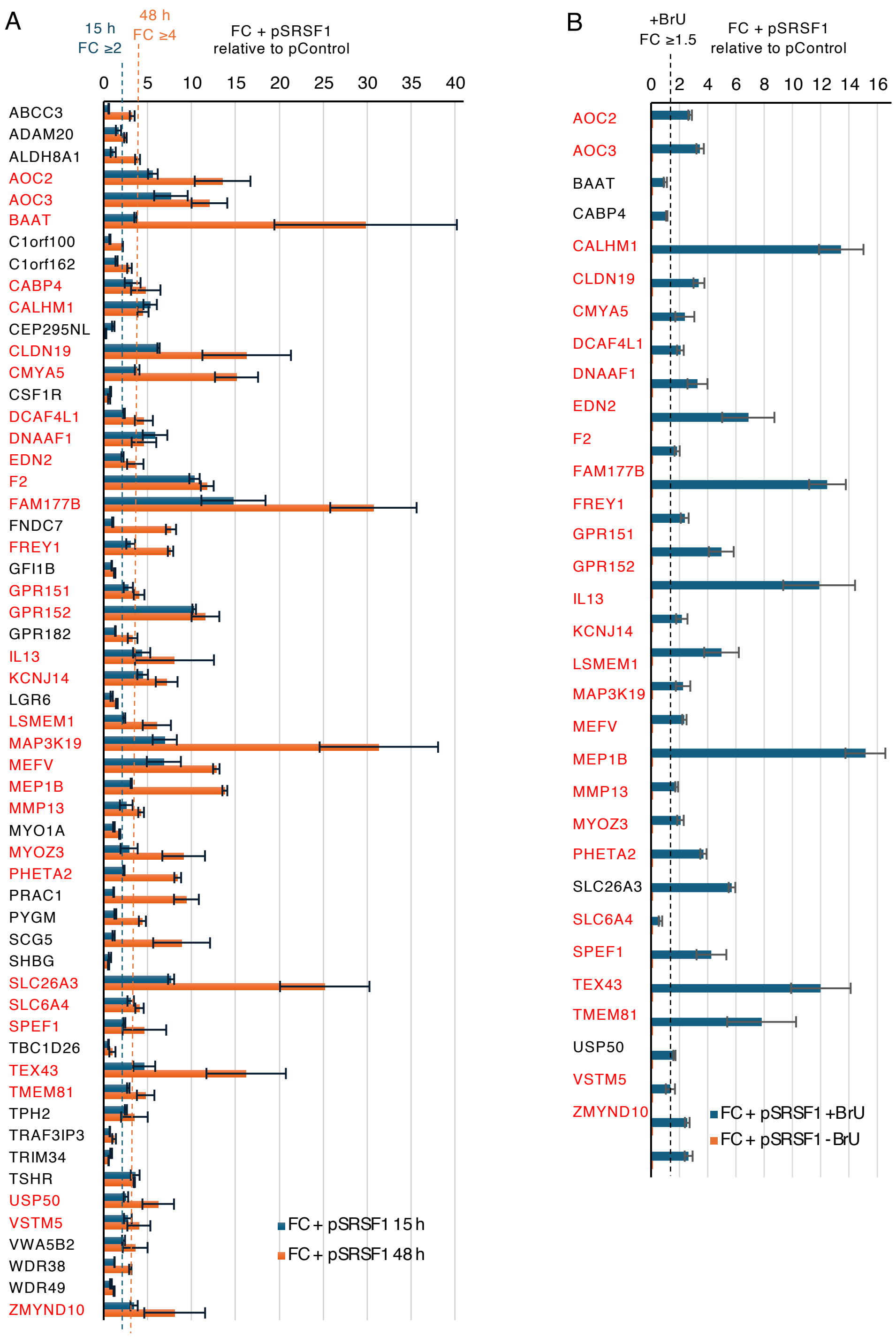
