## Supplementary Figure S5 for "SRSF1 coordinates transcription and alternative splicing to modulate p53 and apoptosis"

**SRSF1 overexpression produces similar decay profiles for selected transcriptionally upregulated transcripts.** HEK293 cells were transfected with control or SRSF1 expression vectors and treated with actinomycin D to inhibit transcription. RNA levels for selected SRSF1-regulated targets were measured by RT-qPCR at 0, 3, 6, and 9 h and expressed relative to the corresponding 0 h value. SRSF1 overexpression did not appreciably alter the decay kinetics of AOC2, AOC3, TEX43, MEFV, MYOZ3, or RPL13A, arguing against increased RNA stability as the basis for the higher RNA levels of the SRSF1-responsive transcripts tested. Data represent mean ± SEM from three independent biological replicates.

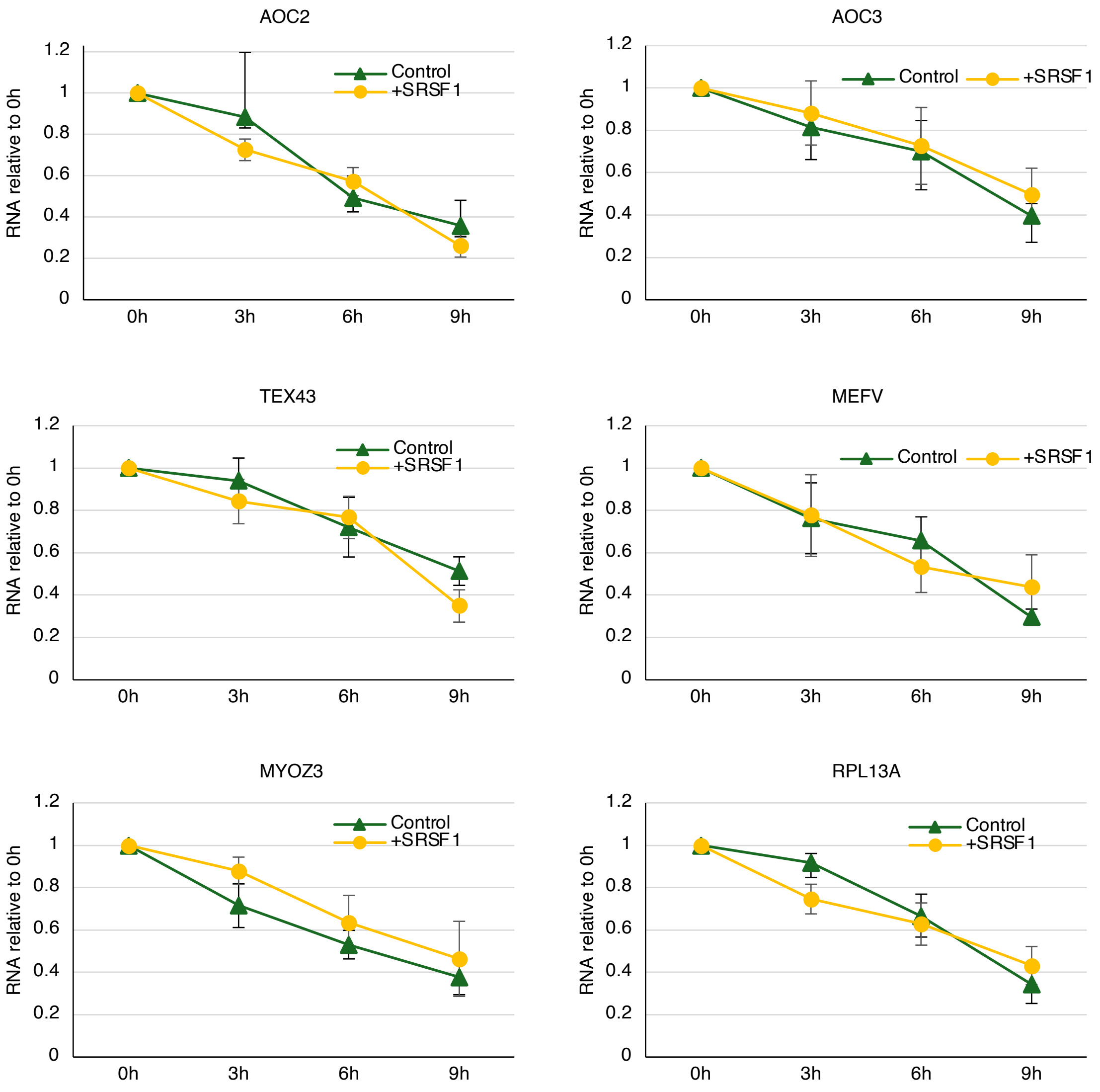
