## Supplementary Figure S6 for "SRSF1 coordinates transcription and alternative splicing to modulate p53 and apoptosis"

**SRSF1 and transcriptional-target associations with cancer subtype and clinical outcomes.** Analyses used TCGA data. The 28 transcriptional targets comprise 18 with conserved responses across HEK293, MCF-7, and PC-3 cells and ten additional regulated genes. **(A)** Adjusted partial correlations between SRSF1 and target-gene expression within breast cancer (BRCA) subtypes; sample sizes are indicated. **(B)** Associations between target expression and thyroid differentiation score in thyroid carcinoma (THCA), adjusted for clinical variables, sample composition, and junction-library size, with additional SRSF1 adjustment in the second column. Colors indicate adjusted partial correlations. **(C)** Selected associations with overall survival (OS) and progression-free interval (PFI) in clear-cell renal carcinoma (KIRC). Cox models adjusted for age and sex and stratified baseline hazards by stage. Squares and lines indicate hazard ratios (HRs) and 95% confidence intervals (CIs) per standard-deviation (SD) increase in $\log_2$(TPM + 1) expression. **(D)** Six leading nominal PFI associations in pancreatic adenocarcinoma (PAAD), adjusted for age, sex, and stage; none passed FDR correction. **(E)** SRSF1 expression across BRCA subtypes, association with thyroid differentiation in THCA, and OS/PFI analyses in KIRC. BRCA comparisons show basal-like minus comparator expression differences in SD units, adjusted for age, stage, composition, and junction-library size. The omnibus *P* value tests overall subtype differences; pairwise q values reflect correction across all six comparisons. THCA shows adjusted correlation coefficients and KIRC shows HRs and 95% CIs. q denotes Benjamini–Hochberg FDR-adjusted *P* values; KIRC retained the original testing denominator. Thyroid differentiation score (TDS) Heatmap symbols: *q < 0.05, **q < 0.0005, ***q < 0.000005; no symbol indicates q ≥ 0.05. Bold q values indicate q < 0.05. [L] indicates expression ≥1 TPM in fewer than 10% of samples.

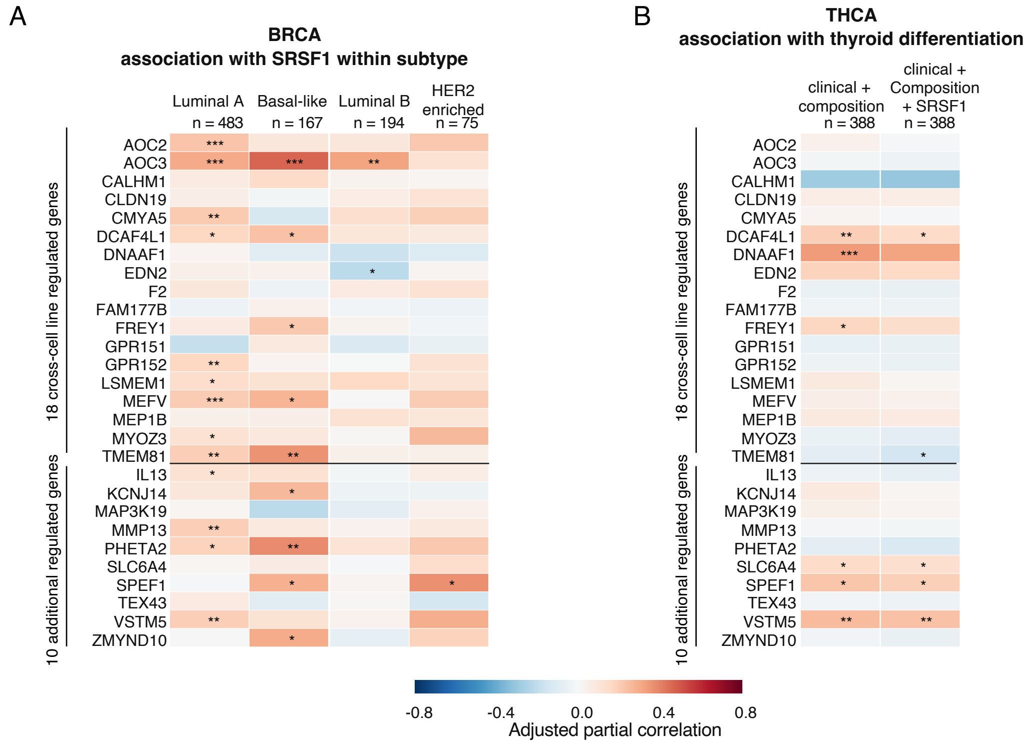


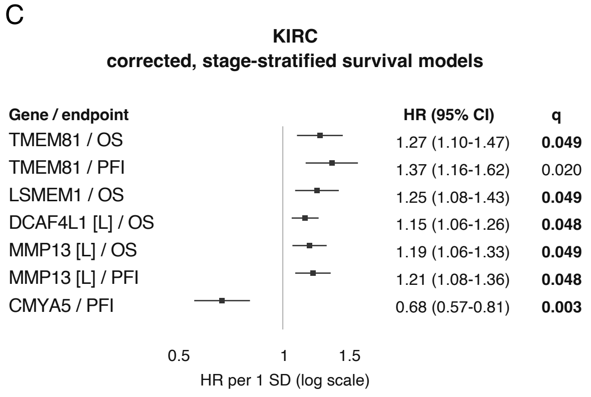


D

**PAAD**
**leading nominal PFI associations**

| Gene | HR (95% CI) | P | q |
|---|---|---|---|
| SLC6A4 | 0.70 (0.53-0.92) | 0.010 | 0.127 |
| F2 [L] | 0.71 (0.54-0.94) | 0.015 | 0.158 |
| AOC2 [L] | 0.79 (0.64-0.98) | 0.032 | 0.234 |
| FAM177B | 0.79 (0.64-0.98) | 0.033 | 0.234 |
| SPEF1 | 0.79 (0.63-0.98) | 0.035 | 0.234 |
| TEX43 [L] | 0.62 (0.39-1.00) | 0.048 | 0.280 |

E

**BRCA SRSF1 expression**

Basal-like vs Luminal B –0.732 SD; q = $1.67\times10^{-8}$
Basal-like vs HER2-enriched –0.396 SD; q = 0.00110
Basal-like vs Luminal A –0.283 SD; q = 0.0269
Subtype omnibus P = $1.52\times10^{-13}$

**THCA Differentiation**

SRSF1 vs TDS n = 388
Adjusted r = +0.139
P = 0.027; q = 0.079

**KIRC SRSF1 survival models**

OS: HR 1.025 (0.874–1.203) P = 0.759; q = 1.00
PFI: HR 0.939 (0.803–1.099) P = 0.435; q = 1.00
