## Supplementary Table S1 for "SRSF1 coordinates transcription and alternative splicing to modulate p53 and apoptosis"

| **Supplementary Table S1.** | |  |
| --- | --- | --- |
| **Primer pairs used for RT-PCR and RT-qPCR.** | | |
| Primer sets for RT-PCR | |  |
| **Gene** | **Forward Primer** | **Reverse Primer** |
| ACER2 | ACATGCGTGTGTTTAAGCTG | GATGACAGGGCCTTGCTCAG |
| ANKLE2 | CAGACGGCTGAGGCCTCTCAC | CAGAGCCCATTTCTGCTGCTG |
| ANXA11 | GAGAGCCACTCCCGCCCACG | CAAATGTCCCAGACATGTTG |
| ATR | CTCTTCAGAAACCAAAGAAG | GTAGGATCAGGGAATGTTCTC |
| ATXN3 | CAAGAAGGCTCACTTTGTGC | GTTTTCCTAATTTTCTAACTG |
| ATXN7L3 | GGAACTTAGCAATTCGGATC | TCAGTCATGTCAAAGCTGTC |
| BANP | GCGGAGAACGCCCAACTCGTC | ACGTGATGGATCTGGAGGTTG |
| BCL7B | GTTCCTGTGACAGACAGCAAG | GATCTCCTGGCCCAGCGTTG |
| BIRC5 | CTGTCAAGAAGCAGTTTGAAG | CTTTCCTAAGACATTGCTAAG |
| BIRC6 | AGATACAAGTGACACTGCTG | TCATTCACTGCCTTGACATG |
| BORA | GATGTCAAGGAATCAAAGATG | CTGGAATGACATTGTGAAAG |
| BRCA2 | CGTCTATTTGTCAGACGAATG | GAAGCTTGTTTCCTGTACCAG |
| BRD1 | CCATTAGCACCGCCTCGGC | CTTCTCATTGACGATCTCCAG |
| BRD8 | TTGATTATGCAAACATCCGAG | GCAAGCCCAAGCTCTATCTTG |
| BRIP1 | ACCAGCAGATGAGGGCGTAAG | AGGAGTTTATCTTTTCCAGTGG |
| CALM3 | AGCTGACTGAGGAGCAGATTG | CACTTTGCAGTCATCATCTGT |
| CCDC57 | GCATGAGGAGCTCGACCGTC | ACCTGTGCCTTGAGCTTCAC |
| CCNK Ex 1b | CAAGGAGAGACGTCGCTGAG | CAGTTGCCAGGGTATCATAGTG |
| CCNK EX 4 | CAGTAGACATTGAAACCAGAG | CAATGTCTTTATGTGCGTTTG |
| CDCA7 | ACTGATTCCAACTCCGATTC | CTCACCAACATGTACTTATC |
| CENPU | ATCTAAAGTTTGTAAGGCAG | CTGCATTCCTAAGGGAAGAC |
| CEP295 | CTTGAACAAATTGAACAGCA | GACTCAACTTGAGTCTCTGAC |
| CEP57 | TTGCAGACTGGTCTAGAAAC | GTGTTACTGACAACTTCTTAC |
| CEP120 | ACAGTTGGCTACTGATCCTG | CTACACTGCTACGGATGCGAAC |
| CEP350 | AGTAGCCCCTACAAGTACAG | GTTAGGCATTTCTTCCTCAG |
| CFL1 | TACGCCACCTTTGTCAAGATG | CTGCCAGATGCTCCAGGCAG |
| CHEK1 Ex 3 | AGTGGTGGGCAAAGGACAGTC | TCTGGTTTAATATCCCTGTGAG |
| CHEK1 Ex 8 | CAACCCAGTGACAGCTGTCAG | TAGAAGCACTGTTTACTGGAG |
| CHEK2 | AGAGCTGTTTGACAAAGTGGTG | ACTCCAGCAGTCCACAGCAC |
| CNOT2 | TGACACGACCCTTGAGTGAC | ACATAGAAGACTGGCTGTAG |
| CPSF4 | GACCACATCAAGTTTGACTTG | AACTTGGAGTAGAAGTAGCAC |
| CREBBP | CTTATTAGCAGAGAAAATCTAC | TCTGGACAGAGTGGTTCATTG |
| CSTF3 | AGACAGTAATGAAAGGCTTG | CTCTTCATAATCTGCATATGC |
| CUL1 | CTTGCATTGGTGACTTGGAGAG | CTTTTTCATATATTCAGTAAC |
| CUL7 | TCCTGAAGCAGCTGGTGGAC | CTTGGAGATAGCGTCTCTTG |
| DBF4 | AACAGATGGCGATAAGTATG | GTTTCCTGGGTCTCTTTATAC |
| DDX3X | GATTTCCAGCAAAGTGTTAC | CATATGTAGGCAGCTTCATC |
| DICER1 | TCCAGAGTGTTTGAGGGATAG | GATAACGGCATCTTGGTAATC |
| DTL | AGCAAAGTGTTACTGTGGTC | TATGTAGGCAGCTTCATCAC |
| ELOF1 | CTGAACAGCTGGCTTGTCTTG | TTCACATCACAGGATTTCTC |
| ESCO2 | GTTCTGCCACATGATCCAAG | TCAGAAAATGCTATTTCATC |
| EXO1 | ACCTCGAGTGCCACATGCGAC | GTGAAGCCAGCAATATGTATC |
| FGFR1 | TGACCTGCTGCAGCTTCGCTG | GATGTCCAATATGGAGCTAC |
| FGFR1OP2 | GAGTCCGCATCGCCACAGCTG | CTTGTTACGATGTACCATGTC |
| GAS8 | GTCGATGGGCTCGCTCCAGAG | CAGCCTTCATCTCTGTCAGG |
| GBF1 | CTACTGCCAGCTATGATCTTG | TAGCCTTGGGCAGTAGTTG |
| GIGYF2 | GACCTCATCATCATCTGCTAG | CTTCATCATCAGGTTCTGTG |
| GSK3B | TGAATGTGCACAAGCTTCCAG | CTGGGAACTCCAACAAGGGAG |
| HERC5 | CTGAAGAGGACAATTCCTAC | GAAGTCTTTTGAGCAGATTATC |
| HPF1 | CAAGCCTTGGACTTCAATTAG | ATCTCTCTGTTTCATCTTCAC |
| INTS6 | GAGTTGGAACCTTCACCACTG | TGTCTGCTATTAGGTTAGGTG |
| JMJD1C | GTGAGAGTATATAGACAAGAC | CTTCATCTGGTATACTGCTATC |
| KIAA0753 | AGTAGCCGCGCCTCACCCTAG | CTGCATCTTTACAATGGTATG |
| MBTD1 | CTCAGTATCAAGCTACCTTG | CAATTCCAGCAATCCAGAAG |
| MCM3 | GAAGACTCGCCAGCCAGATG | GTCATCCTGCATCTTGCTCAG |
| MCM8 | GTAAGCATGGAATGTTGATG | GAGTTACCGTCAGACCAGAG |
| MDM1 | TGTAACAAACCATACACCAG | CAGTTTCATGGATGACTGAG |
| MDM4 | GAAGACTTAATTGAAAATTTAG | CTCAGAGGTAACCTCTACATC |
| MED13 | GTTCTGCTAGAAAATTTGGTC | TTCAGTGGTATAAGTAGCTG |
| MITD1 | CAAAGATAATACTAAGAGATG | CTACTTTGCTGCACTTGCTC |
| MOK | GATGGAAACTACTATGCATG | GATGTATTCCGTGTACGGCTG |
| MMS22L | GAGTATGCATCAACAATTGTG | GACCAACTGTTTCATGTACTC |
| MTF2 | TATAGCACACCTATGCCTTTAC | GAATCAGATATAGGTTTGGATG |
| NABP2 | CGCTTCAGTTTTCAAAGGTTG | GCACTCAGTCCATTCCCATTC |
| NCAPD3 Ex 6 | CTAAGAGCTCTCAGGCTAAC | GAGCACAGCAAATACAATCC |
| NCAPD3 Ex 19 | CTTAGGACATATTCTCTGTG | ACAGGTCTTCGTCCATATTC |
| NCAPD3 Ex 32 | CATTCGACTCTTGCTCCAAAC | CAGGCTGGAGTGTCTTTATTC |
| NFYA | GTGTACCTCACAGCCTTCTAG | TGGTTTGACCTTGTCCACCAG |
| NKX2-5 | CTATCCACGTGCCTACAGCGAC | CTTGACCTGCGTGGACGTGAG |
| NME6 | CTGCAGCCGGAGTTCAAACC | GGTCCTTCAGGTCAACGTAG |
| NR2C2AP | GTCTTGTGGAACCGACATGAC | CTGAAGCGAGTTGTTGTCCTC |
| NSD2 | ACTGAAGTATCTGTCTCATC | GAAGGGTTGGAAGCATGGCAG |
| NUP62 | TGGATGTCCGTCTTTTCTCTG | TGTTCCAAAAGTGAAGCCTGTC |
| OARD1 | CTGTATCAGTGAGGATTGTC | GTCAGTGACTCCATTCTTCAG |
| OFD1 | CATGAGGAGTTCGAAAGCTG | GCAACCATACGTGCTAGAAC |
| PAXIP1 | ACCAACTGGTTTAAAGTATG | GGACATTAGCTACTTCATTC |
| PAXX | GAGTGCGGCTGAGGACATC | GAAGTCCACGCCACCAGCTG |
| PCM1 | ACGATTGGATCAGCACAGTG | CTGTAGAGCTCTTAGTTGCTC |
| PLK4 | TCTGAACAGAGCAAGACTAG | CTTACAAGCTGTACAAACCTG |
| POLQ | CAGATCTAGCCAGTTCTCTG | CTGTAATACAGAGTCCAGTC |
| POU2F1 | GCGGACGGAGGAGCAGCGAGTC | AGCCTGCGCCTGGGCTGTTGAG |
| PPP4R2 | GACTATGTGAATTGTTAACAG | GAGGTCGAAGAGTCACTGTG |
| PPP6R3 | GAGACTGCAGAGGCGAAGTG | CTACAAGTGAAACACTAGAC |
| PRIM1 | CATACGCATCATTGACAGAG | TCGAGAATATCTTGATTAAC |
| PRKAB2 | CATAATGACTTTGTTGCCATC | AGTGTCTTTGTTAAGAATAAC |
| PSMD11 | GAAGCATTAAAATGCGTGGCTC | GGCTTTGTTGTAGAGGGAATC |
| PUM1 | CTGACCAGCACCAATGCCAAG | GAGCAGCGCTGATGATGTATG |
| RABGGTB | ACGCTGTATGACAGTATTAATG | TGGGATTCAGAACCTGGTCTG |
| RAD51D | GTGGACCTGGTTTCTGCAGAC | ATCCTGGGTTTTAGCCTGAAG |
| RAD54L | CTCATGACCAGCTGAAGCTTG | TCCTTAGATCCTCCATCGATG |
| RAN Ex 2 | TGCGCTTCCGCCATCTTTCCAG | CTCTGTTGGTGTGGAACACTAG |
| RAN Ex 3 | CCATCTTTCCAGCCTCAGTC | CTTGGATATAATAGCCATCTC |
| RANGAP1 | TGACCCGGACCCTAAATCCGTC | CAAGCTGTCAAAGTCTTCAATC |
| RB1CC1 | CTCAAGCAAAGCAAACATAAG | CGTCCCAGCACTGTAGGTAC |
| REV3L | CTCATCACCAGTAGAGGAGCTG | CTGTAAGTTTTGTATGCTGAC |
| RFC5 | CTGGTCACTGTGCAGGCGCTTG | AGCTGTTTCGCACAGGCTAG |
| RFFL | CAAGGGCTGCTGTGAGAAGTG | TCATTCATGCGCTTGCCACAC |
| RIF1 | GTATCCAGTATAATTGATTC | ATAGCTTCTGAAGTTCTGAG |
| RNPS1 | TCGGCGGCTGAGGAAAGCAG | CTTTGACTCCTAGCAAGCTC |
| SAP30BP | GACAAGATCCAGAAGCTTTATG | CTGGCAGTGGTAGTGGTGGTG |
| SLF1 | CAAAGCATATCATCCAGATG | CAACTTTCCATCTGTGGAAG |
| SKP2 | ATAGTGTCATGCTAAAGAATG | AGTGTCAGTCGGCATTTGATG |
| SMAD6 | TCTGTCCGATTCCACATTGTC | TCGCTGCGCTGCTCCAGGTTG |
| SRPK2 | CATCATAGAGCTGCTAGGCAG | CTACATGCAACATAGGCTAC |
| SRRM1 | ATGAGCAAAGTAAATTTGGAG | GATGTTTTCTTGTGCACTTAG |
| SRSF5 | GGTTACTATTGAACATGCTAG | CTGTGCCTTTTGCTGCCTTC |
| SRSF7 | GTCGAGGTCAAGATCAAGATC | CATCATATGGTCTGAAAATGT |
| SSX2IP Ex 6 | GAACAGGAATTTGCATCAGC | GAAGAACCTTCTTAAGTTCTG |
| TAF1D | CTGAAGTATGAACACCACCTG | ACATTTTCAGGCCTCTCTGTC |
| TAF2 | CATTGCTGAAACATACCACATC | TCTTTAATCCAATGGCGGTAC |
| TAF11 | AGGAGCAGCTGAACCGTTATG | AGGCCTTTCTAGACTTTGGTC |
| TCF7 | AGCACCAAGAATCCACCACAG | GTTGGGAGGCTGAGAGCAGTG |
| TFAM | GCTAGTGGCGGGCATGATAAC | TCTTTGAATCAGGAAGTTCC |
| TFIP11 | CAACCGACAGCGCCACTGGCAG | ACTCTTCCACGGTCTTGTAAG |
| TICRR | CTGCCAGTGCTATGATCCTC | GACTCCATCAAATCGGAACTG |
| TOP3B | TGGATCCGCGGTGGGCTGTG | CATCTTGAAGCGCACTGGCTG |
| TPR | GTATGTCGCTGTGAAGATCTG | CTTCATCAGTTCTTCATGCTG |
| TTI2 | CATGGACTCTGCAACAGGTC | CACAATGGGTGGTGGGTCGAG |
| UBE2I | CTCCACCTGTCCGCTACGCTC | TGGAATGGCGCACTCCCAGTTC |
| USP47 | TCAAGACGGTGGAAGCCTTC | CAGTCTTCTGGAGTCGACTG |
| WRN | GATGATGAAAATGAAGCTAATG | TTAAGCTGTAGCACTTGGTC |
| XPO5 | CGACTGGCAGAGGATGTAGTG | AGTGATGTGACTCATAGACAC |
| ZNF33A | GTGTTCACAAACCAGAGGTG | ACATCACCTTGTTCCTTAGTC |
| ZNF43 | ACTTTGTCGCAGCCTCCGTTAC | GCAAGGTCCAGGCAAAAGATG |
| ZNF45 | CTTCCTAGCGGTGTAACGTTG | AATGTCCATGAAACAGAGGTC |
| ZNF195 | GGCTCTGTGTCCCCTGCTAG | CAACGGAGAACAAGTTTCTG |
| ZNF207 | CAGCCTGTTCAACCTCAGCAAG | CATTCCAGGCATCAGAGGTG |
| ZNF248 | CTGCTGGCTCTGACCGCCCTC | GGATTTGTTCATTTTCCGCTC |
| ZNF317 | ACCCAGGATTCCACCTGTCTC | AGTGAAGTGAGGTTGCTGTAG |
| ZNF324 | GTCACTTGGCTGCTCGCGTCAG | AGTGAGGTCACAAGTGTGAAG |
| ZNF468 | GATTGACTTCTAAAGACTGTTG | GAGACGAGGTTCCTATAATTC |
| ZNF484 | AAGTGGGTCCTGTTCTGGTTC | CACTGAGATCAAGTTGAAATAG |
| ZNF528 | AGTGTTACCTCCATGTTAGAG | GTGAAACTGAGGAATTGTCAC |
| ZNF548 | CTATGGCTCTGTCTGACGTC | GTCCATTTCTGCCATCGCCAG |
| ZNF600 | GACCTGGAAATTCTCGCCTG | GTTTCCGGAAGTACGTAGTCAC |
| ZNF616 | GAGTCAGTGATGGCTACTCAG | ATCATAGCTTTGATGTCTTTC |
| ZNF649 | GTCATCATTTCTGGGAACACTC | TGGACTGTGGATTTCATCTTC |
| ZNF669 | TTCGCCGGAGCCGGCTCTTC | AGATCAAGATTTGGAATTTG |
| ZNF761 | AAGCGGATAGCGTGGAGTGAC | GATGCTGAGGCACAAGAATCAC |
| ZNF772 | GTGTCTTCTTCCCTGAAGAGA | TGATCTGTGACATTCCTATAG |
| ZNRD2 | CAACGGCAACATGGCCCTGAAC | GATTATCTTTATCCACGTCTG |
| Primer sets for RT-qPCR | |  |
| **Gene** | **Forward Primer** | **Reverse Primer** |
| ABCC3 | ACTCCAAGACAGAGACAGAG | GGTAGCGCACAGAATAATTC |
| ADAM20 | TGTATTCAACCGCCTCCATATC | ATGGTTCCACAGTCACATTC |
| ALDH8A1 | AACCAGGCAGGCTACTTTATG | CACACGTCACTGGACCAAATA |
| AOC2 | TGCTGAGAGCTGAGTTTACA | CAGGGTAGAGCCATTGTAGT |
| AOC3 | GATCTCTACTCGCACTACTT | GAAGACCGTATCCCACACAT |
| BAAT | GCTGAAGAGACATGGGAAGAAC | GCACAGCACAGAGGAGAATAG |
| C1orf100 | GAGCTCCCAGATTGGAACCAG | CTCTTCACAACAGTAGAGCTCTG |
| C1orf162 | GCAATGGCTCCACATGTAAAC | TTGTGGTGGTTAGAGAGACAG |
| CABP4 | AGTTTGAGACCAGCCTAACA | TCACCTCAGCCTCCTGAATA |
| CALHM1 | ACTGGGTCAAATGGTGTCTC | GCCTCATCCTCATTGTATAG |
| CEP295NL | CTTAGTGTTCCCTCCCTCAAAG | CTTCCTAGGCTGGACTTCTTAC |
| CLDN19 | TTAGGAGCTTGCAGGCATTAG | TCAGGAGAGGAGGAATGGTAG |
| CMYA5 | CCTGGCCTTCCAAAGAAGAT | GACTTCACCTGTTCCACTGA |
| CSF1R | TGCTGACTGTTGAGACCTTAG | CTCCTGCAGAGATGGGTATG |
| DCAF4L1 | GCAGAAGAGTAGTGACCAGA | TCGGACACAGTGCCATTACT |
| DNAAF1 | AGTGACCCTGAACTGGACTA | CCGCCTTTGAGGTGTCTTTA |
| EDN2 | AGGGACATTTCCACAGTCAAG | TCCAGCTCACGACACTATCTC |
| F2 | GATGGTGGAGATGGACAGGA | CATGAAGAAATCAGCGGAGA |
| FAM177B | GACGAGGGTACAGTAAAGAAG | CAGGATGGTTGTTGGAAGTAC |
| FNDC7 | CAGTTTAGAAGCCGGAACTC | GAGGACTTGTTCTCTGATTG |
| FREY1 | TCTTCCTCCACCTCATCCTG | GAAAGTGGCTGCGAGAGTTC |
| GFI1B | CCGTGTGCAGGAAGATGAAC | AGGCACTGGTTTGGGAATAG |
| GPR151 | GACAAGGTTCCATCTCCAGA | GGAATCTCTGCCTTCTCAGT |
| GPR152 | GGACCATTCTGTCAGCCTATG | GAGCAGGTAGCCAGAGTAGAC |
| GPR182 | CACCTGCTCTACTTCTTCTATG | GAAGTGTGGGCTGAGAAAGT |
| IL13 | CACGGTCATTGCTCTCACTT | CTTCTGGTTCTGGGTGATGTT |
| KCNJ14 | GTGAGAAGCCAATGGTATAG | GTGAATCCAGACCTACAAAC |
| LGR6 | GGGAAGTAAAGACAGTGAAG | ATCCTATCCATGAGTCACAC |
| LSMEM1 | AGATATCTGGGAAAGCCTGAG | GTCTAATCACCTGAGCAGAATC |
| MAP3K19 | GTTGTAGCAGCATGGAGACA | GGAAGGTGATGTGGATAAGA |
| MEFV | CTGCTTCTGGCCTTGTATCT | CCGTGGGCACAGTAACTATT |
| MEP1B | TGGTGGTACATGGGAGAAAG | ATCAGCATCAAGGCAAACAC |
| MMP13 | GCATCTGGAGTAACCGTATTG | CCCGCACTTCTGGAAGTATT |
| MYO1A | TCACCTTGGCAGATTTCATC | CTGGCCATGTCTTGTCTAAG |
| MYOZ3 | GGATTTGAGTGGGACCTTGT | CCCGTGCTCAGTTCCATATT |
| PHETA2 | TCTCAGTCCTGGGTATTCTAG | TCTCGAGCAGCTTCAGTTTAC |
| PRAC1 | GATGTTGTGCGCCCATTTCT | TCTCGCCCAGTAGATGTTTC |
| PYGM | GGAGAACGTGACTGAGCTGA | GGCCAGAGCAAAGTAGTAGT |
| SCG5 | GTGAGAGGCACACTTATCT | CCTCCTCCGCATCTTAGCT |
| SHBG | GATTCCTGGCTGGACAAACA | AGTCCCTGGAGGGAGAAATA |
| RPL13A | AGCCAGAAGACTGATTGGAG | AGTGCTTGCACATTCTAACAG |
| SLC26A3 | GTGTATCCTACGGCTGTGAC | CTCCACGTCAGGTGTAATAG |
| SLC6A4 | GTAAACTGCGTAGGAGAGAAC | CACAGCAGCACATGGATTAG |
| SPEF1 | TAGCGATGGAGTCCTTGTTG | GCTTCTGCTGGAGAGAGTTG |
| SRSF1 | TGCCTACATCCGGGTTAAAGT | TACGGCTTCTGCTACGACTA |
| TBC1D26 | AGGAACCTTCAAACCTCTATG | CAGGCTATGGTATAGCAATG |
| TEX43 | ATCCATACTGGCCCTTTAGA | TCTGGAGAGAGGCGAATGCA |
| TMEM81 | CCCTGAGAAGCTGCAAGAAG | AGACGGTCTCCTCCTTATAG |
| TPH2 | CTGACCACCCAGGATTTAAG | CTGGGAATGGGCTGACCATA |
| TRAF3IP3 | AGTCCAAAGTCCTTCCCTAAC | CCTGGAGCTGCAAAGTCTTC |
| TRIM34 | GTCCTGACTCATGCAACTTC | CTGGCTTCTTCTGGTCCTAAC |
| TSHR | AGCACCTGAAGGAACTGATAG | CAGTGGCTTGGGTAAGAAAG |
| USP50 | ACTCACTAACTTGGACCTCAC | TAGTGGCCACCATCCAAATC |
| VSTM5 | TCAAAGCCACAAGGACAGAG | CGTCACGGTGATGACATAGTAG |
| VWA5B2 | CACCCAACACACTCAAGTCAC | CTCTAGGGAGGGAGTAAGAAG |
| WDR38 | GCCTGCTTATCCAACTGAAG | CTTTGACCATGCGGGAATAG |
| WDR49 | GCCACTCAGCCAGTGTAATAC | CTGGTGTTGAATATCCCAGAG |
| ZMYND10 | AAGGTGTTCCCTGTGTTCTG | GTCTCCAAGAGGTTGATGATG |
| 7SK RNA | CAAGGTCCATTTGTAGGAGA | AGGCAGACTGCCACATGCAG |
