## Supplementary Table S3 for "SRSF1 coordinates transcription and alternative splicing to modulate p53 and apoptosis"

**SRSF1-regulated alternative-splicing events validated in HEK293 cells and across cancer cell models.** The table lists SRSF1-regulated alternative-splicing events validated by RT-PCR in HEK293 cells and includes gene annotation, GO/Reactome functional category, validation in MCF-7 and PC-3 cells, event type, GENCODE annotation status, and the affected exon or intronic region.

| **gene symbol** | **gene definition** | **GO/REAC: cell cycle** | **GO/REAC: regulation of p53 activity** | **GO/REAC: DNA repair** | **GO/REAC: transcription regulation** | **AS HEK293** | **AS MCF-7** | **AS PC3** | **AS type** | **AS Gencode** | **AS exon** | **exon Gencode** |
| --- | --- | --- | --- | --- | --- | --- | --- | --- | --- | --- | --- | --- |
| ACER2 | alkaline ceramidase 2 |  |  |  |  | Yes | No | No | IE | novel | ex 5b | novel |
| ANXA11 | annexin A11 |  |  |  |  | Yes | Yes | No | IE | known | ex 1b | known |
| ATR | ATR serine/threonine kinase |  |  |  |  | Yes | Yes | Yes | SE | known | ex 42 | known |
| BIRC5 | baculoviral IAP repeat containing 5 |  |  |  |  | Yes | Yes | No | IE | novel | ex 3b | novel |
| BIRC6 | baculoviral IAP repeat containing 6 |  |  |  |  | Yes | No | No | IE | novel | ex 14 | known |
| BRD8 | bromodomain containing 8 |  |  |  |  | Yes | Yes | No | IE | novel | ex 21b | novel |
| BRIP1 | BRCA1 interacting helicase 1 |  |  |  |  | Yes | Yes | Yes | SE | known | ex 4 | known |
| CCNK | cyclin K |  |  |  |  | Yes | Yes | NA | SE | novel | ex 4 | known |
| CEP57 | centrosomal protein 57 |  |  |  |  | Yes | Yes | No | SE | known | ex 8 | known |
| CHEK1 | checkpoint kinase 1 |  |  |  |  | Yes | Yes | Yes | SE | known | ex 3 | known |
| CHEK1 | checkpoint kinase 1 |  |  |  |  | Yes | Yes | Yes | SE | novel | ex 8 | known |
| CHEK2 | checkpoint kinase 2 |  |  |  |  | Yes | Yes | Yes | SE | known | ex 10 | known |
| CREBBP | CREB binding protein |  |  |  |  | Yes | Yes | Yes | SE | novel | ex 11 | known |
| CSTF3 | cleavage stimulation factor subunit 3 |  |  |  |  | Yes | Yes | Yes | SE | known | ex 11 | known |
| CUL7 | cullin 7 |  |  |  |  | Yes | Yes | NA | IE | novel | ex 9b | novel |
| DDX3X | DEAD-box helicase 3 X-linked |  |  |  |  | Yes | Yes | NA | SE | known | ex 9 | known |
| DICER1 | dicer 1, ribonuclease III |  |  |  |  | Yes | Yes | Yes | SE | novel | ex 16 | known |
| ESCO2 | establishment of sister chromatid cohesion N-acetyltransferase 2 |  |  |  |  | Yes | Yes | No | SE | known | ex 9 | known |
| EXO1 | exonuclease 1 |  |  |  |  | Yes | Yes | Yes | SE | known | ex 3 | known |
| FGFR1 | fibroblast growth factor receptor 1 |  |  |  |  | Yes | No | No | IE | novel | ex 3b | novel |
| FGFR1OP2 | FGFR1 oncogene partner 2 |  |  |  |  | Yes | No | No | SE | novel | ex 2 | known |
| GBF1 | golgi brefeldin A resistant guanine nucleotide exchange factor 1 |  |  |  |  | Yes | NA | Yes | SE | novel | ex 23 | known |
| GSK3B | glycogen synthase kinase 3 beta |  |  |  |  | Yes | Yes | Yes | IE | novel | ex 8b | novel |
| HERC5 | HECT and RLD domain containing E3 ubiquitin protein ligase 5 |  |  |  |  | Yes | No | Yes | SE | novel | ex 10 | known |
| INTS6 | integrator complex subunit 6 |  |  |  |  | Yes | Yes | Yes | SE | novel | ex 10 | known |
| JMJD1C | jumonji domain containing 1C |  |  |  |  | Yes | Yes | Yes | SE | novel | ex 6 | known |
| MBTD1 | mbt domain containing 1 |  |  |  |  | Yes | No | No | IE | novel | ex 6b | novel |
| MCM3 | minichromosome maintenance complex component 3 |  |  |  |  | Yes | Yes | Yes | IE | novel | ex 16b | novel |
| MDM4 | MDM4 regulator of p53 |  |  |  |  | Yes | Yes | Yes | SE | known | ex 9 | known |
| MED13 | mediator complex subunit 13 |  |  |  |  | Yes | Yes | Yes | SE | known | ex 25 | known |
| MMS22L | MMS22 like, DNA repair protein |  |  |  |  | Yes | Yes | No | SE | novel | ex 16b | known |
| NABP2 | nucleic acid binding protein 2 |  |  |  |  | Yes | Yes | Yes | RI | novel | int 5 | known |
| NCAPD3 | non-SMC condensin II complex subunit D3 |  |  |  |  | Yes | NA | NA | SE | novel | ex 5+6 | known |
| NCAPD3 | non-SMC condensin II complex subunit D3 |  |  |  |  | Yes | NA | NA | IE | novel | ex 19b | novel |
| NFYA | nuclear transcription factor Y subunit alpha |  |  |  |  | Yes | Yes | Yes | SE | novel | ex 3 | known |
| NR2C2AP | nuclear receptor 2C2 associated protein |  |  |  |  | Yes | Yes | Yes | SE | known | ex 3 | known |
| OFD1 | OFD1 centriole and centriolar satellite protein |  |  |  |  | Yes | NA | Yes | A3SS | novel | ex 15 | known |
| PLK4 | polo like kinase 4 |  |  |  |  | Yes | NA | NA | SE | novel | ex 9 | known |
| POLQ | DNA polymerase theta |  |  |  |  | Yes | NA | NA | SE | novel | ex 7 | known |
| PRKAB2 | protein kinase AMP-activated non-catalytic subunit beta 2 |  |  |  |  | Yes | Yes | Yes | SE | novel | ex 5 | known |
| RAN | RAN, member RAS oncogene family |  |  |  |  | Yes | Yes | Yes | SE | known | ex 2 | known |
| RAN | RAN, member RAS oncogene family |  |  |  |  | Yes | Yes | Yes | SE | known | ex 3 | known |
| RB1CC1 | RB1 inducible coiled-coil 1 |  |  |  |  | Yes | No | No | IE | novel | ex 2b | novel |
| REV3L | REV3 like, DNA directed polymerase zeta catalytic subunit |  |  |  |  | Yes | Yes | Yes | SE | known | ex 15 | known |
| SLF1 | SMC5-SMC6 complex localization factor 1 |  |  |  |  | Yes | Yes | Yes | SE | novel | ex 3 | known |
| SRRM1 | serine and arginine repetitive matrix 1 |  |  |  |  | Yes | No | No | IE | novel | ex 4b | novel |
| SRSF7 | serine and arginine rich splicing factor 7 |  |  |  |  | Yes | NA | NA | SE | known | ex 7 | known |
| SSX2IP | SSX family member 2 interacting protein |  |  |  |  | Yes | No | Yes | SE | known | ex 6 | known |
| SSX2IP | SSX family member 2 interacting protein |  |  |  |  | Yes | Yes | Yes | SE | novel | ex 12 | known |
| TAF11 | TATA-box binding protein associated factor 11 |  |  |  |  | Yes | Yes | Yes | SE | known | ex 4 | known |
| TAF1D | TATA-box binding protein associated factor, RNA polymerase I subunit D |  |  |  |  | Yes | Yes | Yes | SE | novel | ex 5 | known |
| TAF2 | TATA-box binding protein associated factor 2 |  |  |  |  | Yes | Yes | Yes | SE | known | ex 8 | known |
| TFAM | transcription factor A, mitochondrial |  |  |  |  | Yes | Yes | Yes | SE | novel | ex 2 | known |
| TFIP11 | tuftelin interacting protein 11 |  |  |  |  | Yes | No | NA | RI | novel | int 5 | known |
| TICRR | TOPBP1 interacting checkpoint and replication regulator |  |  |  |  | Yes | Yes | Yes | IE | novel | ex 4b | novel |
| TTI2 | TELO2 interacting protein 2 |  |  |  |  | Yes | Yes | NA | SE | known | ex 4 | known |
| ZNF207 | zinc finger protein 207 |  |  |  |  | Yes | Yes | Yes | SE | novel | ex 6 | known |
| ZNF248 | zinc finger protein 248 |  |  |  |  | Yes | NA | NA | SE | novel | ex 2 | known |
| ZNF317 | zinc finger protein 317 |  |  |  |  | Yes | Yes | Yes | RI | novel | int 3 | known |
| ZNF45 | zinc finger protein 45 |  |  |  |  | Yes | Yes | Yes | A3SS | novel | ex 2 | known |
| ZNF468 | zinc finger protein 468 |  |  |  |  | Yes | Yes | Yes | IE | novel | ex 2b | novel |
| ZNF528 | zinc finger protein 528 |  |  |  |  | Yes | Yes | No | SE | novel | ex 6b | novel |
| ZNF616 | zinc finger protein 616 |  |  |  |  | Yes | NA | NA | IE | novel | ex 2b | novel |
| ZNF649 | zinc finger protein 649 |  |  |  |  | Yes | Yes | Yes | A5SS | novel | ex 3 | known |
| ZNF669 | zinc finger protein 669 |  |  |  |  | Yes | Yes | Yes | SE | known | ex 3 | known |
| ZNF772 | zinc finger protein 772 |  |  |  |  | Yes | Yes | No | IE | known | ex 2 | known |
