## Supplementary Table S4 for "SRSF1 coordinates transcription and alternative splicing to modulate p53 and apoptosis"

**Curated regulatory and functional relationships used to construct the TP53-centered interaction network.**

| **Protein A** | **Protein B** | **Interaction Type** | **Key Experimental Evidence** | **STRING Score** |
| --- | --- | --- | --- | --- |
| ATR | CHEK1 | Activation [phosphorylation] | ATR phosphorylates and activates CHK1 during replication stress (54). | 0.999 |
| ATR | CHEK2 | DDR signaling | ATR/ATM phosphorylates and activates CHEK2 during replication stress (55). | 0.903 |
| BRIP1 | ATR | Replication checkpoint | BRIP1 helicase promotes replication checkpoint signaling (56). | 0.741 |
| TICRR | ATR | Replication initiation / checkpoint | TICRR regulates origin firing and checkpoint activation stimulating ATR-mediated phosphorylation of Chk1 (57) | 0.816 |
| MCM3 | TICRR | Replication initiation complex | TICRR–MTBP complex promotes assembly of the CMG helicase on the MCM2–7 helicase complex (58). | 0.954 |
| ATR | EXO1 | DNA resection regulation | ATR signaling regulates EXO1 during replication stress (59). | 0.864 |
| EXO1 | REV3L | Damage tolerance coordination | REV3L acts to counteract EXO1-dependent resection linked to translesion synthesis (60). | 0.703 |
| CHEK2 | p53 | Activation [phosphorylation] | CHEK2 Phosphorylates p53 at Ser20 and Ser15 and residues at the C terminus in response to DNA damage enhancing its stability (35). | 0.979 |
| CHEK1 | p53 | Checkpoint modulation | CHEK1 Phosphorylates p53 at Ser20 and residues at the C terminus in response to DNA damage enhancing its stability (35). | 0.812 |
| CREBBP | p53 | Activation [acetylation] | CREBBP acetylates p53 enhancing transcription (61). | 0.996 |
| MDM4 | p53 | Inhibition [binding] | MDM4 binds and inhibits p53 transcriptional activity (62). | 0.998 |
| SRSF1 | p53 | Ribosomal stress signaling | SRSF1 stabilizes p53 via RPL5-MDM2 pathway (49). | 0.78 |
| GSK3B | p53 | Activation [phosphorylation] | GSK3β phosphorylates and stabilizes p53 increasing its transcriptional activity (38). | 0.842 |
| PRKAB2 | p53 | Metabolic stress signaling | AMPK phosphorylates and stabilizes p53 during metabolic stress (40). | 0.721 |
| TAF11 | TAF2 | TFIID complex assembly | Structural interaction within TFIID complex (63). | 0.996 |
| TAF2 | NFYA | Promoter complex interaction | NF-Y interacts with basal transcription machinery ((64). | 0.882 |
| TAF2 | TP53 | Transcriptional co-regulation | TFIID supports transcription of p53 targets (41). | 0.75 |
| DICER1 | ATR | RNA-mediated DDR signaling | DICER-dependent small RNAs contribute to DNA-damage-response signaling (65). | 0.744 |
| DICER1 | p53 | p53 binding / transcriptional repression. | DICER binds unacetylated p53 and represses p53-dependent transcription through recruitment of SUV39H1 (29). | 0.788 |

The table lists experimentally supported interactions incorporated into the TP53-centered network, including interacting protein pairs, interaction type, supporting evidence, reference, and STRING confidence score. The interactions represent DNA-damage response, replication checkpoint, transcriptional regulation, metabolic and ribosomal stress, and p53-regulatory pathways relevant to SRSF1-regulated alternative-splicing events.
