## Supplementary material for "SRSF1 coordinates transcription and alternative splicing to modulate p53 and apoptosis": Source images RT-PCR

DNA ladders

Hyper ladder IV 100bp
(Thermo Fisher)

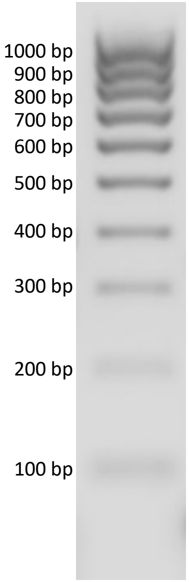


Gene ruler 50bp
(Thermo Fisher)

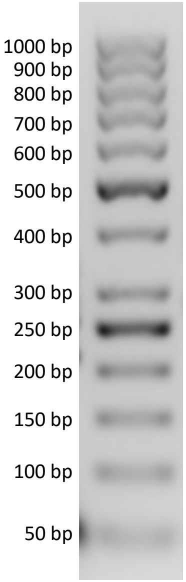


Gene ruler 100 bp+
(Thermo Fisher)

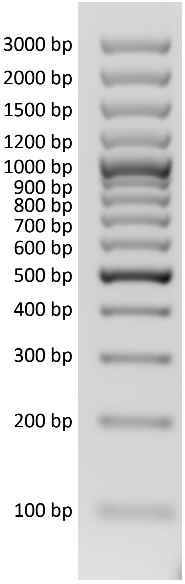

### Supplementary Figure S2 and Figure 2D source RT-PCR

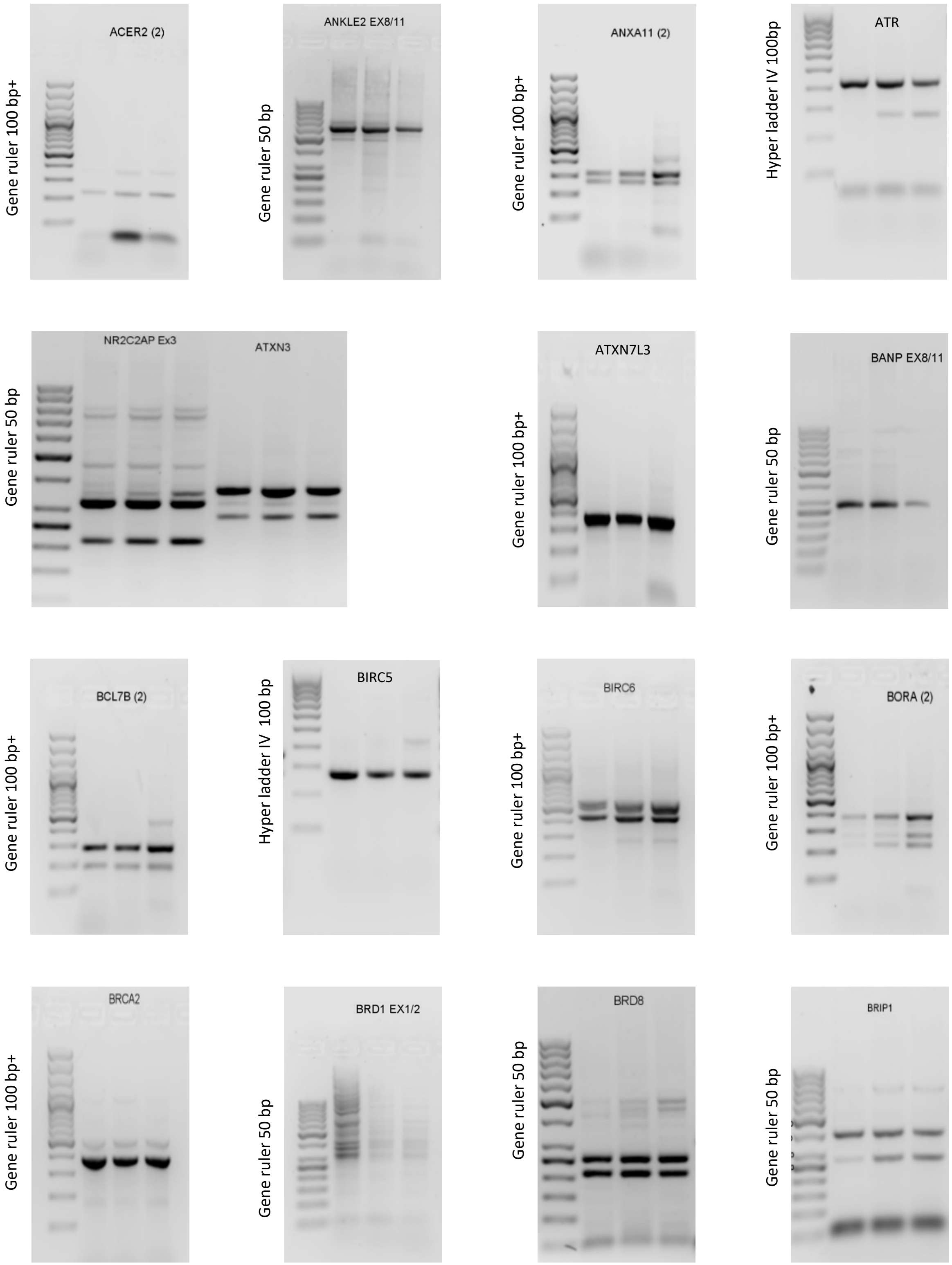

Supplementary Figure S2 and Figure 2D source RT-PCR

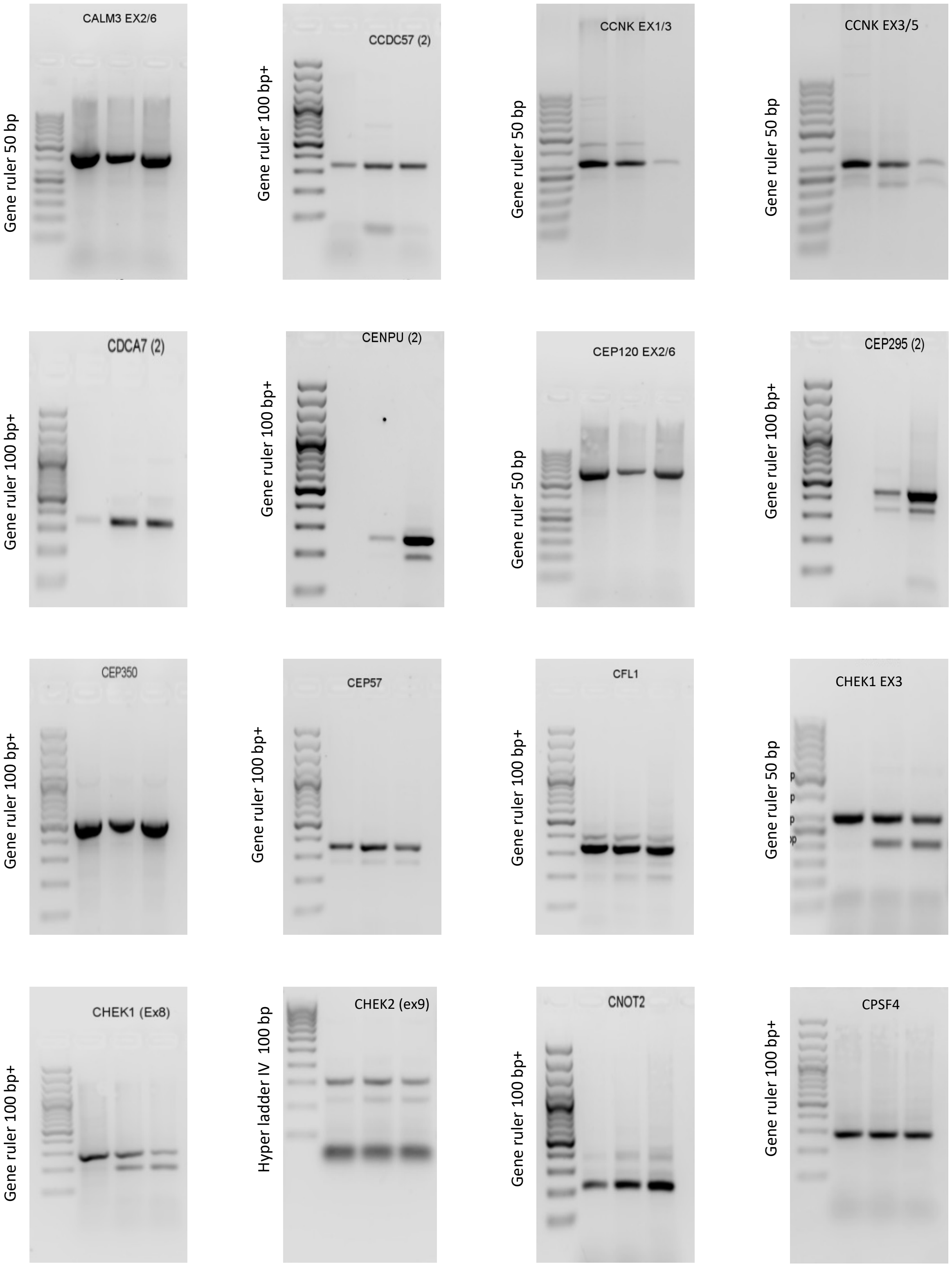

Supplementary Figure S2 and Figure 2D source RT-PCR

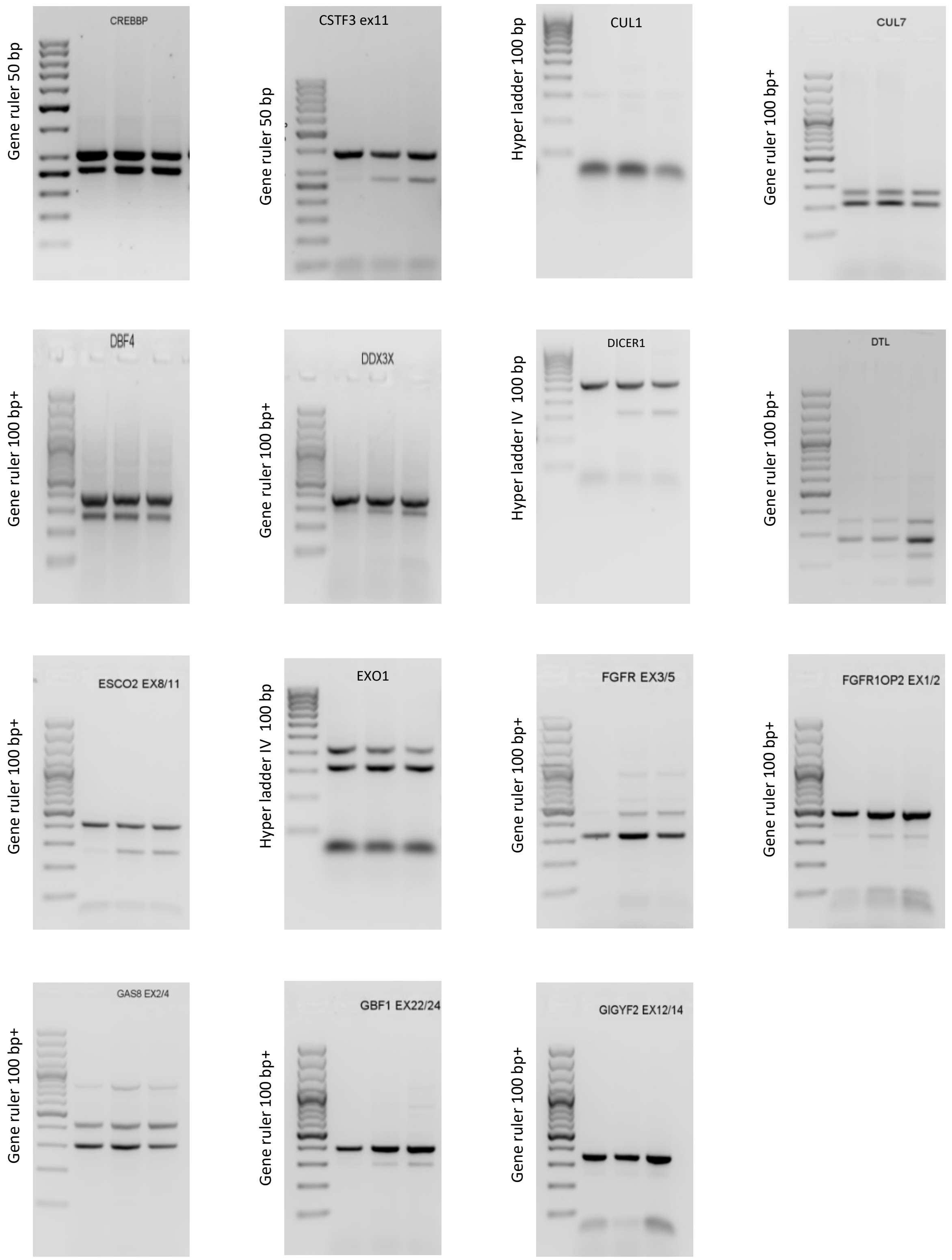

Supplementary Figure S2 and Figure 2D source RT-PCR

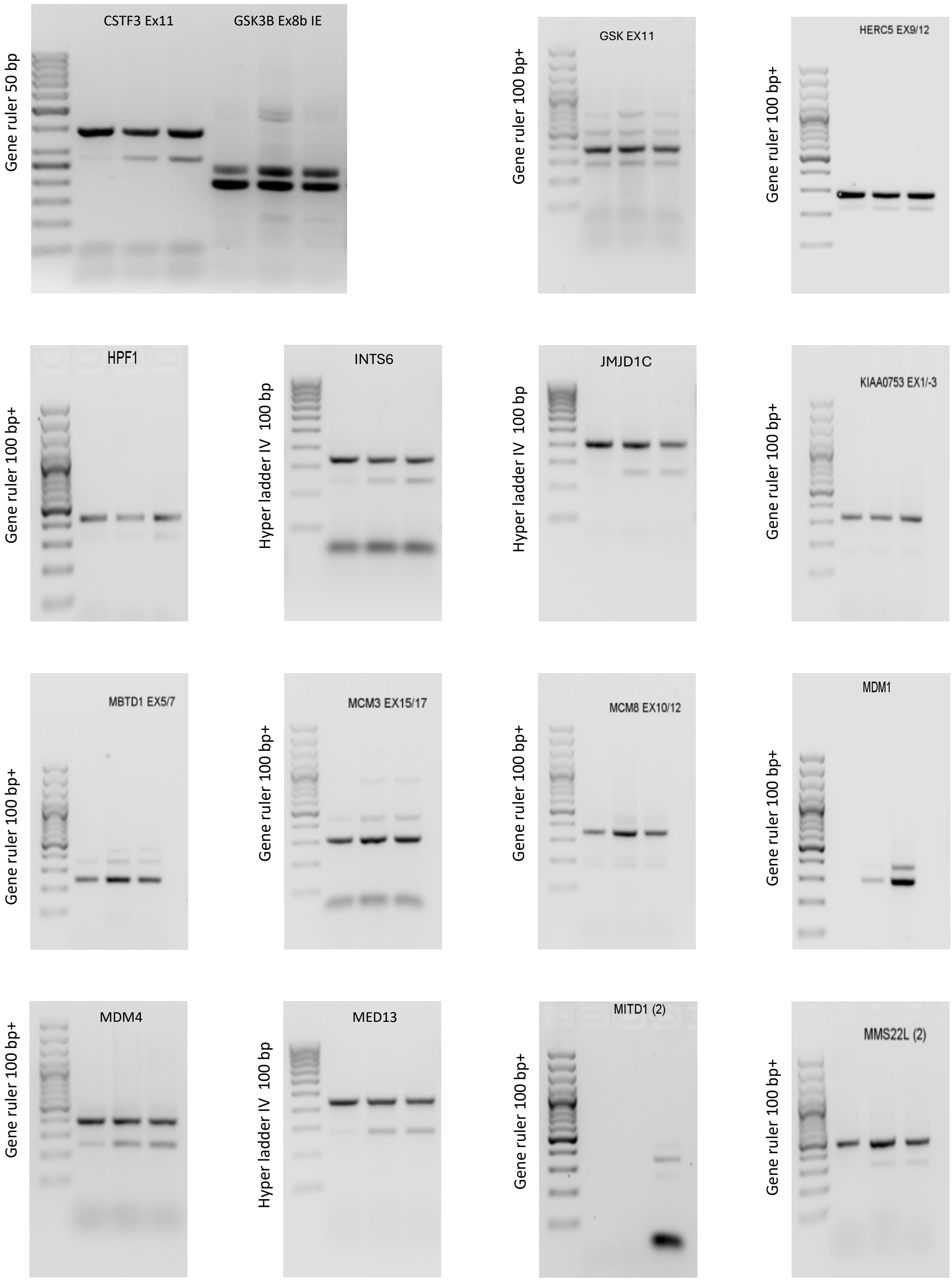

Supplementary Figure S2 and Figure 2D source RT-PCR

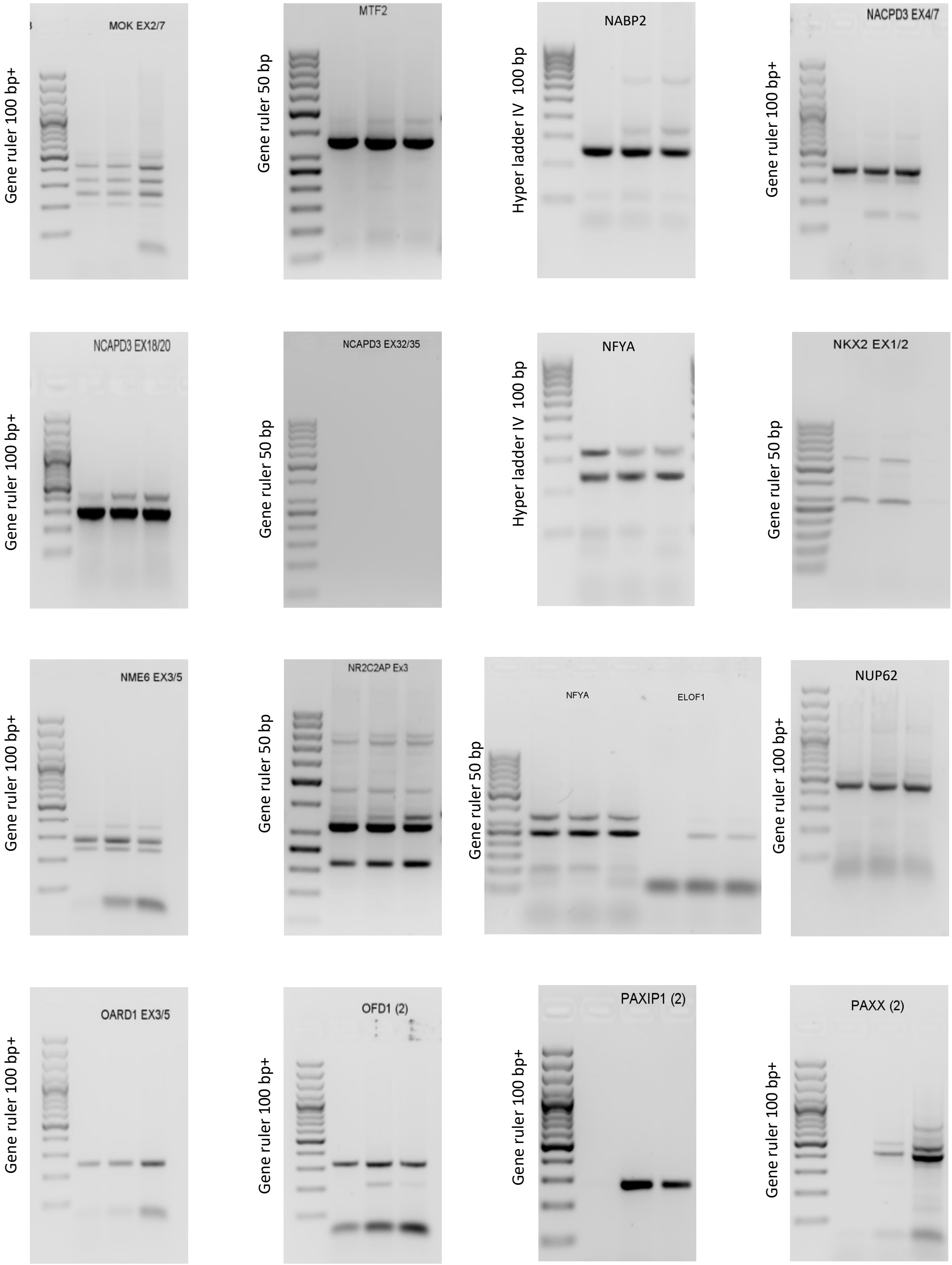

Supplementary Figure S2 and Figure 2D source RT-PCR

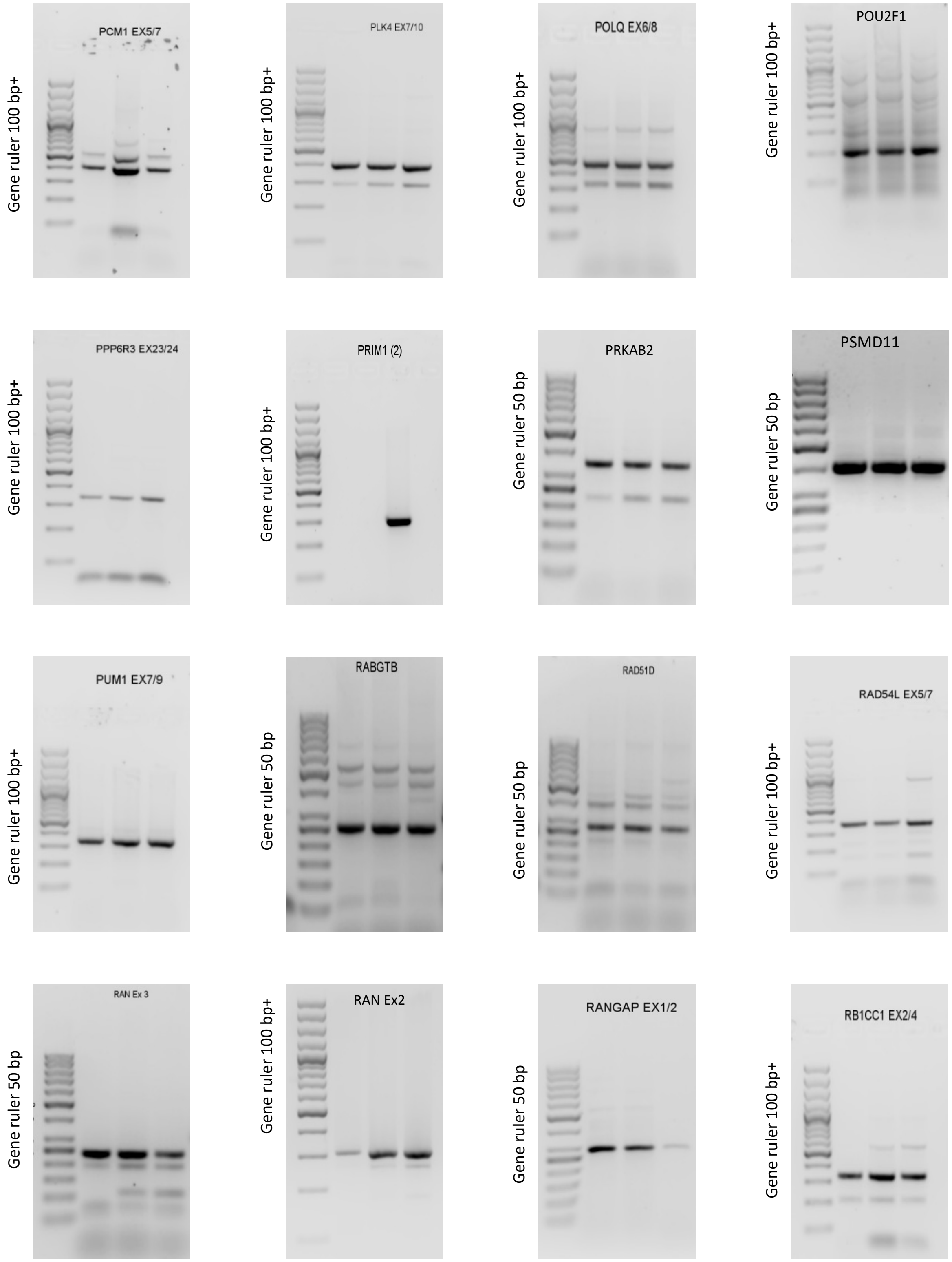

Supplementary Figure S2 and Figure 2D source RT-PCR

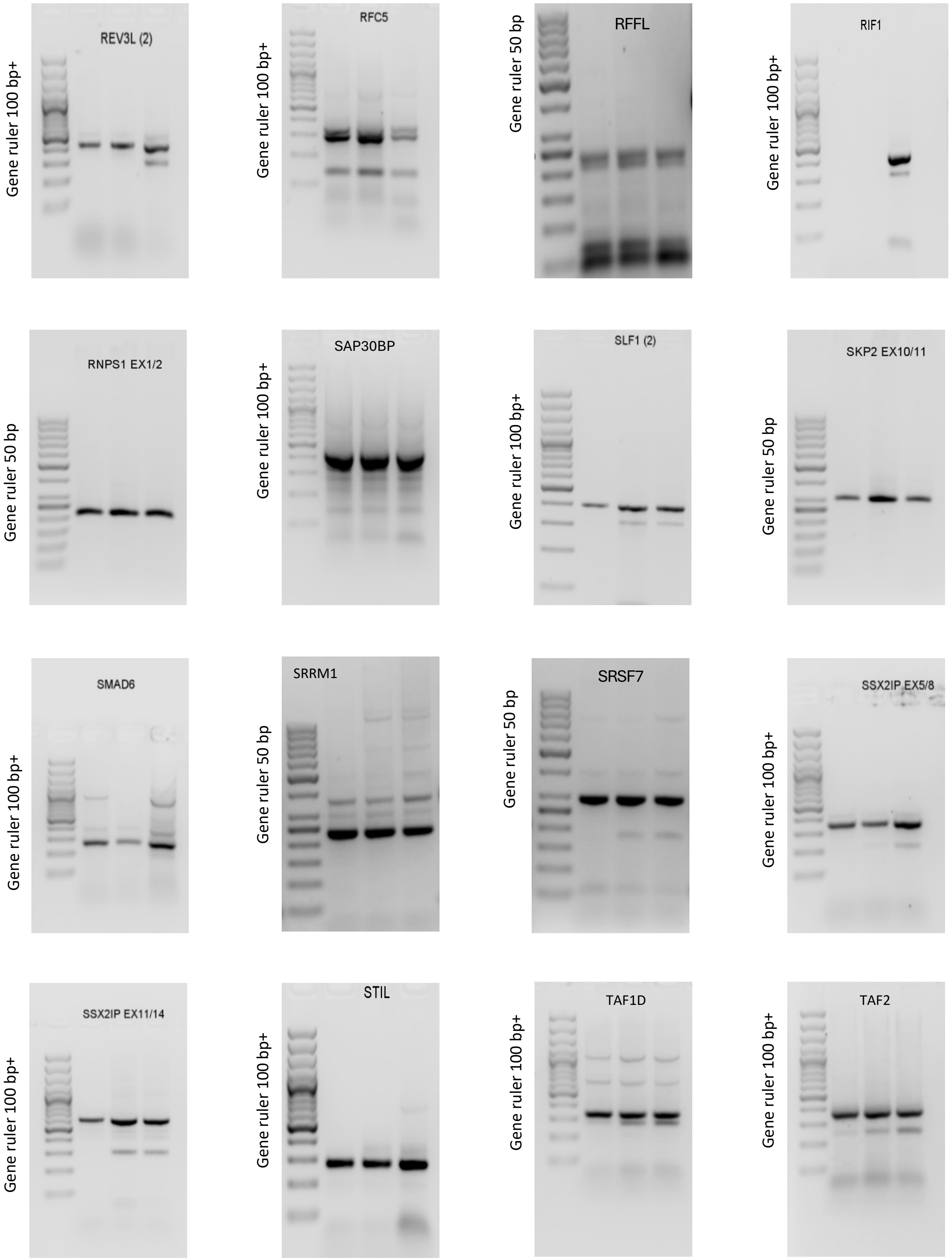

Supplementary Figure S2 and Figure 2D source RT-PCR

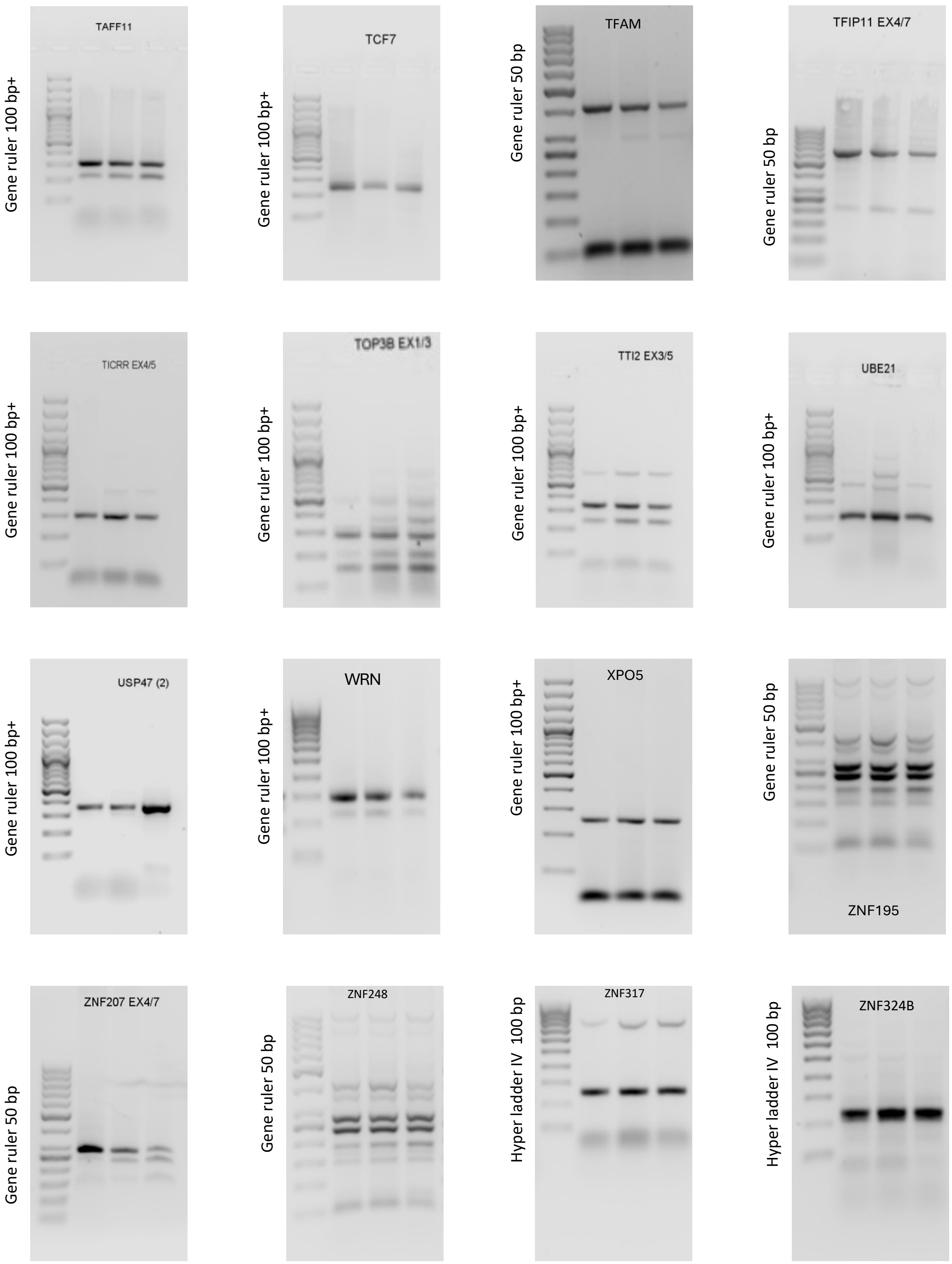

Supplementary Figure S2 and Figure 2D source RT-PCR

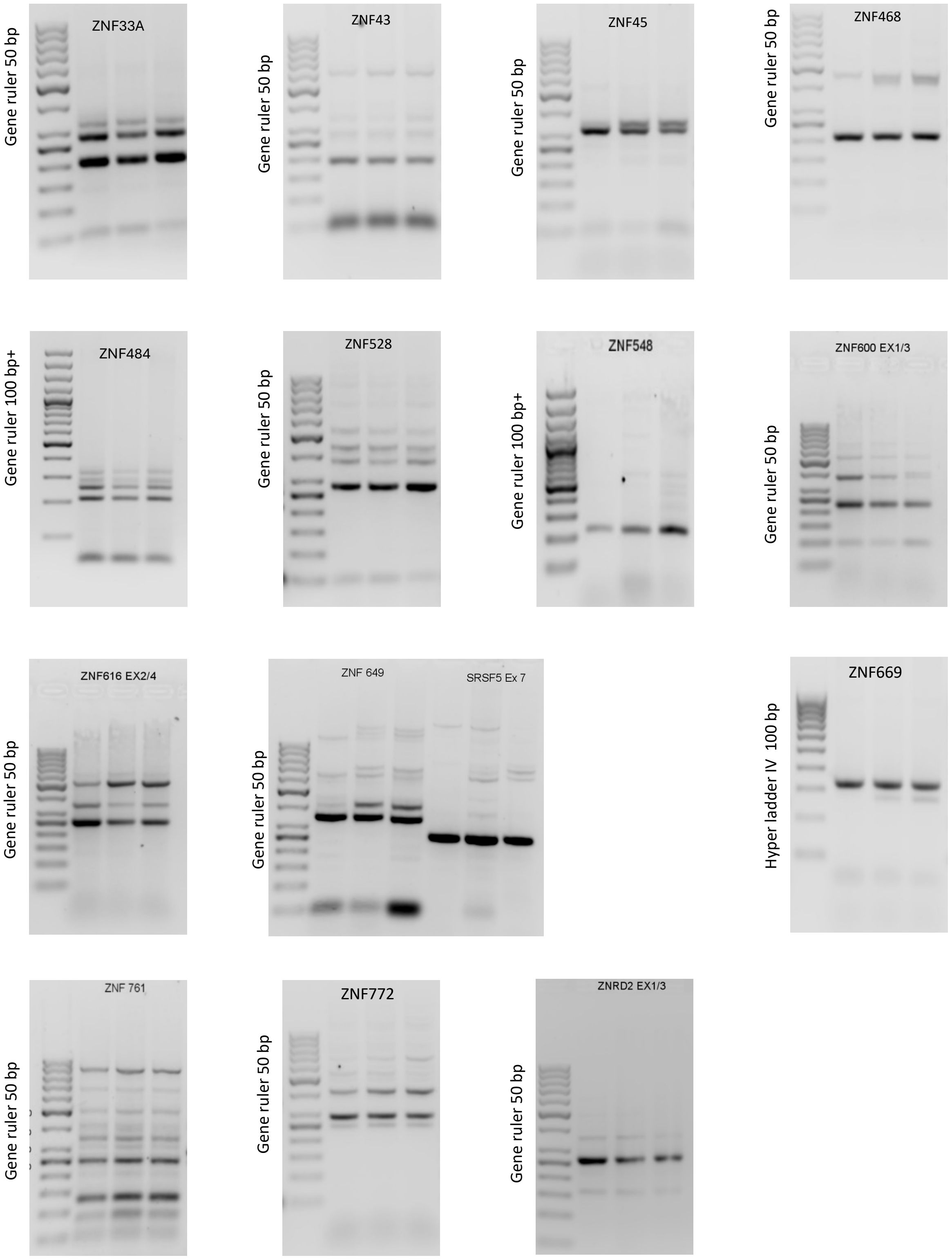

Supplementary Figure 3A, B  and Figure 2D source RT-PCR

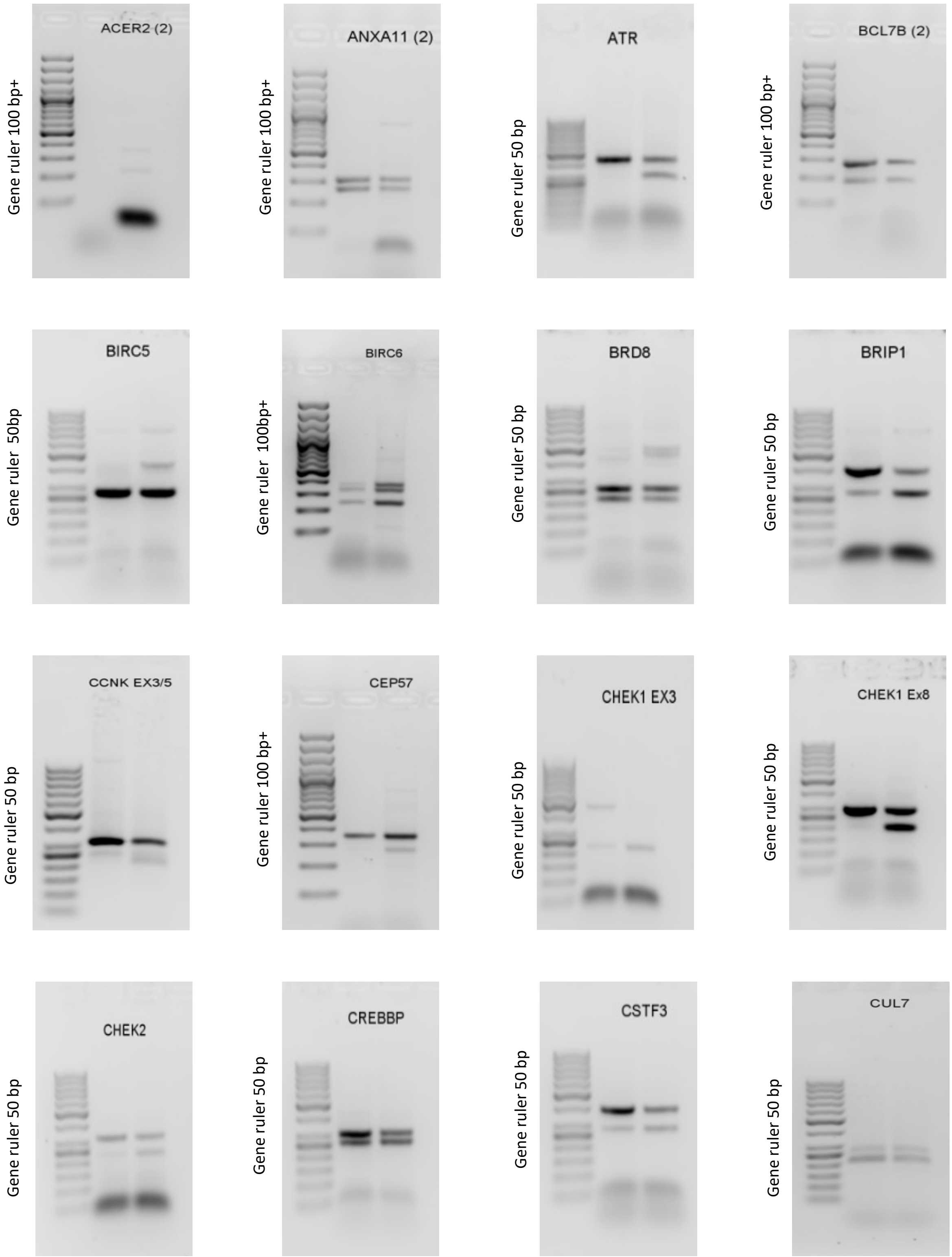

Supplementary Figure 3A, B and Figure 2D source RT-PCR

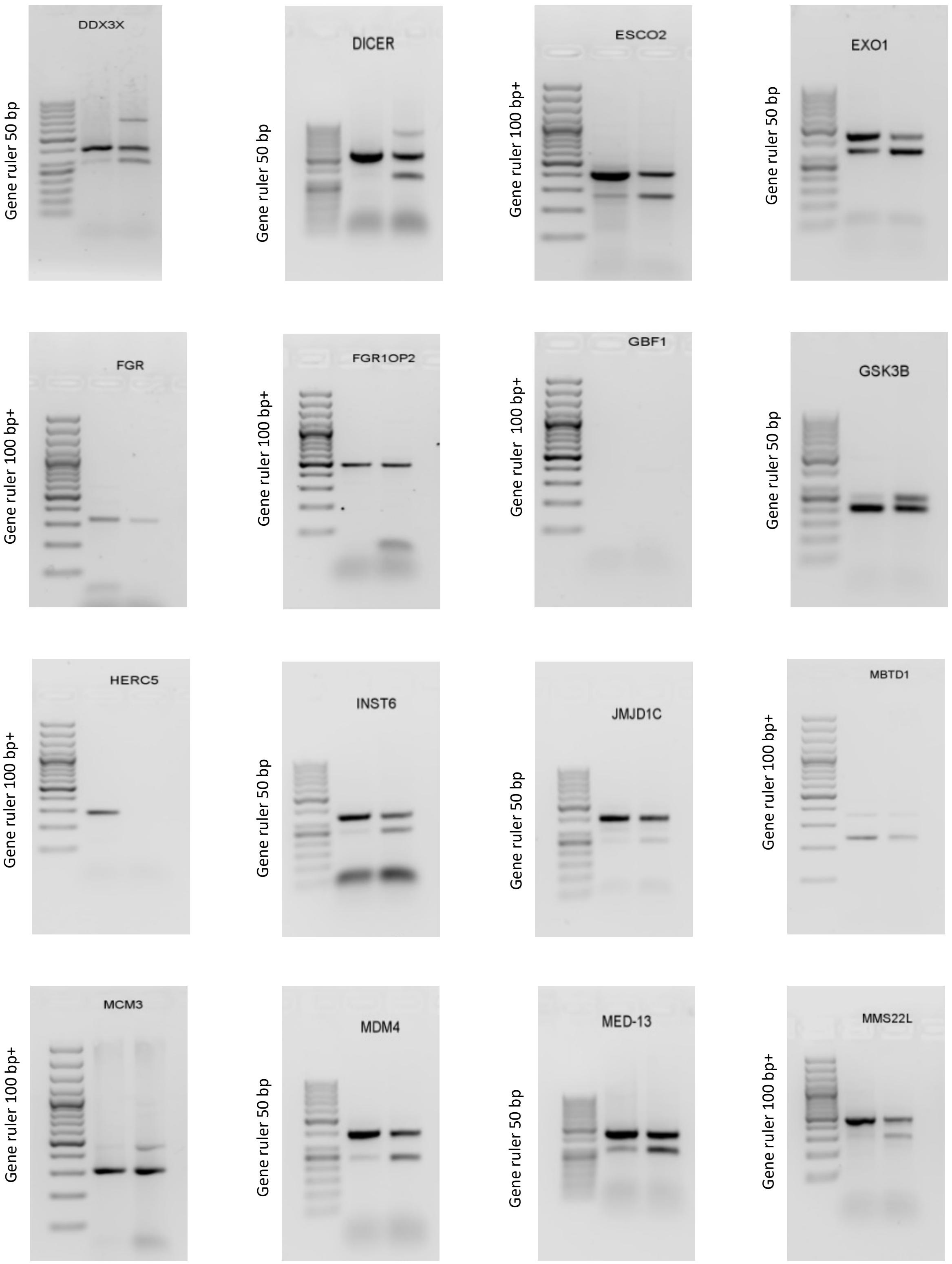

Supplementary Figure 3A, B  and Figure 2D source RT-PCR

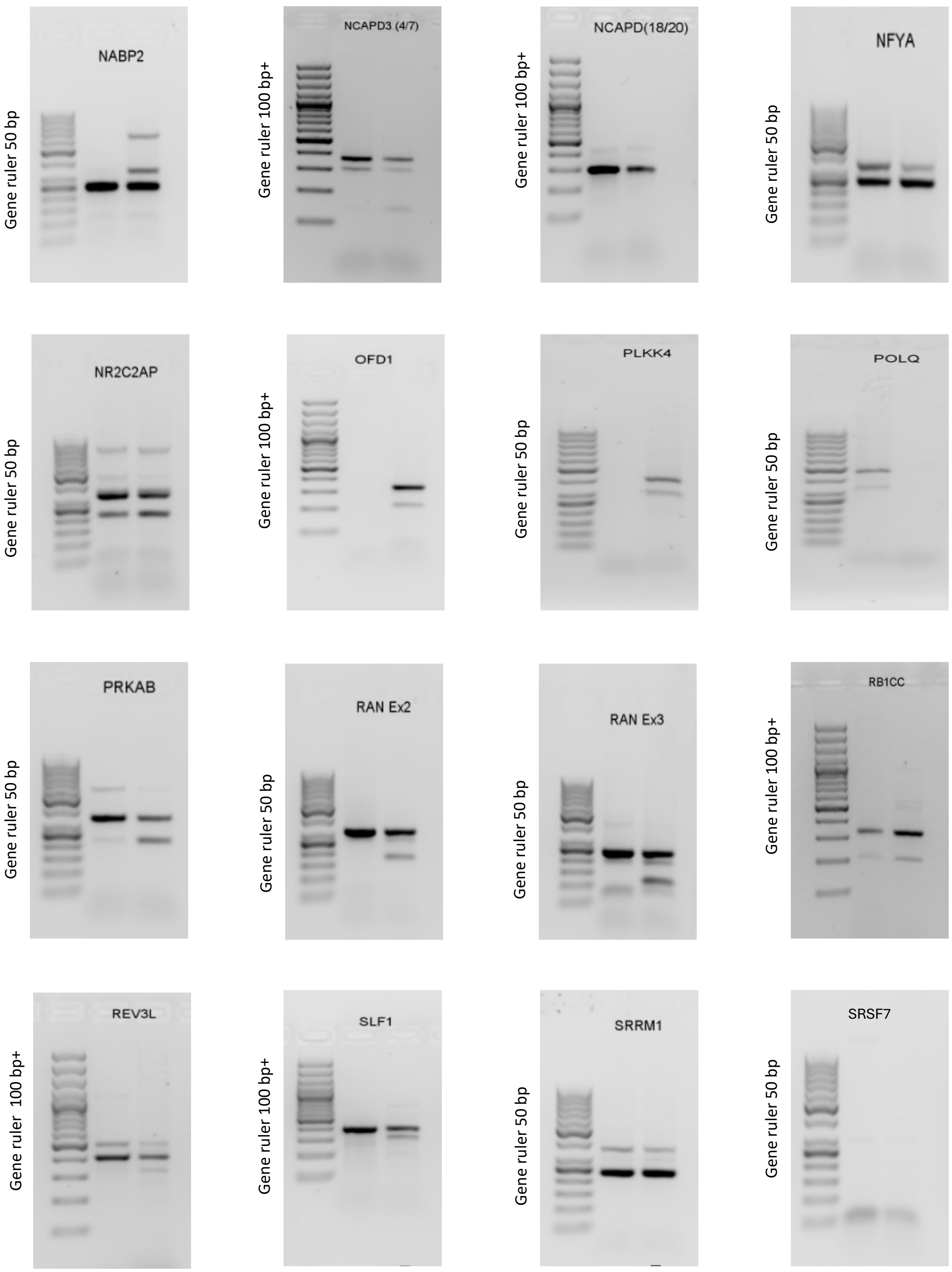

Supplementary Figure 3A, B  and Figure 2D source RT-PCR

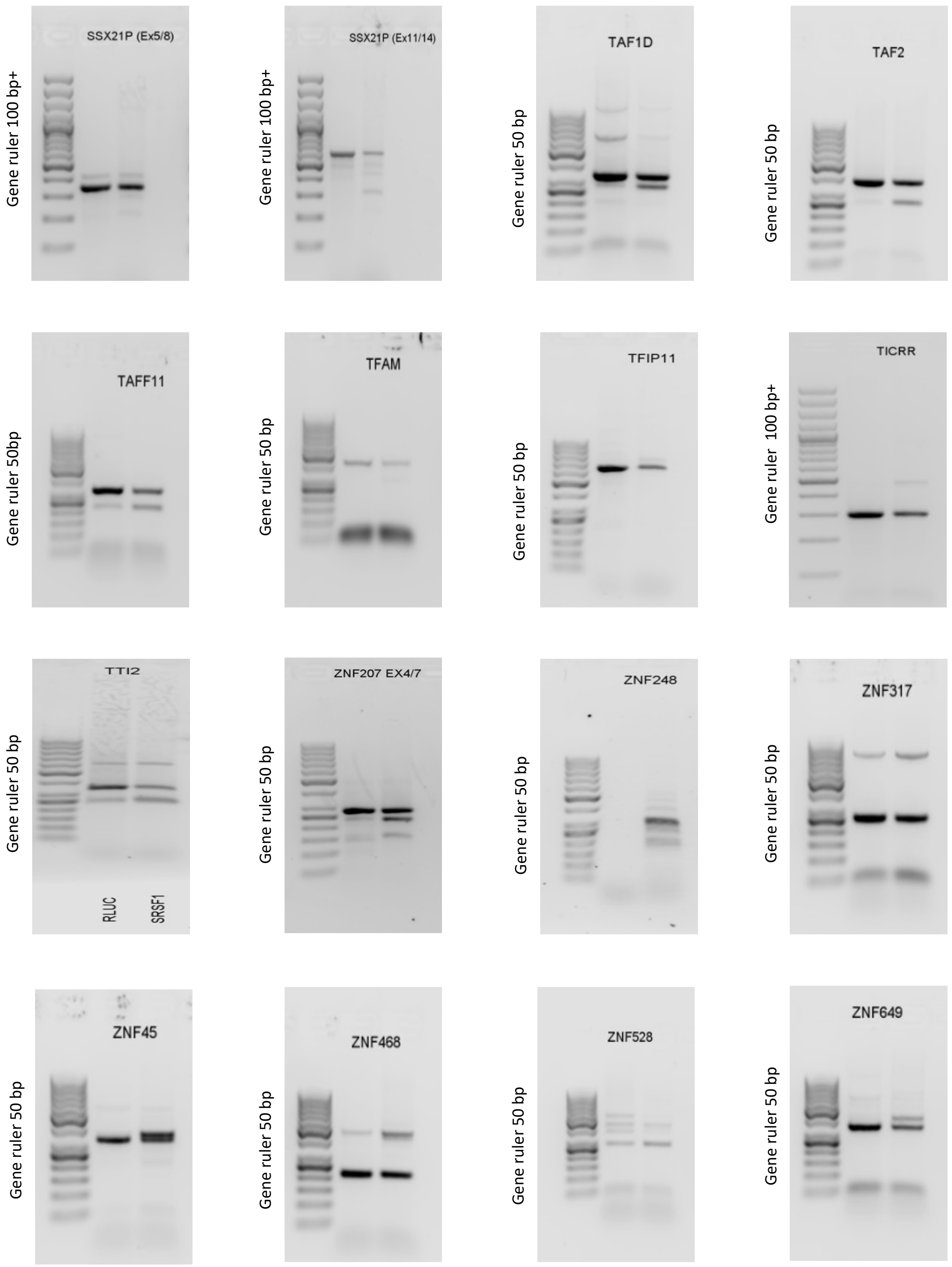

Supplementary Figure 3A, B  and Figure 2D source RT-PCR

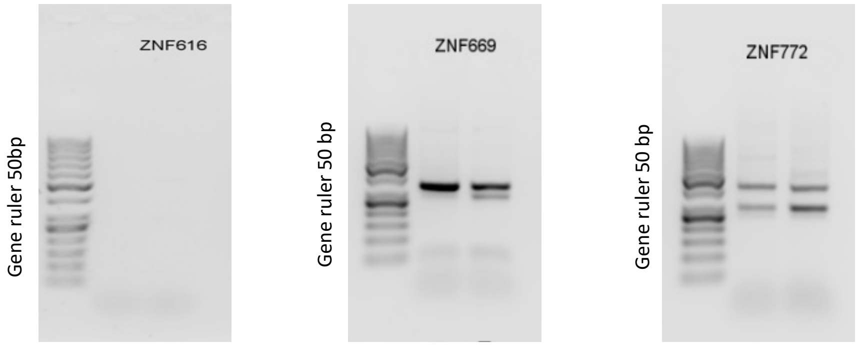

Supplementary Figure 3C, D  and Figure 2D source RT-PCR

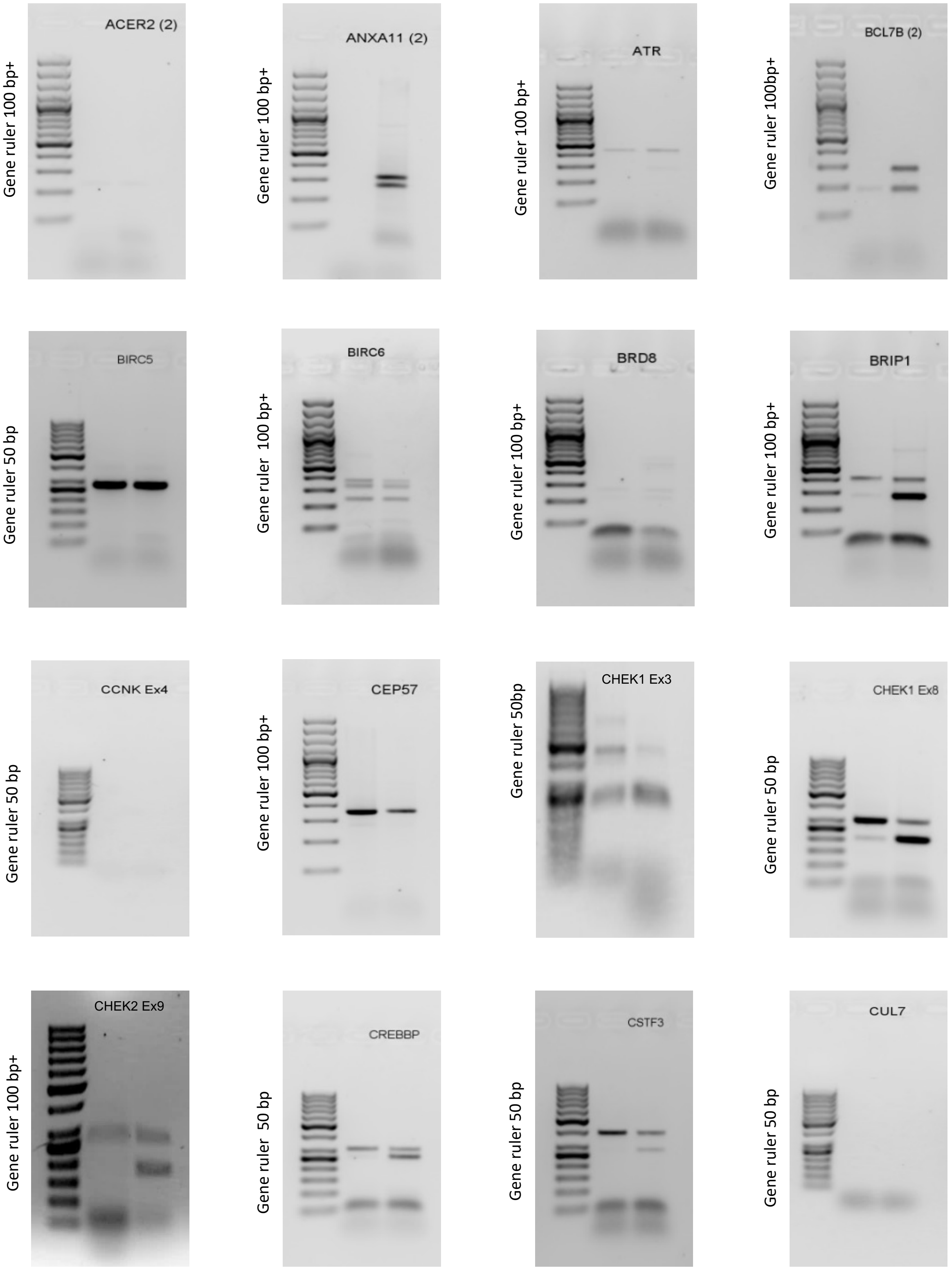

Supplementary Figure 3C, D  and Figure 2D source RT-PCR

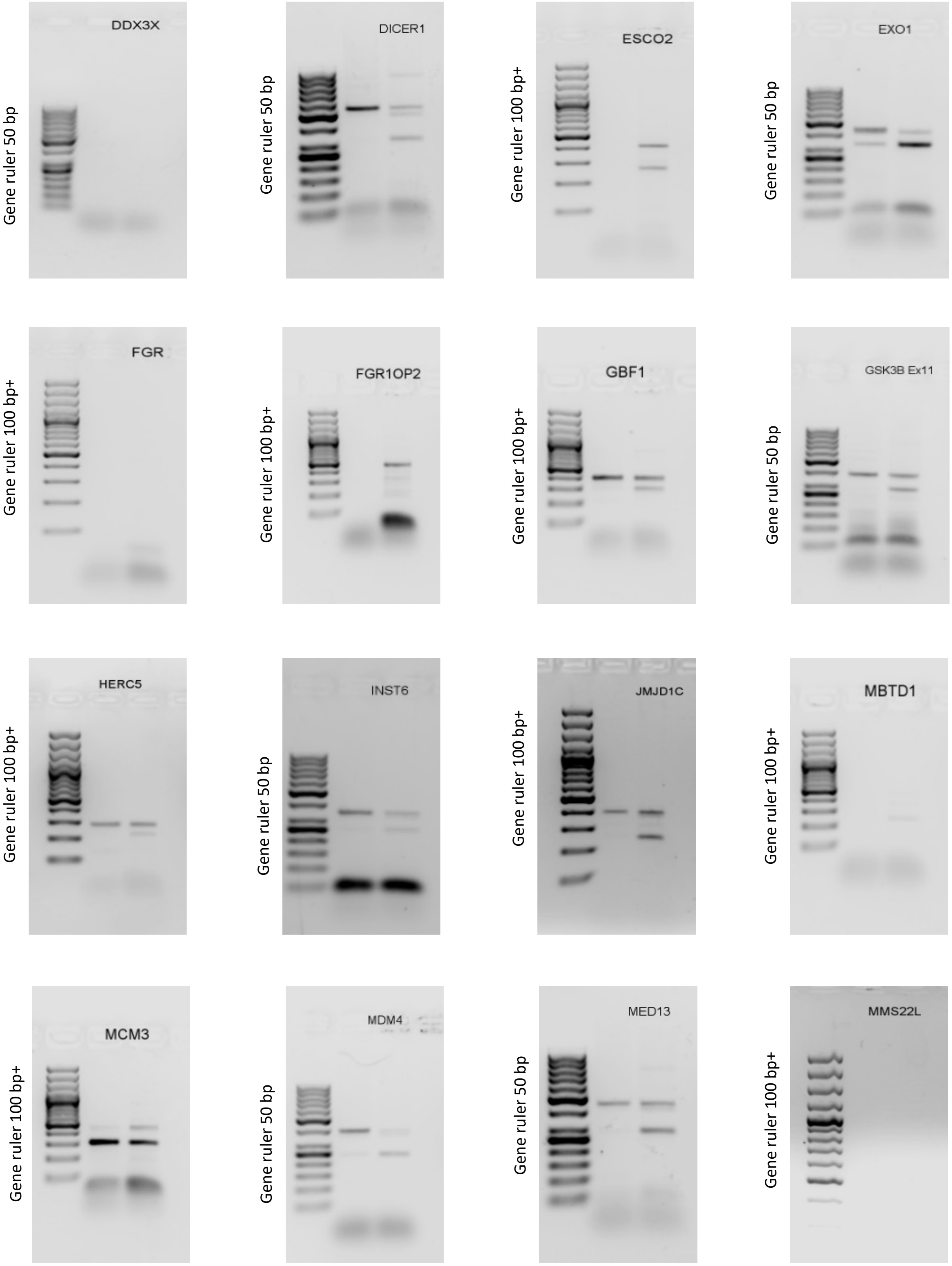

Supplementary Figure 3C, D  and Figure 2D source RT-PCR

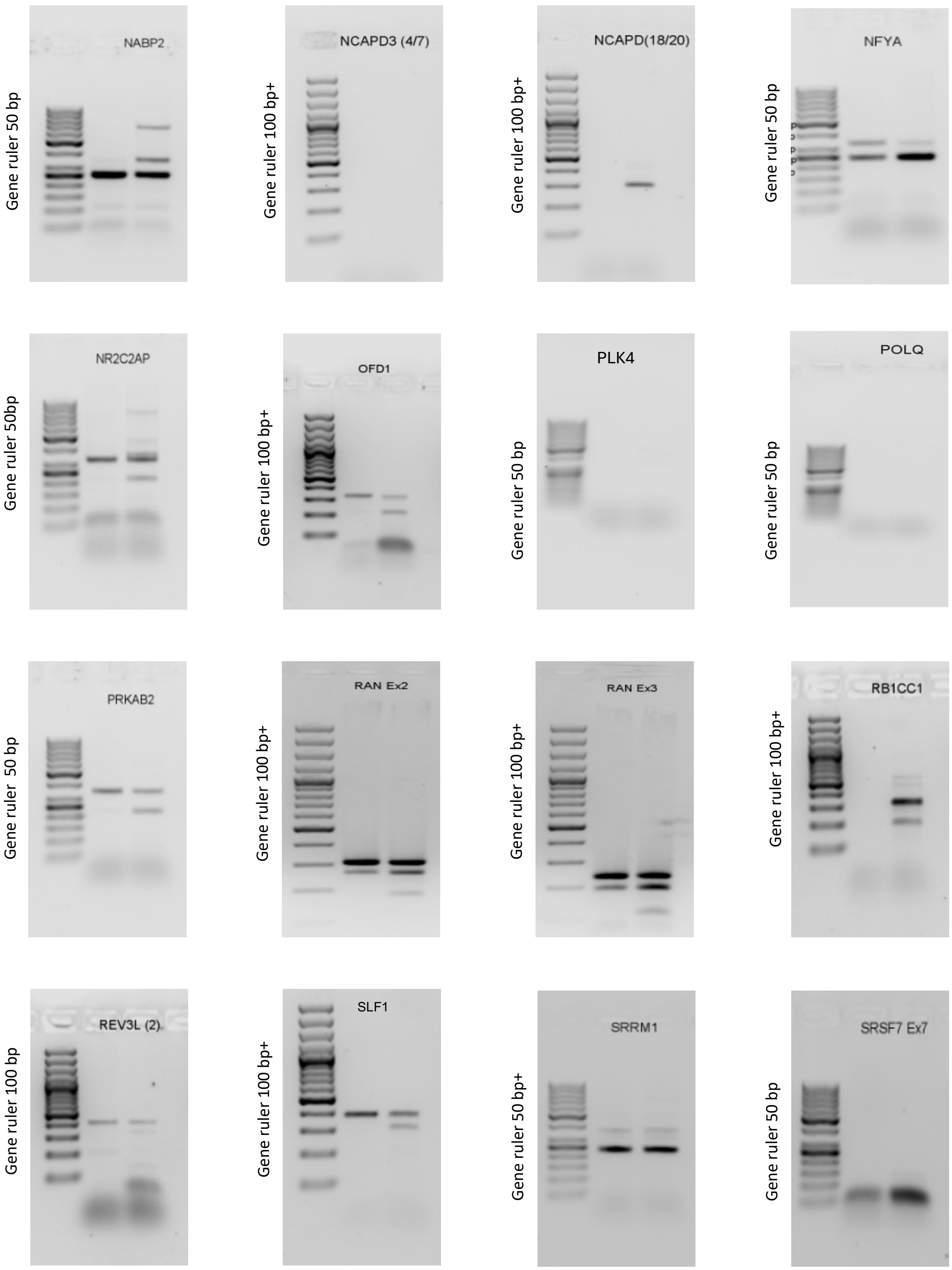

Supplementary Figure 3C, D and Figure 2D source RT-PCR

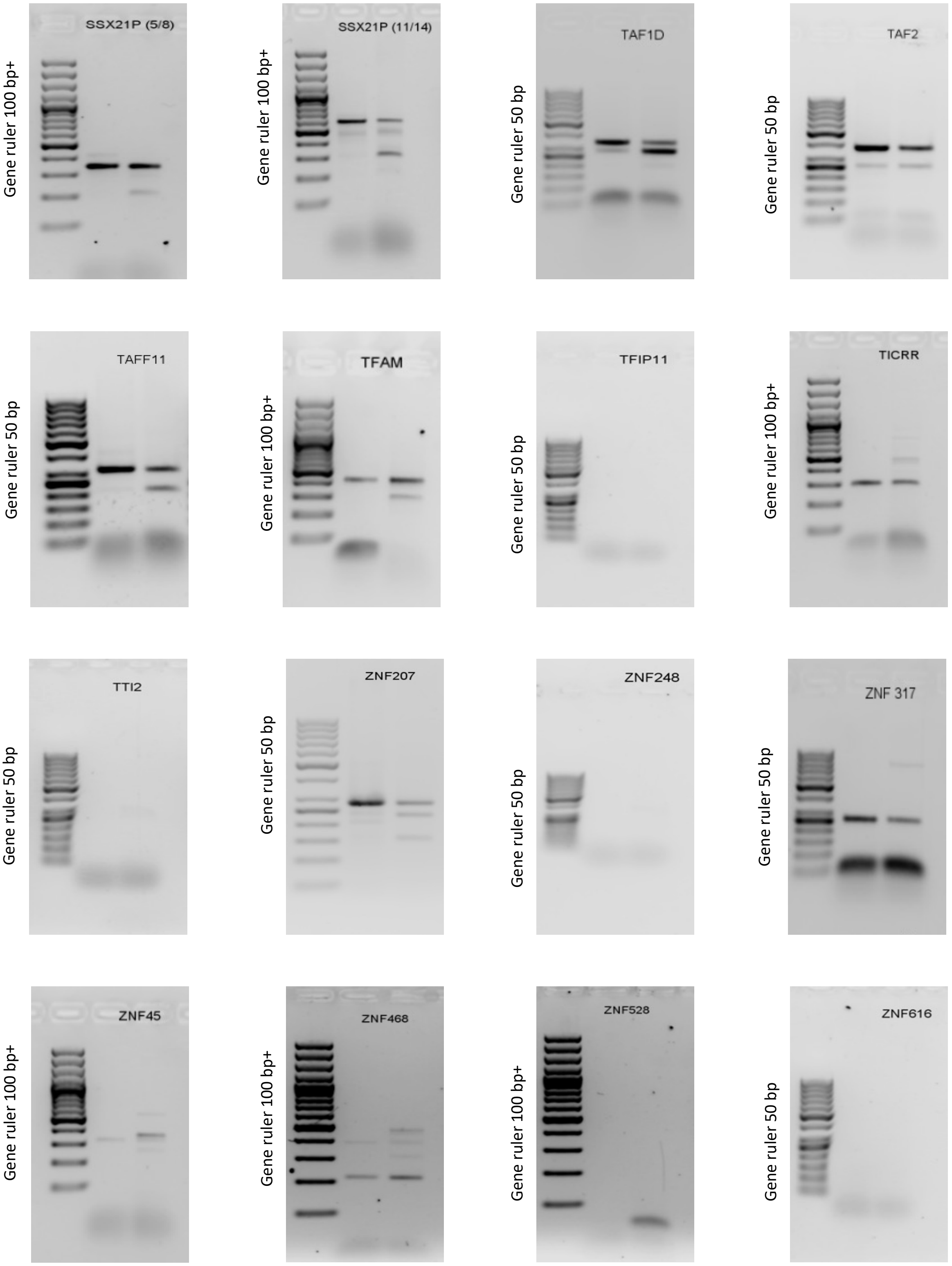

Supplementary Figure 3C, D  and Figure 2D source RT-PCR

Gene ruler 50 bp

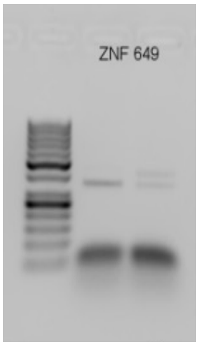


Gene ruler 50 bp

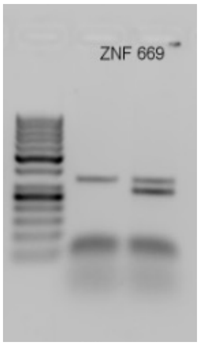


Gene ruler 100 bp+

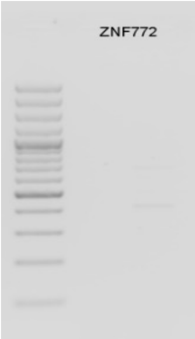
