## Supplementary figures and images for "SRSF1 coordinates transcription and alternative splicing to modulate p53 and apoptosis"

### Source images Western Blot

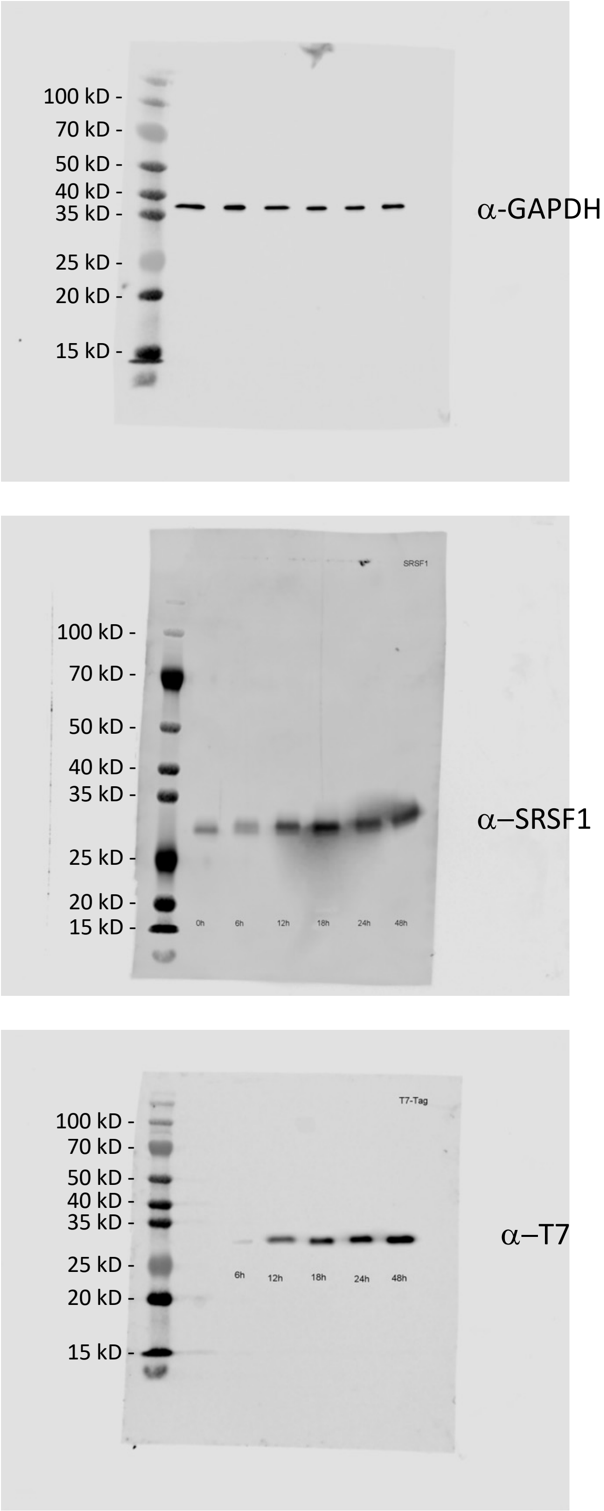

Figure 3B

Doxorubicin

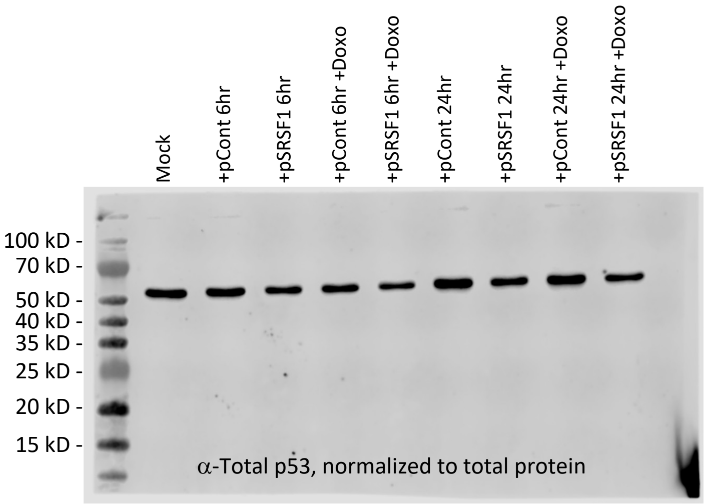


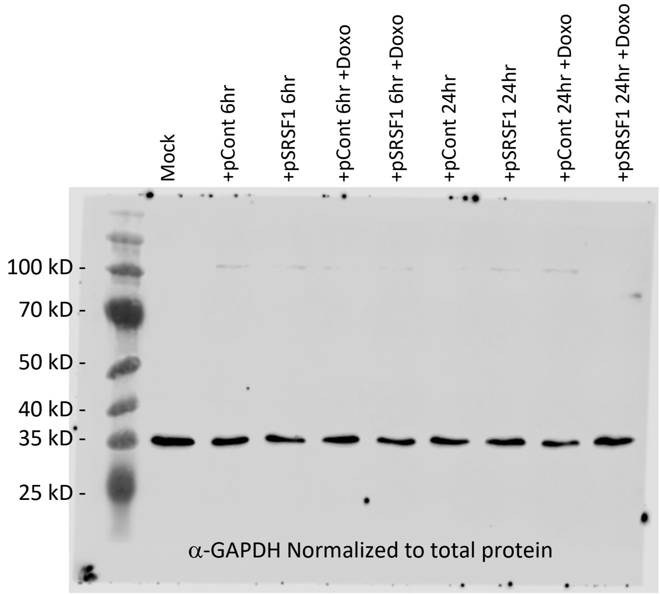


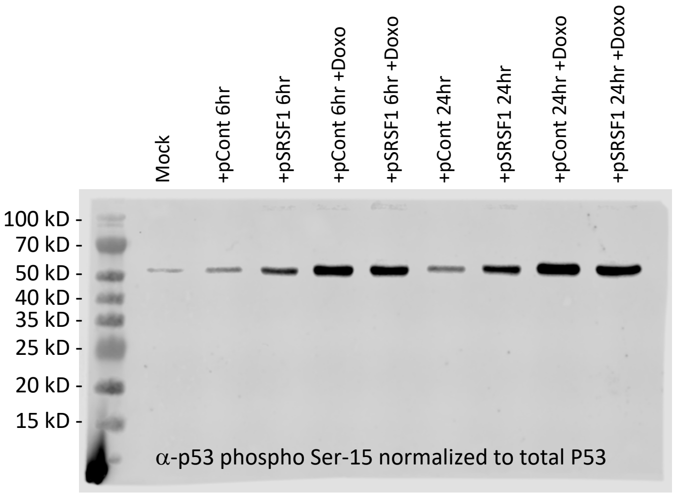


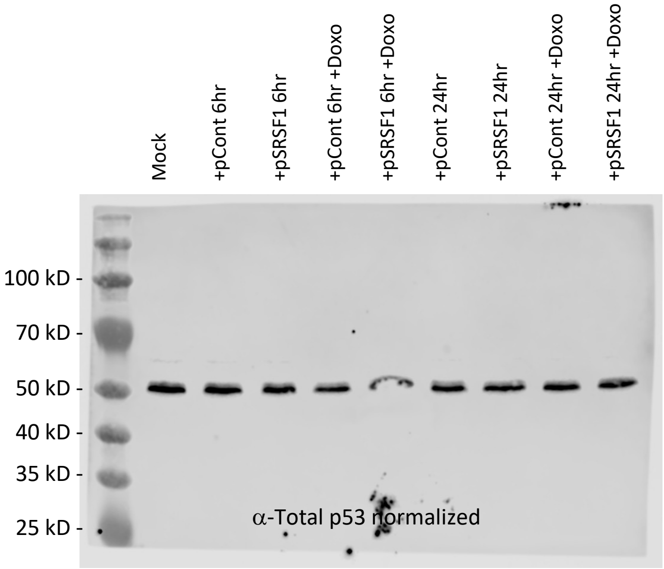

Figure 3B

Cisplatin

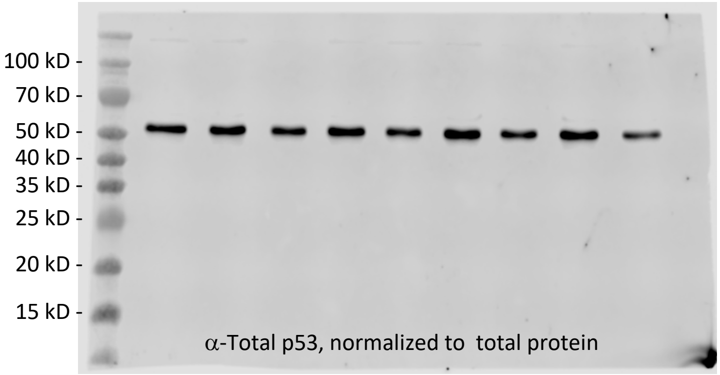


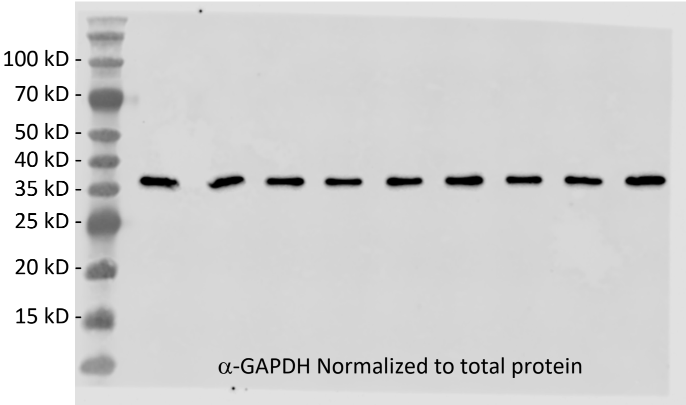


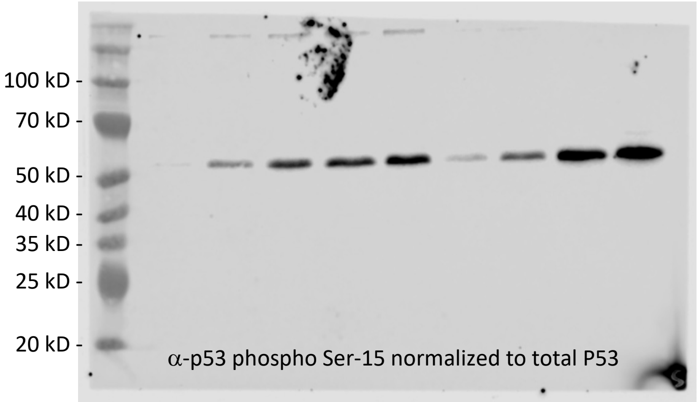


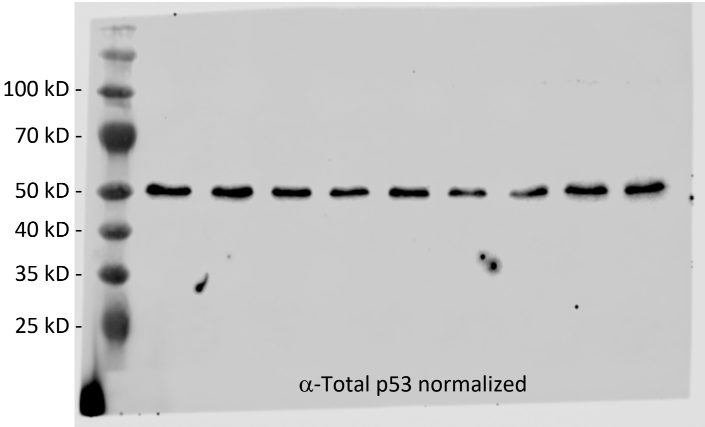

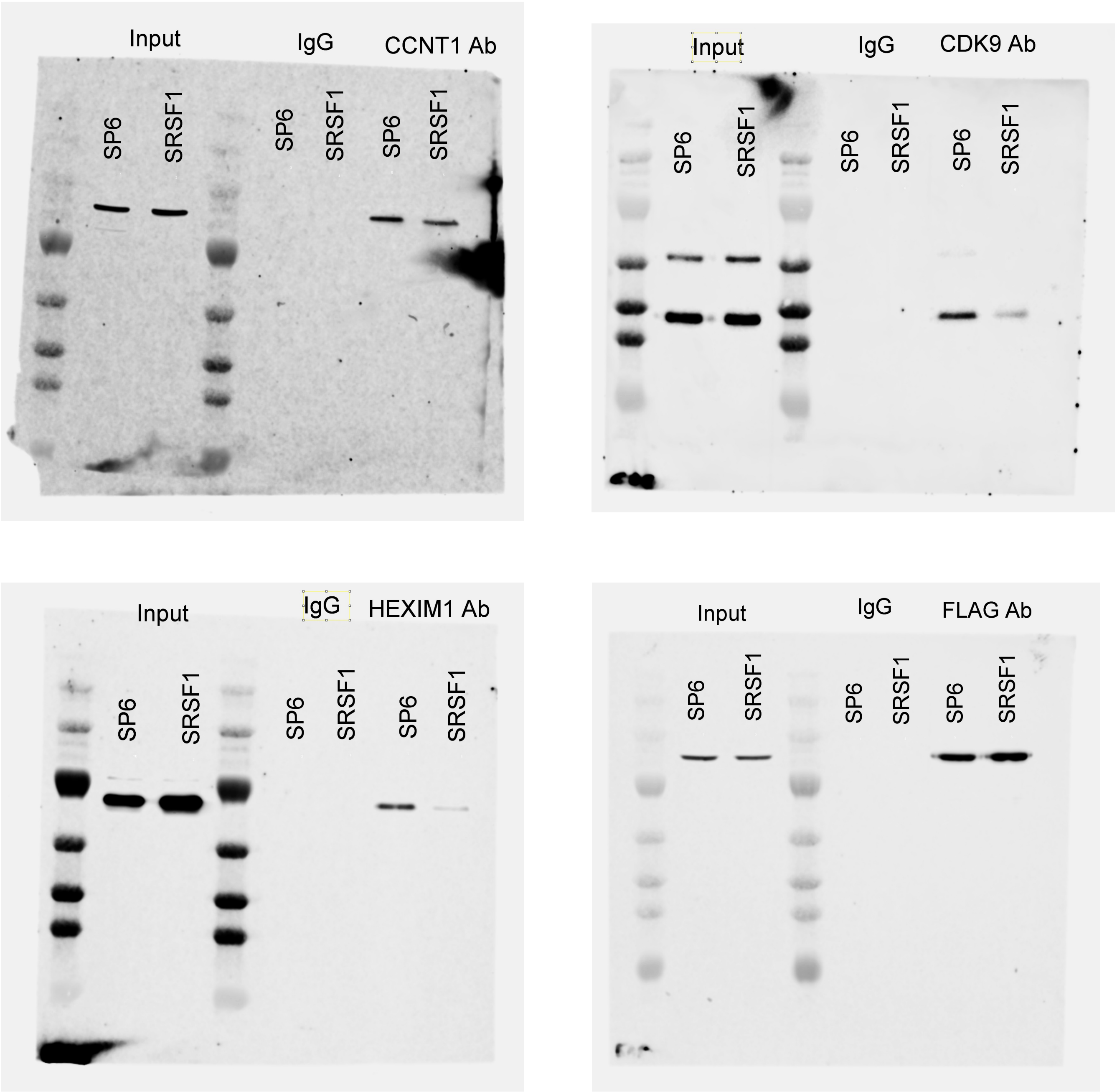
